# Metabolic engineering of yeast for *de novo* production of kratom monoterpene indole alkaloids

**DOI:** 10.1101/2024.05.22.595370

**Authors:** Maxence Holtz, Daniela Rago, Ida Nedermark, Frederik G. Hansson, Beata J. Lehka, Lea G. Hansen, Nils E. J. Marcussen, Wouter J. Veneman, Linda Ahonen, Juraithip Wungsintaweekul, Ron P. Dirks, Carlos G. Acevedo-Rocha, Jie Zhang, Jay D. Keasling, Michael K. Jensen

**Author notes:** To whom correspondence should be addressed. Michael K. Jensen, and Jay D. Keasling.

## Abstract

Monoterpene indole alkaloids (MIAs) from *Mitragyna speciosa* (“kratom”), such as mitragynine and speciogynine, are promising novel scaffolds for opioid receptor ligands for treatment of pain, addiction, and depression. While kratom leaves have been used for centuries in South-East Asia as stimulant and pain management substance, the biosynthetic pathway of these psychoactives have only recently been partially elucidated. Here, we demonstrate the *de novo* production of mitragynine and speciogynine in *Saccharomyces cerevisiae* through the reconstruction of a five-step synthetic pathway from common MIA precursor strictosidine comprising fungal tryptamine 4-monooxygenase to bypass an unknown kratom hydroxylase. Upon optimizing cultivation conditions, a titer of ∼290 µg/L kratom MIAs from glucose was achieved. Untargeted metabolomics analysis of lead production strains led to the identification of numerous shunt products derived from the activity of strictosidine synthase (STR) and dihydrocorynantheine synthase (DCS), highlighting them as candidates for enzyme engineering to further improve kratom MIAs production in yeast. Finally, by feeding fluorinated tryptamine and expressing a human tailoring enzyme, we further demonstrate production of fluorinated and hydroxylated mitragynine derivatives with potential applications in drug discovery campaigns. Altogether, this study introduces a yeast cell factory platform for the biomanufacturing of complex natural and new-to-nature kratom MIAs derivatives with therapeutic potential.

## Introduction

Addiction to opioids, including morphine and synthetic derivatives is among the leading causes of death in several age categories in the US with >80,000 deaths in 2021 (National Institute on Drug Abuse, 2023). In addition to their addiction risk, use of opioids is subject to tolerance build-up towards their analgesic effect as well as adverse effects, including nausea, dizziness, constipation and respiratory depression (Chakraborty & Majumdar, 2021; Queremel Milani & Davis, 2023). While effective for the treatment of pain, the abuse and adverse effects cause a huge unmet medical need to discover and develop safer alternatives for the treatment of chronic pain.

*Mitragyna speciosa* (“kratom”) is a tropical tree from Southeast Asia that has been used for centuries by local farmers to combat pain, fatigue and increase productivity (Cinosi et al., 2015). Kratom has since become popular in Western countries for the (self)treatment of pain, depression, and to alleviate opioid withdrawal symptoms (Grundmann, 2017). It is currently estimated that kratom is used by millions in the USA alone (Swogger et al., 2022). Although kratom is currently a controlled substance in many countries, several recent preclinical and clinical studies show a promising therapeutic potential and safety profile in humans for kratom alkaloids motivating further preclinical and clinical research (Prevete et al., 2022).

The bioactivities of kratom are attributed to a mixture of > 50 monoterpene indole alkaloids (MIAs) of which the most abundant are mitragynine (MG), paynantheine, speciogynine (SG), speciociliatine and 7-OH-mitragynine (7-OH-MG). MG and 7-OH-MG are both partial agonists of the human µ-opioid receptor (MOR) which is the main receptor targeted for pharmacotherapy of pain (Ehrlich et al., 2019). Contrarily to classical opioids, MG and 7-OH-MG exhibit a biased signaling at MOR being selective only for the G protein signaling pathway, while side-effects of classical opioids are believed to be attributed to the activation of the β-arrestin pathway (Kruegel et al., 2016). In addition to its potential for pain treatment, MG is also a κ-opioid receptor (KOR) antagonist, making it a potential dual-action antidepressant drug (Kruegel et al., 2016). Other kratom alkaloids such as SG, which does not affect MOR signaling, could be promising leads for Alcohol Use Disorder (AUD) treatment by modulating the δ-opioid receptor (DOR) (Gutridge et al., 2021). To summarize, the wide array of bioactivities of kratom alkaloids on the different opioid receptors makes them interesting scaffolds to develop new treatments for pain management, addiction, and depression.

Supplying MIAs at scale for medicinal use through extraction or total chemical synthesis is challenged by their high structural complexity and general low availability in plant tissues (Courdavault et al., 2021; Zhang et al., 2022). In addition, these considerations hinder the generation of new to nature derivative libraries in sufficient quantities for drug discovery campaigns. Refactoring MIA biosynthesis in yeast cell factories has been proposed as an alternative supply source for these high-value compounds, promising to be more reliable, sustainable, and scalable by fermentation of relatively cheap feedstocks (Etit et al., 2024). Indeed, others and we have recently reported the engineering of yeast for production of anticancer catharanthine and vindoline (D. Gao et al., 2022; J. Gao et al., 2023; Zhang et al., 2022), anti-psychotic alstonine and various other yohimbines and heteroyohimbines (Bradley et al., 2023, 2024; Dror et al., 2024; T. Liu et al., 2022). Besides manufacturing natural MIAs, yeast cell factories can be used as platforms to generate MIA derivatives that would be challenging to produce chemically, either by feeding analogs of pathway precursors or by expressing tailoring enzymes (Bradley et al., 2020, 2023, 2024).

Recent work established that MG and SG (its C20 epimer) are produced in *M. speciosa* in five enzymatic steps from the common MIA precursor strictosidine (**Fig. 1, “Canonical pathway”**) (K. Kim et al., 2023; Schotte et al., 2023). Strictosidine first undergoes de-glycosylation (SGD, strictosidine-O-β-D-glucosidase), followed by action of a medium chain alcohol dehydrogenase (DCS, dihydrocorynantheine synthase), enol O-methyltransferase (OMT, EnolMT), and finally oxidation and methylation at C9 position. The stereoselectivity of the different DCS variants is responsible for the production of either mitragynine (20S) or speciogynine (20R). While the last two enzymes responsible for the formation of the methoxy group in C9 in *M. speciosa* are still unknown, Kim et al., (2023) reported an enzyme named *Hp*9OMT from firebush *Hamelia patens* capable of methylating 9-OH-corynantheidine to form MG and SG, and further reported the refactoring of a 6-enzyme bioconversion of supplemented MIA precursors tryptamine and secologanin to MG and SG at µg/L-scale in yeast (Kim et al., 2023). Over the course of this study, the core kratom pathway, comprising SGD, *Ms*DCS1 and EnolMT, was refactored in a strictosidine-producer yeast strain leading to 0.9 mg/L *de novo* corynantheidine production (Dror et al., 2024). The introduction of the methoxy group at the C9 position, which is essential for the bioactivity of MG/SG at opioid receptors (Kruegel & Grundmann, 2018), was however not reported.

**Figure 1.**
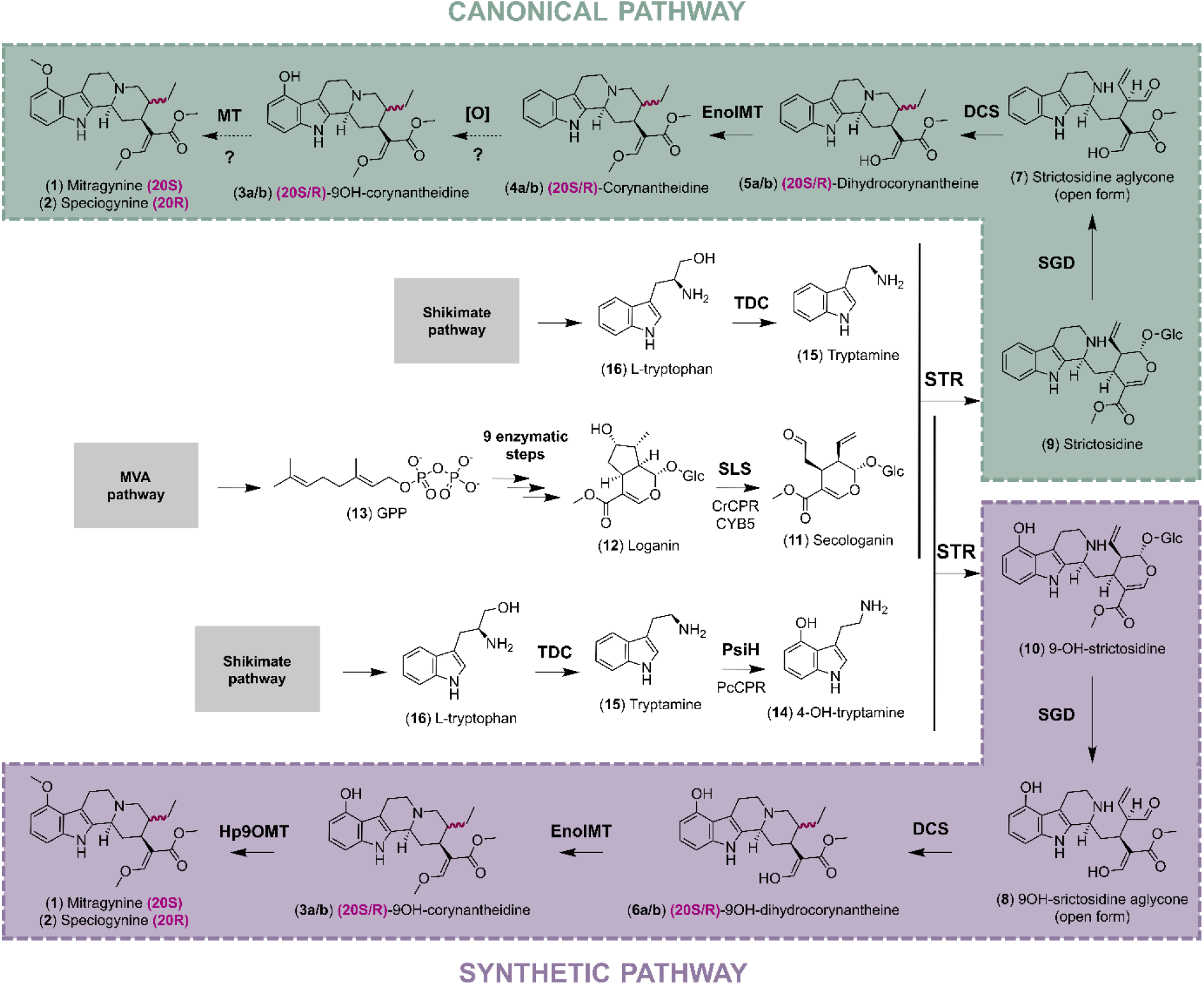
Biosynthetic pathways for production of mitragynine and speciogynine in yeast. **A)** Canonical pathway from kratom plant *Mitragyna speciosa* (Schotte et al., 2023). **B)** Synthetic biosynthetic pathway engineered in yeast based on fungal tryptamine 4-monooxygenase *Pc*PsiH circumventing an unknown enzyme in the native kratom plant biosynthetic pathway. Abbreviations not already defined in the main text are as follows: GPP, geraniol pyrophosphate; CPR, NADPH-cytochrome P450 reductase; CYB5, cytochrome b5; SLS, secologanin synthase; TDC, tryptophan decarboxylase; STR, strictosidine synthase. Dashed arrows indicate that the corresponding enzyme is unknown. Multiple arrows indicate more than one enzymatic step. The information for all genes is also listed in **Supp. Table 1**.

In this paper, we aimed to capitalize on a previously engineered parental strictosidine-producing yeast strain to explore the possibility of *de novo* production of MG and SG in yeast, as well as the opportunity to use this platform for the biosynthesis of new-to-nature kratom alkaloids.

## Results and Discussion

### Synthetic MG pathway refactoring in yeast

To refactor *de novo* kratom alkaloid biosynthesis in yeast, a strictosidine platform strain, MIA-CZ-1, was further optimized by deleting *ROX1* and overexpressing *INO2* and *ZWF1* in previously reported parental strain MIA-CM-5 (Zhang et al., 2022). These modifications were made to increase the activity of multiple cytochrome P450s that limit flux through the secologanin biosynthetic pathway (Bradley et al., 2023; Zhang et al., 2022). Specifically, ROX1 is a transcriptional repressor of the heme biosynthetic gene *HEM13* in *S. cerevisiae*, and its inactivation has been shown to improve cytochrome P450 (CYP) activity in yeast cell-factories (Bradley et al., 2023; Q. Liu et al., 2021). Secondly, overexpression of ZWF1, catalyzing the first committed step of the pentose phosphate pathway, was introduced to promote NADPH generation used as cofactor by P450s. Finally, INO2 is a transcription factor involved in lipid biosynthesis regulation, and its overexpression has been shown to cause endoplasmic reticulum expansion, also supporting P450 activity (J.-E. Kim et al., 2019). MIA-CZ-1 produced 25.5 mg/L of strictosidine in 96 deepwell (DW96) plates and 90.9 mg/L in Ambr 250-mL fed-batch microbioreactors, representing a 39% improvement over MIA-CM-5 (**Supp. Fig. 1**).

While the first enzymes downstream of strictosidine aglycone (**7**) in the MG (**1**) / SG (**2**) pathway (referred to as “kratom MIAs” in this paper) were recently discovered (Schotte et al., 2023), the last two enzymes responsible for hydroxylation and O-methylation of the corynantheidine (**4a/b**) scaffold at position C9 are still unknown (**Fig. 1, “Canonical pathway”**). To circumvent the missing C9 hydroxylase in yeast, co-expression of the fungal tryptamine 4-monooxygenase *Pc*PsiH from *Psilocybe cubensis* involved in production of the psychedelic psylocibin (together with the enzymes downstream strictosidine) was recently successfully tested, enabling production of MG and SG in yeast upon supplementation of secologanin and tryptamine (Kim et al., 2023). In this synthetic kratom pathway, the hydroxyl group in C9 position of the MG scaffold is introduced earlier through production of 4-OH-tryptamine (**14**). The production of 9-OH-corynantheidine (**3a/b**) is then dependent on the promiscuity of the different pathway enzymes to accept hydroxylated versions of their natural substrate (**Fig. 1, “Synthetic pathway”**). This approach was also recently reported in a *N. benthamiana* leaf transient expression platform, but 9-OH-corynantheidine (**3a/b**) was not observed in tobacco leaves although 9-OH-strictosidine was detected (Schotte et al., 2023). To validate the substrate promiscuity observed by Kim *et al*. (2023) and demonstrate *de novo* production of 9-OH-corynantheidine (**3a/b**) in yeast, we expressed the fungal tryptamine 4-monooxygenase *Pc*PsiH from *P. cubensis* together with its reductase *Pc*CPR in MIA-CZ-1, generating MIA-KM-1. Here, we employed a CYP/CPR design previously engineered in yeast to produce psylocibin (Milne et al., 2020). MIA-KM-1 was cultivated for 144 h in optimized medium for small-scale MIA production (3×SC supplemented with 3mM tryptophan and 10g/L soy peptone) with glucose as carbon source (Zhang et al., 2022). The supernatant was subsequently analyzed by Liquid Chromatography-High Resolution Tandem Mass Spectrometry (LC-MS/MS) to assess the presence of MIAs. Production of (**14**) was validated by comparing *m/z*, retention time, and fragmentation pattern with an authentic 4-OH-tryptamine standard (**Supp. Fig. 2A**). We also detected a peak matching the mass of 9-OH-strictosidine (**10**) ([M+H]^+^ m/z=547.2286). An MS/MS spectrum extracted from this peak showed a mass shift relative to the strictosidine standard consistent with a hydroxyl substitution in the indole moiety of the molecule (**Supp. Fig. 2B**).

As it seemed that *Cr*STR was able to accept (**14**) as a substrate, we next integrated the three downstream enzymes SGD, DCS and *Ms*EnolMT into the genome of MIA-KM-1. We used *Rs*SGD from *Rauwolfia serpentina*, which was the most active homolog tested in yeast previously (Zhang et al., 2022). Three variations of this core kratom pathway were assembled, one for each of the DCS homologs described by Schotte *et al*. (2023): *Ms*DCS1 and *Ms*DCS2 from *M. speciosa* (MIA-KM-2 and MIA-KM-3, respectively) and *Cp*DCS from *C. pubescens* (MIA-KM-4) (**Supp. Table 3**). Supernatant analysis from these different strains showed apparition of two peaks (RT = 6.47 min and 6.57 min) matching the mass of 9-OH-corynantheidine (**3a/b**) ([M+H]^+^ m/z=385.2122) (**Fig. 2A**). While standards for (**3a/b**) are not commercially available, the MS/MS spectrum extracted from these two peaks showed a mass shift relative to a 20S-corynantheidine standard that again points to a hydroxyl substitution in the molecule’s indole moiety (**Fig. 2BC**). These strains also produced both 20S- and 20R-corynantheidine (**4a/b**) ([M+H]^+^ m/z=369.2172) as well as putative 20S- and 20R-dihydrocorynantheine (**5a/b**) ([M+H]^+^ m/z=355.2016) and 20S- and 20R-9OH-dihydrocorynantheine (**6a/b**) ([M+H]^+^ m/z=371.1965) peaks that all were absent from the parental strain MIA-KM-1 (**Supp. Fig. 3**). Altogether this led us to confirm the *de novo* production of **3a/b** in yeast. Depending on the DCS variant expressed, different product stereochemistries were obtained in accordance with previous work (K. Kim et al., 2023; Schotte et al., 2023). *Ms*DCS1 mainly produced corynanthe MIAs with 20S stereochemistry in yeast with minor 20R product. Conversely, *Ms*DCS2 mostly produces 20R stereochemistry and an additional unidentified peak with m/z=385.2122 (RT=7.09 min), while *Cp*DCS only produced 20R corynanthe MIAs.

**Figure 2.**
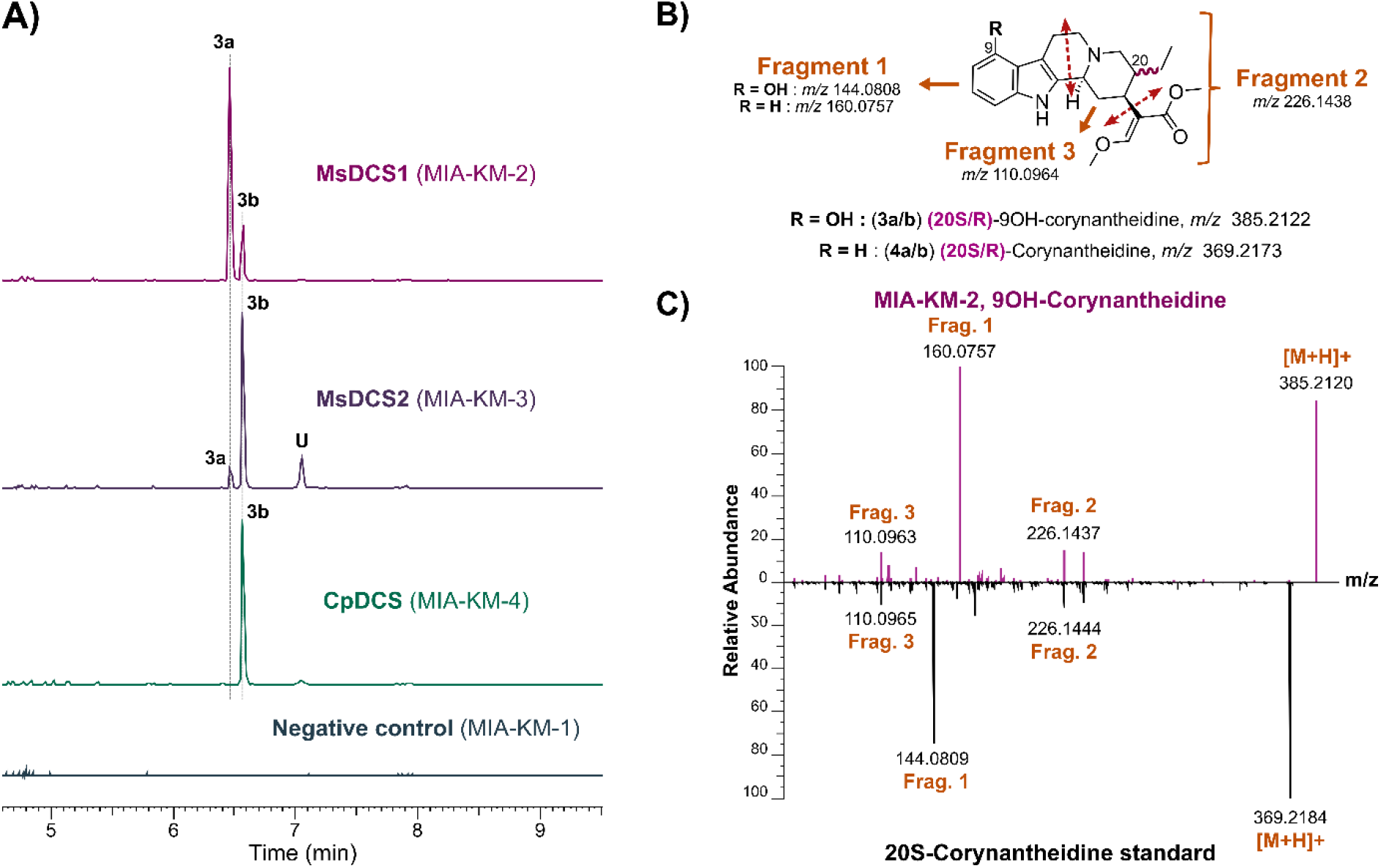
*De novo* biosynthesis of 9-OH-corynantheidine (3a/b) in yeast. **A)** Extracted Ion Chromatogram (EIC) for m/z = 385.2122 (5 ppm mass resolution) obtained upon high-resolution LC-MS/MS of supernatants from MIA-KM-1 to MIA-KM-4. **B)** MS/MS fragmentation pattern of (9OH-) corynantheidine. **C)** Representative MS/MS spectra of 20S-corynantheidine standard and 9-OH-corynantheidine produced by MIA-KM-2 to MIA-KM-4. The MS/MS fragmentation patterns are identical for 20R and 20S stereoisomers.

In summary, we report here *de novo* production of **3a/b** in a heterologous platform with a synthetic *Pc*PsiH-based pathway shortcutting an unknown enzyme in the canonical kratom biosynthesis pathway. In yeast, the kratom biosynthetic enzymes show remarkable promiscuity relative to the artificially introduced C9-OH group in the corynanthe skeleton. The ratio of peak area between (**6a/b**) and (**4a/b**) was 1:1.5 for *Ms*DCS2, 1:4.7 for *Cp*DCS, and 1:8.3 for *Ms*DCS1, indicating that a substantial part of the carbon flux goes through the designed synthetic pathway. The differences in ratio observed can be attributed to the different promiscuity of these enzymes towards the non-natural substrate 9-OH-strictosidine aglycone (**8**).

### De novo production of mitragynine and speciogynine in yeast

To complete the full refactoring of the synthetic pathway for *de novo* production of MG or SG, we next attempted to identify the enzyme responsible for the last C9 O-methylation in kratom. We generated an RNA-seq based transcriptome data set of different *M. speciosa* plant tissues and at different developmental stages (**Supp. Fig. 4**). Based on analysis of differential gene expression between leaf samples harvested at different developmental stages and between leaves with either red or green veins, we identified 11 putative OMTs (**Supp. Fig. 5**). In addition to these gene candidates, we also explored the opportunity to identify genes encoding OMTs from another recent RNAseq data set (Brose et al., 2021) (7 candidates), as well as OMTs from other resources identified based on homology to OMTs from plants producing MIAs (12 candidates) (**Supp. Fig. 5**). The 30 selected *Ms*OMT gene candidates were individually cloned on 2µ yeast expression plasmids and transformed into MIA-KM-2 producing both **3a** and **3b**, substrates of the candidate OMTs. Unfortunately, none of the tested enzymes led to production of MG or SG (data not shown). Nonetheless, the transcriptome information obtained could still be of interest for the kratom research community and used in complement to already published datasets (Brose et al., 2021; Pootakham et al., 2022) for further bioprospection.

Next, we instead adopted the enzyme from firebush *Hamelia patens* named *Hp*9OMT, which is capable of methylating **3a/b** to yield MG and SG and was discovered during the course of this study (K. Kim et al., 2023). Thus, the gene encoding *Hp*9OMT was introduced in our **3a/b** *de novo* producers MIA-KM-2, MIA-KM-3 and MIA-KM-4 yielding MIA-KM-5, MIA-KM-6 and MIA-KM-7, respectively. The resulting strains produced two different peaks at m/z = 399.2278 that eluted at the same retention time and had identical MS/MS fragmentation as authentic analytical standards of MG (RT = 7.04 min) and SG (RT = 7.09 min) (**Fig. 3A**). Thus, we accomplished *de novo* production of MG and SG from simple feedstocks in yeast cell factories. MIA-KM-5 with *Ms*DCS1 produced 1.4 ± 0.1 µg/L MG and 1.7 ± 1.0 µg/L SG, whereas the highest SG production was achieved by MIA-KM-7 expressing *Cp*DCS with 6.3 ±1.4 µg/L.

**Figure 3.**
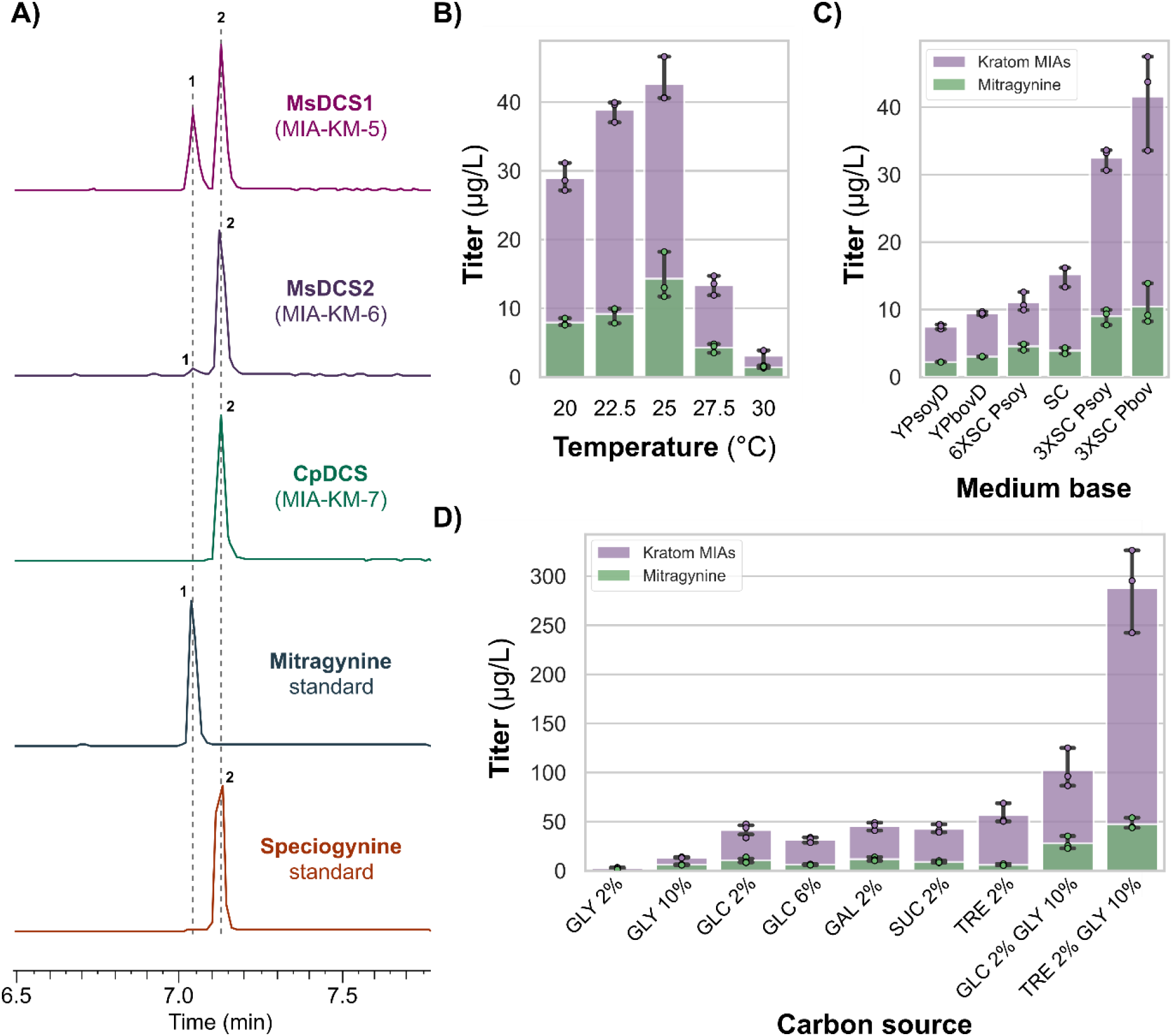
Yeast cell factory for *de novo* production of mitragynine and speciogynine. **A)** Detection of *de novo* mitragynine (MG) and speciogynine (SG) in MIA-KM-5 to MIA-KM-7 supernatants analyzed by LC-MS/MS (EIC m/z = 399.2278, 5 ppm mass resolution). **B)** Optimization of temperature for MG and SG production in MIA-KM-5 (“kratom MIAs” represents MG + SG). **C)** Optimization of media base for MG and SG production in MIA-KM-5. Abbreviations: SC, synthetic complete medium; YPD, yeast extract peptone dextrose medium; Psoy, soy peptone; Pbov, bovine peptone. **D)** Carbon source screen for MG and SG production in MIA-KM-5. Abbreviations: GLY, glycerol; GLC, glucose; GAL, galactose; SUC, sucrose; TRE, trehalose. Measurements for each condition represent the average of three biological replicates. Error bars represent S.D. +/− the mean.

### Fermentation optimization to improve kratom MIAs production

We demonstrated *de novo* production of MG and SG in yeast, but the initial titers obtained were low. Therefore, we explored the effect of different cultivation temperature, base media and carbon source on the MG/SG production strain MIA-KM-5. Different temperatures ranging from 20 to 30 °C were assessed in reference production media (3×SC, 3 mM tryptophan, 10 g/L soy peptone, 2 % glucose) for cultivation of MIA-KM-5 over 144 h (**Fig. 3B**). The optimal production temperature was 25 °C with 14.3 ± 2.8 µg/L MG and 28.3 ± 0.7 µg/L SG, representing >13-fold increase in kratom MIAs production compared to 30 °C. This is probably due to improved folding and activity of bottleneck pathway enzymes in yeast, especially SLS in accordance with recent work on the production of alstonine and serpentine in yeast (Bradley et al., 2023). Surprisingly, MIA-KM-5 produced significantly more SG than MG from glucose at 25 °C while accumulating more 20S-corynantheidine (MG branch) (**Supp. Fig. 6**). This was not observed by Kim et *al*., (2023) who showed production of 145 µg/L MG as the main product from bioconversion of 10 mg/L secologanin and tryptamine in yeast upon co-expressing *Ms*DCS1, *Pc*PsiH, *Cr*STR, *Cr*SGD, *Eno*lMT and *Hp*9OMT. Our findings could indicate a condition-dependent stereoselectivity of DCS and/or a substrate preference of *Hp*9OMT in these conditions for the 20R-9OH-corynantheidine over the 20S isomer. Differences in the DCS stereoselectivity between *in vitro, in planta* and yeast assays have been reported (K. Kim et al., 2023; Schotte et al., 2023). Notably, Kim et *al*., (2023) expressed heterologous genes in yeast on high-copy 2µ plasmids, whereas we integrated the genes into the yeast genome and our production conditions varied significantly.

Having fixed the production temperature at 25 °C, we next investigated different base media and peptone supplementation. Specifically, synthetic complete (SC) medium at different concentrations and complex medium (Yeast extract and peptone, YP) were investigated with different peptone origins (**Fig. 3C**). The best combination was 3×SC base supplemented with bovine peptone (Pbov), yielding 10.4 ± 2.5 µg/L MG and 31.2 ± 4.2 µg/L SG. This is in accordance with our previous work on production of MIAs in yeast (Bradley et al., 2023; Zhang et al., 2022).

We then assessed different carbon sources in the best base medium (3×SC, 3 mM tryptophan, 10 g/L Pbov) including glucose, galactose, sucrose, glycerol, and trehalose at different concentrations (**Fig. 3D**). Additionally, we attempted to combine glycerol with sugar carbon sources which had been shown to improve production of noscapine, a member of the benzylisoquinoline alkaloid (BIA) family found in plants, by yeast (Li et al., 2018). The highest titer was obtained combining trehalose 2% and glycerol 10% with production of 47.5 ± 4.6 µg/L MG and 240.6 ± 34.6 µg/L SG, representing a 7-fold increase in total kratom MIAs titer compared to 2% glucose used before. This results from a synergistic effect of 10% glycerol and 2% trehalose. Cultivation of MIA-KM-5 with 10% glycerol alone only led to production of 13.5 ± 0.4 µg/L kratom MIAs, while growth monitoring showed that it also contributed very marginally to biomass formation compared to sugar substrates (**Supp. Fig. 7)**. Glycerol assimilation leads to NADPH generation used by the multiple MIA pathway oxidoreductases as cofactor (Costenoble et al., 2000). In addition, it has been shown that glycerol and trehalose can act as chemical chaperones facilitating proper protein folding, localization and preventing aggregation – all of which are common issues when transplanting complex plant pathways in heterologous microbial hosts (Perlmutter, 2002; Sato et al., 1996; Singer & Lindquist, 1998; Tapia & Koshland, 2014). Furthermore, trehalose assimilation rate in yeast has been shown to be controlled by the rate of extracellular trehalose hydrolysis, mimicking the conditions of glucose-limited continuous cultures which potentially promotes MIA production by reducing overflow metabolism (Jules et al., 2004). In the optimized batch conditions, MIA-KM-7 produced 172.6 ± 12.2 µg/L SG (**Supp. Fig. 8**).

Altogether, the best media and cultivation conditions led to >90-fold improvement in kratom MIAs titers in DW96 plates, highlighting the importance of cultivation condition optimization to improve production of complex plant natural products in a microbial host. Importantly, the greatest improvements were obtained by reducing the temperature to 25 °C and switching to a trehalose/glycerol carbon source mix. These changes most likely had an impact on the folding and activity of heterologous enzymes, the supply of cofactors, and the global regulation of metabolism (Jules et al., 2004; Xiberras et al., 2019).

### Untargeted metabolomics reveals multiple shunt products associated with DCS and STR promiscuities

During the optimization of fermentation conditions, we observed the formation of multiple side products with MIA-like MS/MS fragmentation in the total ion count (TIC) from LC-MS/MS data acquired. In an effort to systematically identify these side products, we processed the untargeted metabolomics data acquired for the different strains described in this paper using MZmine (Pluskal et al., 2010; Schmid et al., 2023) and generated feature-based molecular networks based on MS/MS spectra (Nothias et al., 2020; Wang et al., 2016). We found clusters containing multiple previously identified kratom MIAs as well as products whose spectra was matched with MIAs in public databases. The different clustered peaks were manually inspected, and SIRIUS was used for molecular structure prediction (Dührkop et al., 2019). The predicted structures were evaluated based on knowledge on the biochemistry of the MIA pathway.

Two types of shunt products were identified. The first ones originate from the ability of STR to accept other aldehydes than secologanin for the Pictet-Spengler condensation reaction with tryptamine leading to formation of 1,2,3,4-tetrahydro-β-carboline (THBC) scaffold (Eger et al., 2020; Maresh et al., 2008; Sheng & Himo, 2020) (**Supp. Fig. 9)**. The canonical product of this reaction is tetrahydroharman (THH), resulting from condensation of tryptamine with acetaldehyde naturally produced by yeast (Aranda & del Olmo, 2004). Interestingly, we also identified putative 5-OH-THH and 5-MeO-THH resulting from condensation of *Pc*PsiH-derived 4-OH-tryptamine with acetaldehyde and subsequent O-methylation by *Hp*9OMT. 5-MeO-THH is an isomer of tetrahydroharmine (7-MeO-THH), a psychedelic bioactive of ayahuasca (Grella et al., 1998). This STR promiscuity towards their carbonyl substrate is generalizable to other Pictet-Spenglerases as shown for norcoclaurine synthase involved in the biosynthesis of plant benzylisoquinoline alkaloids in yeast (Pyne et al., 2020). Regarding kratom MIAs production, this represents a 4-OH-tryptamine sink and a cofactor waste, with both NADPH being used by *Pc*PsiH/*Pc*CPR and S-adenosyl methionine (SAM) by *Hp*9OMT. It was suggested that tryptamine adopts a different binding position in the STR active site when reacting with a short aliphatic chain than when reacting with secologanin, implying that STR specificity could be engineered in order to improve the flux of the pathway (Eger et al., 2020).

The other class of shunt product identified originates from the reduction of strictosidine aglycone (**7**) by DCS (**Supp. Fig. 10)**. In MIA plants, strictosidine is deglycosylated by SGD, yielding a reactive dialdehyde, coexisting as several isomers in chemical equilibrium that is then stabilized through reduction by medium chain alcohol dehydrogenases (ADHs) such as DCS. Depending on which isomer is reduced, different MIA scaffolds can be formed (Gerasimenko et al., 2002; O’Connor & Maresh, 2006). In MIA-KM-5 (*Ms*DCS1) samples, we identified tetrahydroalstonine, rauwolscine, yohimbine and allo-yohimbine peaks matching retention time, m/z and MS/MS fragmentation pattern of authentical analytical standards. We also identified production of putative geissoschizine methyl ether (GME) by MIA-KM-5, hinting that *Ms*DCS1 produces geissoschizine that is subsequently O-methylated at C17 by EnolMT. These activities were just reported by Kim *et al*. (2023), who identified these products to be of the 19Z-form. We further report here that MIA-KM-5 produces putative 9-OH-GME and 9-MeO-GME, suggesting that 4-OH-tryptamine gets also channeled into this pathway branch, and that the resulting 9-OH-GME gets methylated by *Hp*9OMT. This observation was also made for yohimbane-type and heteroyohimbane-type MIAs with apparition of two 9-MeO-yohimbane peaks ([M+H]^+^ m/z=385.2122, RT = 6.71 min and 6.85 min respectively) as well as 9-MeO-tetrahydroalstonine ([M+H]^+^ m/z=383.1965). Overall, DCS promiscuity leads to important 4-OH-tryptamine and secologanin sinks, the latter being the main bottleneck identified in our previous yeast-based MIA production effort (Bradley et al., 2023; Zhang et al., 2022). *Ms*DCS1 would therefore be an important enzyme to engineer to improve microbial production of MG by *i)* altering its stereoselectivity to improve the MG/SG ratio and *ii)* reducing production of MIA shunt products that exhaust precursors and cofactors pools. MIA ADH promiscuity is an issue that is generalizable beyond kratom (Bradley et al., 2023, 2024; Stander et al., 2023; Stavrinides et al., 2016) and important enzyme engineering work must be carried to allow efficient conversion of strictosidine into specific downstream MIA scaffolds in microbial cell factories.

### Engineered yeast as platform for derivatization of kratom MIAs

The classical approach to drug discovery begins with hit-to-lead campaigns based on a pharmacophore that interacts with a target, then onto semi-synthetic lead optimization, followed by screening for enhanced pharmacological properties. Nevertheless, producing analogs of plant-based natural product drug candidates presents difficulties due to the complex structures often encountered in plant natural products, and due to synthetic chemistry constraints (Bradley et al., 2020; Hong et al., 2020). Having demonstrated *de novo* kratom MIAs production in yeast, we next sought to explore biosynthesis of kratom alkaloid analogs using cell-factories. We first became interested in hydroxylating MG to form 7-OH-MG because it is also a partial agonists of MOR, about 10-fold more potent than MG (Kruegel et al., 2016, 2019; Kruegel & Grundmann, 2018). In humans, orally absorbed MG undergoes hydroxylation mediated by *Hs*CYP3A4 in the liver as part of its degradation pathway, resulting in production of active 7-OH-MG together with other side products (**Fig. 4A**) (Chakraborty, Uprety, et al., 2021; Kamble et al., 2019; Kruegel et al., 2019). Depending on the degree of conversion to 7-OH-MG, certain people with increased or decreased *Hs*CYP3A4 activity may have stronger or less enhancing effects after consuming MG or kratom (Kruegel et al., 2019). To avoid this variability effect and benefit from improved potency, this modification has been employed semi-synthetically in multiple drug discovery campaigns around the MG scaffold (Bhowmik et al., 2021; Chakraborty, DiBerto, et al., 2021; Gutridge et al., 2021; Matsumoto et al., 2014; Takayama et al., 2006). Therefore, we next integrated *Hs*CYP3A4 together with its reductase *Hs*CPR in the genome of MIA-KM-5 resulting in strain MIA-KM-8. This strain was cultivated for 144 h using the optimized conditions previously identified (**Fig. 3B**-**D**, 25°C, 3×SC, 3mM tryptophan, 1 % Pbov, 2 % trehalose, 10 % glycerol). Extracted ion chromatograms (EIC) from supernatant analysis at m/z = 415.22 led to apparition of several peaks compared to the parental strain. However, the signal was insufficient to acquire MS/MS fragmentation patterns and accurately identify the products (data not shown). As a mitigating method, MIA-KM-8 was grown in medium supplemented with 10 mg/L MG to boost the signal of *Hs*CYP3A4-derived bioconversion products and potentially permit MS/MS capture. In this setting, the production of 7-OH-MG was validated comparing RT, exact m/z, and MS/MS fragmentation with an authentic standard (**Fig. 4A**). Four other unknow compounds noted U1 to U4 were also produced while absent from the parental strain MIA-KM-5 (**Supp. Fig. 10)**. These most likely originate from the promiscuity of *Hs*CYP3A4, which has been shown to generate different mono-oxidation products of MG on the indoloquinolizine moiety and potentially oxoindole rearrangement products (Kamble et al., 2019, 2020).

While the activity of *Hs*CYP3A4 in MIA-KM-8 was low, we demonstrated the integration of human enzymes in yeast to derivatize bioactive kratom MIAs. Our work demonstrates how synthetic biology may be used to overcome evolutionary barriers and produce plant natural product analogs by expressing tailoring enzymes. Further work on *Hs*CYP3A4/*Hs*CPR expression optimization and balancing could allow *de novo* production of these oxidized kratom MIAs that could then be purified and tested on human opioid receptors.

**Figure 4.**
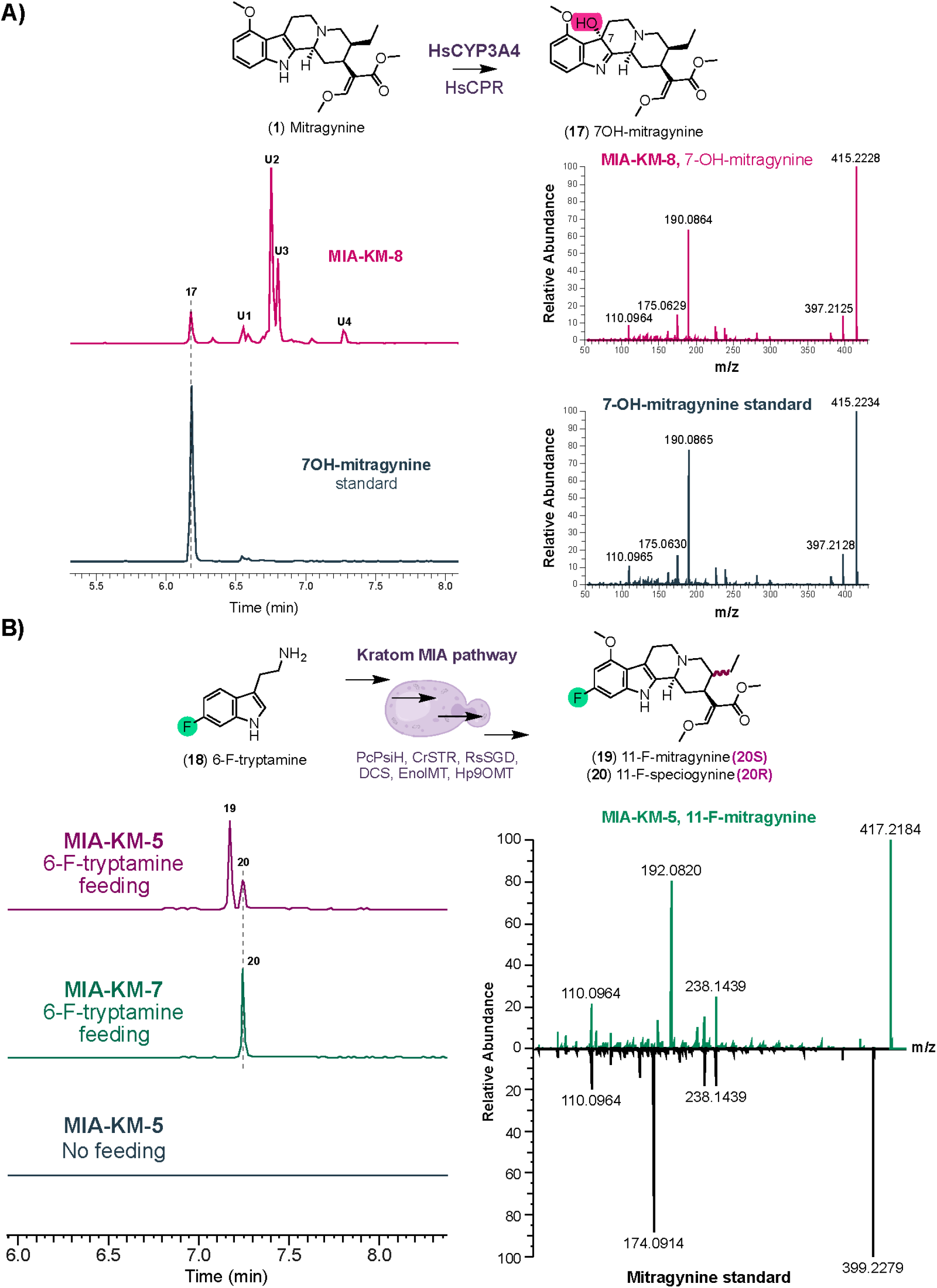
Kratom yeast cell factories as platforms for production of mitragynine (MG) and speciogynine (SG) derivatives. **A)** Expression of human cytochrome P450 *Hs*CYP3A4 and its reductase *Hs*CPR in MIA-KM-8 leads to production of 7-OH-mitragynine and other mono-hydroxylated MG derivatives (EIC m/z = 415.2227, 5 ppm mass resolution). **B)** Feeding 6-F-tryptamine to MIA-KM-5 and MIA-KM-7 allows production of 11-F-MG (**19**) and 11-F-SG (**20**) in yeast by exploiting pathway enzyme promiscuities.

Another way of derivatizing natural products using cell factories is to feed precursor analogs and harness potential substrate promiscuity of pathway enzymes. Halogenation in particular greatly impacts the physico-chemical properties of drugs, and halogen atoms are present in about 25% of licensed pharmaceuticals (Xu et al., 2014). Bhowmik et *al*. (2021) docked MG in human MOR and showed that the aromatic indole ring is pointing inside the receptor’s binding pocket, which is most likely crucial for binding. Targeting this indole ring with halogen atoms is therefore expected to strongly affect binding at the different opioid receptors as shown in several medicinal chemistry campaigns around the MG scaffold (Bhowmik et al., 2021; Matsumoto et al., 2014; Takayama et al., 2006). Thus, we next fed 6-F-tryptamine to MIA-KM-5 (*Ms*DCS1) and MIA-KM-7 (*Cp*DCS) cultivated in previously optimized conditions (25 °C, 3×SC, 3 mM tryptophan, 1 % Pbov, 2 % trehalose, 10 % glycerol) for 144 h. These cultures were analyzed by LC-MS/MS, and we observed two peaks that matched the exact masses of 11-F-MG and 11-F-SG, which were absent when no 6-F-tryptamine was supplemented. The MS/MS fragmentation for these peaks showed a mass addition of 17.99 Da, coherent with the substitution of a hydrogen atom with fluorine on the indole ring of the molecule (**Fig. 4B**). Two distinct peaks were observed for MIA-KM-5 while only one for MIA-KM-7, leading us to identify the first one (RT = 7.17 min) as 11-F-MG and the second one (RT = 7.24 min) as 11-F-SG. We therefore report here the first biotechnological production of fluorinated MG and SG showcasing substrate promiscuity of kratom pathway enzymes when expressed in microbial cells. Similarly, Schotte et *al*. (2023) reported production of fluorinated 9F-, 10F, and 11F-corynantheidine upon feeding secologanin and respectively 4F-, 5F-, or 6F-tryptamine in *Nicotiana benthamiana* leaves. However, these derivatives lacked the methoxy group in C9 that is essential for bioactivity of MG/SG at opioid receptors (Kruegel & Grundmann, 2018). While production of halogenated kratom MIAs with total chemical synthesis is highly labor intensive and not scalable due to the numerous reaction and chromatographic separation steps, late-stage halogenation of plant-extracted MG has been achieved for position C10 (Takayama et al., 2006) and more recently for C11 and C12 (Bhowmik et al., 2021). Bhowmik *et al*. (2021) reported the first suite of C11 halogenated MG derivatives using complex multistep chemical synthesis procedures from MG and showed that combining C11 fluorination with C7 hydroxylation resulted in promising lower efficacy opioids which are of great interest for the treatment of opioid use disorder. Overall, we showcase here that yeast cell factories can be an alternative source of halogenated kratom MIAs for drug discovery and lead optimization.

## Conclusion

In this work, we report the first *de novo* production of the kratom MIAs mitragynine and speciogynine in yeast cell factories using a synthetic pathway based on the fungal tryptamine 4-monooxygenase *Pc*PsiH. We further highlight the importance of optimizing media composition and cultivation conditions to improve plant natural product production in yeast with temperature and carbon source optimization having the highest impact on kratom MIA production in yeast yielding up to ∼290 µg/L. Analysis of the untargeted metabolomics data generated in this study led to identification of multiple side products generated by these strains owing to the promiscuity of STR and DCS. These unwanted products are a source of precursor and cofactor waste, and enzyme engineering in this pathway should prove instrumental in improving strain performance. Extending from the biomanufacturing of kratom MIAs, we demonstrate production of hydroxylated and fluorinated MG and SG analogs using yeast cell factories. Although further work is needed on engineering pathway enzymes, refining bioprocess development, and streamlining downstream processing of kratom MIAs analogs, this study sheds light on a microbial route to manufacture kratom MIAs as well as providing new-to-nature kratom MIA analogs of relevance for further drug discovery and development to mitigate unmet medical needs within pain, addiction, and depression.

## Materials & Methods

### Chemical standards

All chemical standards (**Supp. Table 4**) had a purity of 95% or higher.

### Plasmid and Yeast Strain Construction

All plasmids were assembled using USER (Nour-Eldin et al., 2006) or Golden Gate (Engler et al., 2008) cloning. The *Pc*PsiH/*Pc*CPR module was PCR amplified from pCFB9359 (Milne et al., 2020) and inserted into pCFB9359, a vector part of the EasyClone-MarkerFree expansion for CRISPR-Cas9 based genomic integration at site VIII-1 (Babaei et al., 2021). With the exception of the *Pc*PsiH/*Pc*CPR module, all the other pathway genes were cloned using the Yeast MoClo (modular cloning) toolkit (Lee et al., 2015). Assembly reactions were transformed into *E. coli* DH5α competent cells and plated on Luria–Bertani agar containing 100 μg.ml^−1^ ampicillin, 50 μg.ml^−1^ kanamycin or 25 μg.ml^−1^ chloramphenicol and grown at 37 °C overnight. Colonies were picked, grown overnight in 5 mL LB with appropriate antibiotics; the next day plasmid DNA was extracted (Nucleospin Plasmid, Macherey-Nagel) and sent for Sanger sequencing (Eurofins Genomics). The full list of plasmids used in this study is available in **Supp. Table 2**.

MoClo plasmids followed the hierarchical DNA construction workflow as described in the original Yeast MoClo publication (Lee et al., 2015). Coding sequences of pathway genes were codon-optimized for *S. cerevisiae* and synthesized by Integrated DNA Technologies with relevant flanking type IIS restriction enzyme sites and overhangs. These synthesized fragments were assembled into the Level 0 entry vector pYTK001 using the NEBridge Golden Gate Assembly Kit (BsmBI-v2) (New England Biolabs) following the manufacturer’s protocol. Level 1 plasmids were built by combining a promoter, CDS and terminator part into pre-assembled pMHGG1_Vec suite containing a yeast URA selection marker, an *E. coli* ampicillin resistance gene, a 2µ yeast origin of replication, connectors determining position of transcription unit in pathway and a GFP dropout cassette for visual screening of correct assembly in *E. coli* (NEBridge Golden Gate Assembly Kit BsaI-v2). Finally, these Level 1 transcription unit plasmids were combined with BsmbI-v2 into integrative Level 2 vectors pMHGG2_Vec custom built in this study to target genomic sites II-1, IV-1 or XIII-1 described in the EasyClone-MarkerFree expansion (Babaei et al., 2021). The Level 2 EasyClone-MarkerFree integration compatible vector were built by expanding the work of (Otto et al., 2021) who extended the Yeast MoClo toolbox for genomic integration, generating similar plasmids targeting the sites available in the original EasyClone-MarkerFree paper (Jessop-Fabre et al., 2016). We needed to access more integration sites as the base strain MIA-CZ-1 used in this study is heavily engineered and already had constructs in all the sites of the original EasyClone-MarkerFree work.

All the integrative plasmids generated in this study were linearized using NotI (New England Biolabs) and the DNA fragments were transformed in yeast together with relevant gRNA plasmid from EasyClone-MarkerFree (Jessop-Fabre et al., 2016) using standard lithium acetate methods (Gietz & Schiestl, 2007). The integration of heterologous genes was verified as previously described (Babaei et al., 2021; Jessop-Fabre et al., 2016).

### Media and yeast cultivation

All yeast strains used and constructed in this study (**Supplementary Table 3**) are based on CEN.PK2-1C. The *de novo* base strictosidine platform MIA-CM-5 used to build MIA-CZ-1 was described by (Zhang et al., 2022). For routine propagation and transformation purposes, yeast strains were grown at 30°C in YPD (yeast extract 1 % w/v, peptone 2 % w/v and glucose 2 % w/v). Unless specified otherwise, strains were grown in 3xSC medium supplemented with 3mM tryptophan, and optionally 1% w/v peptone if specified to test for MIA production. Media pH was adjusted to 5.5 with 2 M NaOH and filter sterilized. Strains were inoculated from plate in 1 mL of media matching the production media in 15 mL preculture tubes and incubated overnight at 30°C and 300 r.p.m shaking. The next day 100 µL of preculture was used to inoculate 500 µL of matching media in a 96-deep well plate. 3mM 6F-tryptamine was supplemented in the case of fluorinated kratom MIAs production. All cultivations were run as biological triplicates originating from different colonies. Plates were incubated at specified temperature for 144 h with shaking at 300 r.p.m. Following 144 h, 100 μL of each sample was filtered through a filter plate (PALL, AcroPrep Advance, 0.2-μm Supor membrane for medium/water) by centrifugation at 2,200 g for 1.5 min. Samples were stored at -20 °C prior to injection.

### Fed-batch fermentation of MIA-CZ-1 for strictosidine production

Batch-fed fermentations were performed on an Ambr 250 system (Sartorius) using single-use microbial vessels. The fermentation started by inoculating 100 ml of 3xSC + 3 mM Trp + 2% glucose to an initial OD600 of 0.5. The same medium, except containing 36% glucose, was fed continuously into the bioreactor between 20 and 144h using a linear feeding profile:

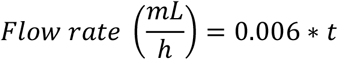

The temperature was kept at 30 °C throughout the whole fermentation. The pH of all bioreactors was controlled at 5.0 using 10% NH4OH solution. The air flow rate was initially at 1 volume of air per unit of medium per unit of time (vvm), and dissolved oxygen (dO2) was maintained above 40% by increasing the agitation speed. CO2 in the exhaust gas was monitored by a gas analyser. An automated liquid handler was used to take 1.0 ml broth samples every 24 h for metabolite analysis.

### Strain growth characterization with microtiter plate reader

To characterize strain growth with different carbon sources, three single colonies were inoculated separately from plate into 3 mL of tested media (3xSC, 1 % bovine peptone, 3 mM tryptophan, pH 5.5 supplemented with tested carbon source) in 15 mL preculture tubes overnight (30 °C, 300 r.p.m). The next day, the optical density at 600 nm (OD600) was measured and cells were diluted in matching media at a starting OD600 = 0.5. 150 µL of each cell resuspension was transferred to a clear-bottomed 96-well plate covered with a Breathe-Easy sealing membrane (Diversified Biotech BEM-1) and incubated in a SynergyMX microtiter plate reader (BioTek) at 25 °C and 250 r.p.m. Growth curves were processed and analyzed using a QurvE (Wirth et al., 2023).

### Metabolite analysis by liquid chromatography-high resolution tandem mass spectrometry

Samples generated in this study were analyzed using an untargeted metabolomics system constituted of a Vanquish Duo UHPLC binary system (Thermo Fisher Scientific, USA) connected to an Orbitrap ID-X Tribrid mass spectrometer (Thermo Fisher Scientific. USA). Chromatographic separation was achieved under reverse-phase conditions as previously described (Bradley et al., 2023; Kildegaard et al., 2021). The MS measurements were performed in positive-heated electrospray ionization mode with a voltage of 3,500 V acquiring the full MS/MS spectra (data-dependent acquisition-driven MS/MS) in the mass range of 70–1,000 Da. The following data-dependent acquisition settings were used: automatic gain control target value of 4 × 10^5^ for full-scan MS and 5 × 10^4^ for the MS/MS spectral acquisition and a mass resolution of 120,000 for full-scan MS and 30,000 for MS/MS events. Precursor ions were fragmented by stepped high-energy collision dissociation using collision energies of 20, 40 and 50. In general, standards for non-natural hydroxylated or fluorinated MIAs are not available commercially; instead, they were identified using their exact mass, retention time, and shifts in MS/MS spectra with respect to the non-hydroxylated or non-fluorinated standard as done previously (Bradley et al., 2023; Misa et al., 2022; Schotte et al., 2023).

### Untargeted metabolomics data processing

LC-MS data were qualitatively inspected using the FreeStyle Software (Thermo Fischer Scientific, USA). The peaks integration was done using Thermo Xcalibur Quant Browser Software (Thermo Fisher Scientific, USA) using 5-ppm mass tolerance. The integrated peaks were also manually inspected to ensure correct peak assignment and integration. For shunt product identification the raw files were processed using MZmine 2.53 (Pluskal et al., 2010; Schmid et al., 2023) to generate a table of metabolic features. The generated feature files were used as input for GNPS FBMN (Feature-Based Molecular Networking) which is one of the most significant and often used tools for molecular networking, annotation, and visualization in the field of metabolomics (Nothias et al., 2020; Wang et al., 2016). Features of interest found in clusters containing known MIAs were computed using the SIRIUS executable for formula and structural predictions (Dührkop et al., 2019).

### Transcriptomics on M. speciosa tissue

Two sets of kratom leaves, red- and green-veined, were collected from two-year-old plants in Hat Yai, Songkhla province, Thailand, during July 2021. Herbarium specimens of red-veined and green-veined kratom, designated as N5/001 (PSU) and N1/001 (PSU), were deposited at the Department of Biology, Faculty of Science, Prince of Songkla University, Thailand. Different developmental stages of kratom leaves were cut from the apex: 1-pair, 2-pair, and 3-pair leaves, and analyzed for the contents of mitragynine, speciogynine, and paynantheine using the HPLC-UV method (Sengnon et al., 2023) (**Supp. Fig. 4**). All leaves were freshly cut, cleaned with sterile water, and immediately frozen in liquid nitrogen. They were stored at -80°C until used.

### RNA extraction and sequencing

RNA was extracted from frozen *Mitragyna speciosa* leaf tissue samples using PureLink Plant RNA Reagent according to the manufacturer’s instructions (Thermo Fisher Scientific, USA). Integrity of the RNA was checked using the TapeStation 4200 system (Agilent Technologies, USA) and the RNA concentration was measured using the Qubit 3.0 Fluorometer (Thermo Fisher Scientific, USA). Barcoded RNA-seq libraries were prepared from 1 μg of total RNA using the TruSeq Stranded mRNA Library Prep kit according to the manufacturer’s instructions (Illumina Inc., USA). RNA-seq libraries were sequenced on an Illumina NovaSeq 6000 System according to the manufacturer’s protocol. Image analysis and base calling were done by the Illumina pipeline (Illumina Inc., USA). RNA-seq data yield varied from ∼34 to ∼90 million paired-end 2 × 150 bp reads per sample, corresponding to ∼10 to ~27 Gb per sample (**Supp. Fig. 4**).

### Selection of Ms9OMT gene candidates

Two distinct sample groups were employed for differential gene expression analysis: red-versus green-veined leaves (A vs C), focusing on mitragynine down-regulation, and first versus second and third leaves, focusing on mitragynine up-regulation (**Supp. Fig. 4**). RNA-seq data was analyzed using DESeq2 by Future Genomics (Leiden, The Netherlands) to obtain log2FoldChange and base mean values. Data was visualized to identify significantly regulated genes between sample groups that were classified as methyltransferases using Pfam (Finn et al., 2008). RNA sequences of these candidates were translated into protein sequences using the longest open reading frame, and sequences that were too short were further analyzed using Basic Local Alignment Search Tool (BLAST) against a previously published kratom transcriptome (Brose et al., 2021). Finally, a protein BLAST was carried out to identify the best candidates to be tested experimentally based on sequence similarity with characterized methyltransferases. The full bioinformatic pipeline is detailed in **Supp. Fig. 5A**.

## Supporting information

Holtz et al, 2024 Supplementary Information

## Acknowledgements

This work was supported by the Novo Nordisk Foundation Copenhagen Bioscience Ph.D. Program grant No. NNF22SA0078231, Novo Nordisk Foundation grant No. NNF20CC0035580 and European Union Horizon 2020 research and innovation program grant number 814645 (MIAMi).

## Data availability

RNA-seq data have been deposited in the National Center for Biotechnology Information (NCBI) Gene Expression Omnibus (GEO) database under the study ID GSE268113 (accession numbers GSM8285926 - GSM8285941).

