## Supplementary material for "Metabolic engineering of yeast for *de novo* production of kratom monoterpene indole alkaloids": Holtz et al, 2024 Supplementary Information

### Contents of Supplementary Information

**Supplementary Figure 1.** Performance of strictosidine platform strain CZ-1 in deepwell plate and Ambr 250mL bioreactor

**Supplementary Figure 2.** 4-OH-tryptamine and 9-OH-strictosidine production by MIA-KM-1

**Supplementary Figure 3.** Corynantheidine and dihydrocorynantheine produced by MIA-KM-2, MIA-KM-3 and MIA-KM-4

**Supplementary Figure 4.** RNA-seq transcriptome of *Mitragyna speciosa* experimental setup and quality metrics

**Supplementary Figure 5.** Candidate MsOMT gene candidates screened and selection strategy

**Supplementary Figure 6.** Stereochemistry of different pathway intermediates generated by MIA-KM-5

**Supplementary Figure 7.** Growth characterization of MIA-KM-5 on trehalose/glycerol mix

**Supplementary Figure 8.** Production of speciogynine by MIA-KM-7 in optimized DW96 batch conditions

**Supplementary Figure 9.** STR-derived shunt products identified in kratom yeast strains using untargeted metabolomics data processing pipeline.

**Supplementary Figure 9.** DCS-derived shunt products identified in kratom yeast strains using untargeted metabolomics data processing pipeline.

**Supplementary Figure 11.** MS/MS fragmentation of HsCYP3A4-generated oxidation products.

**Supplementary Table 1.** Gene sequences used in this study.

**Supplementary Table 2.** Plasmids used in this study

**Supplementary Table 3.** Strains used in this study

**Supplementary Table 4.** Chemical standards used in this study

**Supplementary Table 5.** Metabolite retention times, masses and identifying fragments used in LC-MS/MS.

### Supplementary Figures

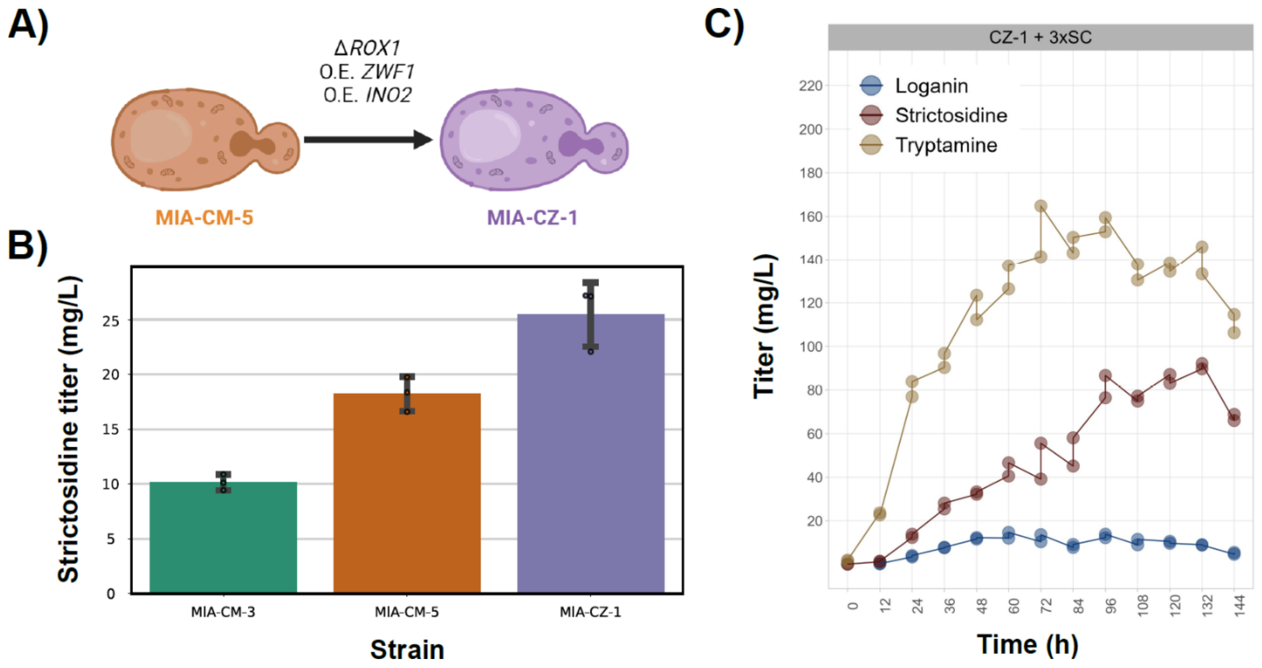

**Supplementary Figure 1.** Performance of strictosidine platform strain CZ-1 in Deepwell plate and Ambr 250mL bioreactor. **A)** Construction of MIA-CZ-1 from MIA-CM-5. **B)** Strictosidine titer produced by MIA-CZ-1 *de novo* in DW96 compared to previously reported strictosdine platform strains**. C)** MIA-CZ-1 fermentation in 250mL ambr fed-batch bioreactor with glucoses as carbon source.

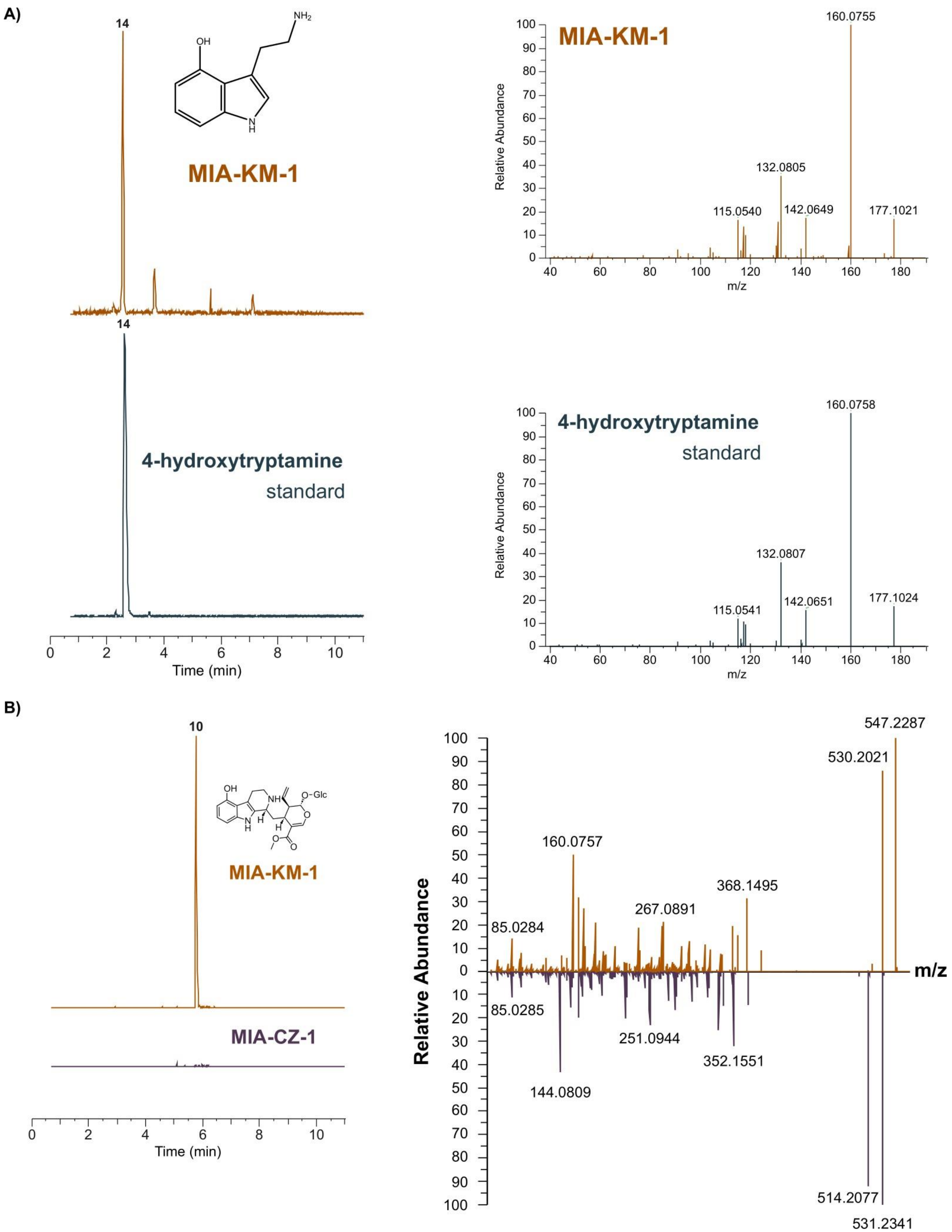

**Supplementary Figure 2.** 4-OH-tryptamine **(A)** and 9-OH-strictosidine **(B)** production by MIA-KM-1.

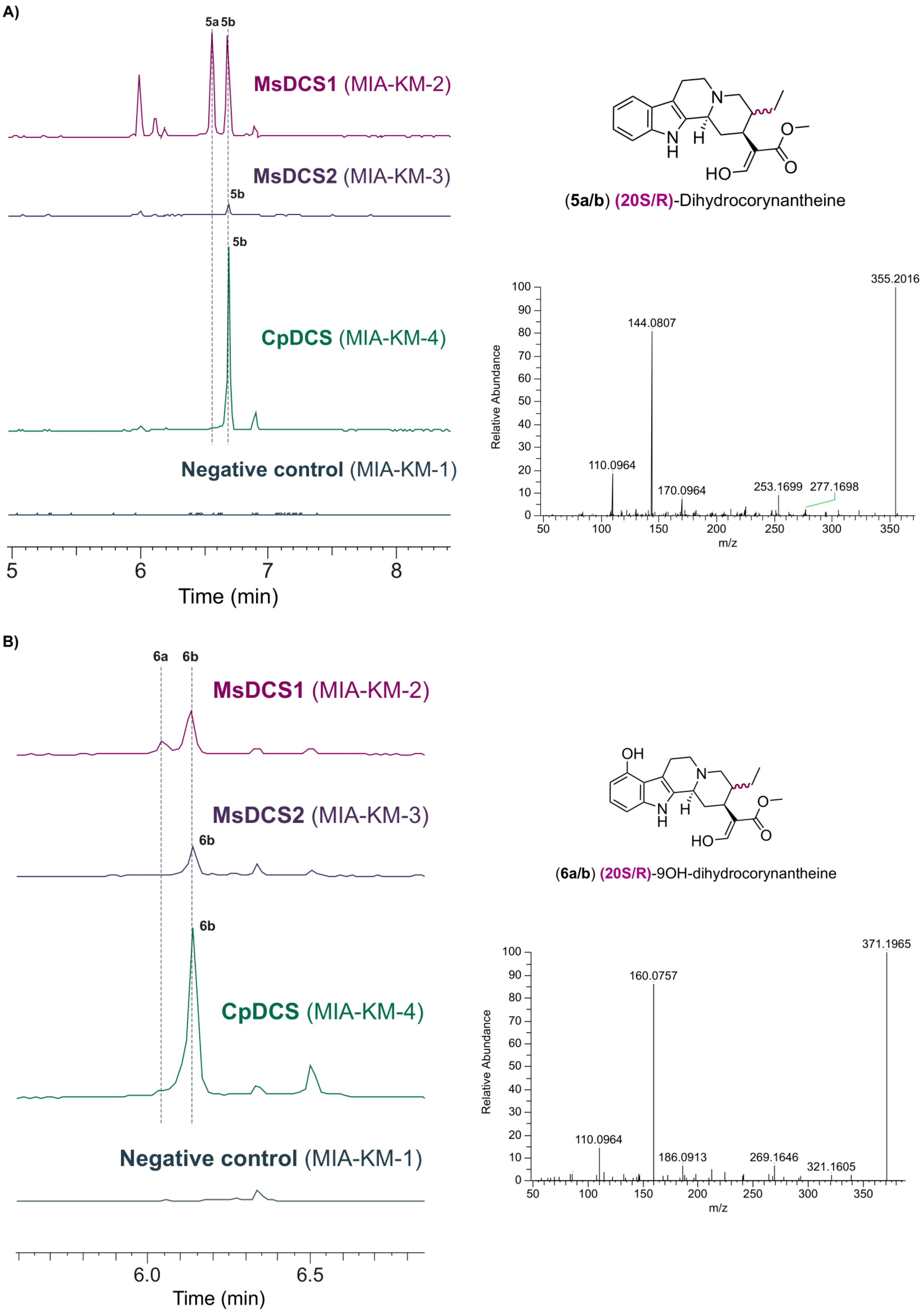
A)
5a
5b
MsDCS1 (MIA-KM-2)
MsDCS2 (MIA-KM-3)
CpDCS (MIA-KM-4)
Negative control (MIA-KM-1)
Time (min)
(5a/b) (20S/R)-Dihydrocorynantheine
355.2016
144.0807
110.0964
170.0964
253.1699
277.1698
Relative Abundance
m/z
B)
6a
6b
MsDCS1 (MIA-KM-2)
MsDCS2 (MIA-KM-3)
CpDCS (MIA-KM-4)
Negative control (MIA-KM-1)
Time (min)
(6a/b) (20S/R)-9OH-dihydrocorynantheine
371.1965
160.0757
110.0964
186.0913
269.1646
321.1605
Relative Abundance
m/z

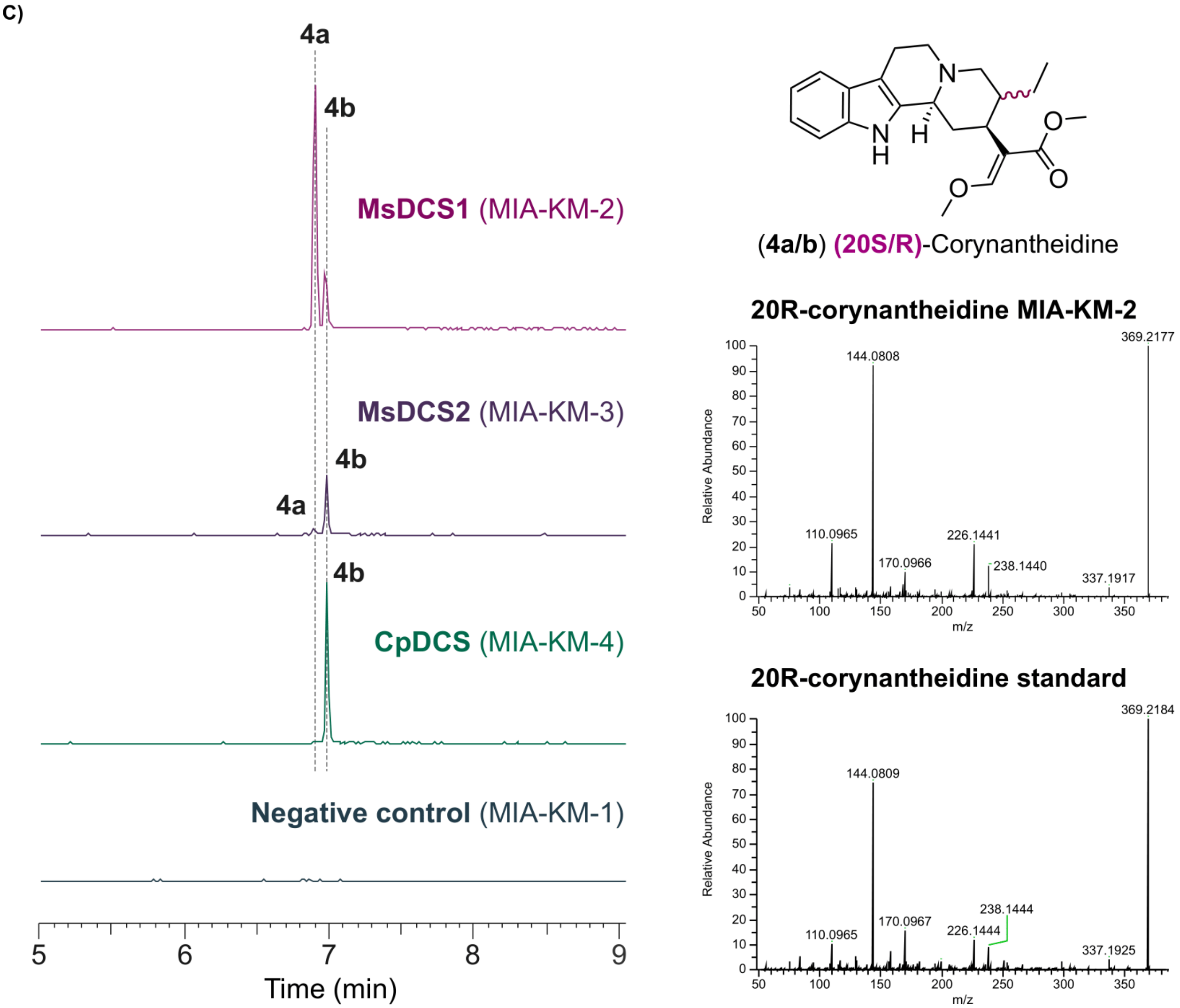

**Supplementary Figure 3.** Dihydrocorynantheine **(A)**, 9-OH-dihydrocorynantheine **(B)** and corynantheidine **(C)** produced by MIA-KM-2, MIA-KM-3 and MIA-KM-4

A)

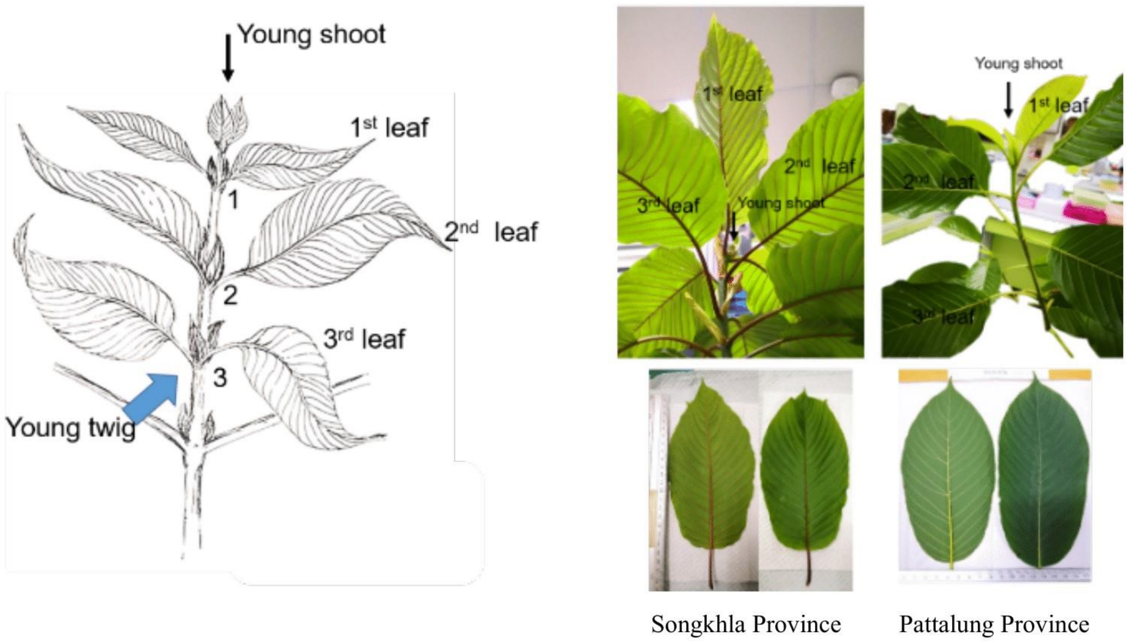

B)

**Alkaloid contents of wild-kratom leaves**

| Kratom leaf | Alkaloid contents (mg/g dry weight) | | |
|---|---|---|---|
| | Mitragynine | Speciogynine | Paynantheine |
| Set A: Red-veined | | | |
| 1st leaf | 9.383 ± 0.312 | 0.821 ± 0.031 | 2.303 ± 0.086 |
| 2nd leaf | 13.852 ± 0.656 | 1.240 ± 0.026 | 2.808 ± 0.072 |
| 3rd leaf | 12.111 ± 0.733 | 0.961 ± 0.067 | 2.244 ± 0.179 |
| Set B: Red-veined | | | |
| 1st leaf | 11.679 ± 1.478 | 1.112 ± 0.166 | 2.989 ± 0.415 |
| 2nd leaf | 12.180 ± 1.111 | 0.877 ± 0.094 | 2.356 ±0.251 |
| 3rd leaf | 13.912 ± 0.543 | 1.107 ± 0.040 | 2.475 ± 0.066 |
| Set C: Green-veined | | | |
| 1st leaf | 5.947 ± 0.106 | 0.716 ± 0.204 | 1.954 ± 0.455 |
| 2nd leaf | 11.604 ± 1.229 | 0.915 ± 0.095 | 2.300 ± 0.258 |
| 3rd leaf | 10.764 ± 0.897 | 0.912 ± 0.083 | 2.143 ± 0.183 |
| Set D: Green-veined | | | |
| 1st leaf | 11.469 ± 0.819 | 1.698 ± 0.093 | 3.777 ± 0.178 |
| 2nd leaf | 11.328 ± 0.983 | 1.042 ± 0.095 | 2.192± 0.240 |
| 3rd leaf | 9.214 ± 0.668 | 0.875 ± 0.069 | 1.719 ± 0.174 |

C)

| Sample # | Sample code | # clusters | yield (MB) | Mapped reads against reference | Mapped reads against reference (%) |
|---|---|---|---|---|---|
| FG2160_17 | set A 1st leaf | 39,290,582 | 11,865 | 26,733,250 | 68.0 |
| FG2160_18 | set A 2nd leaf | 59,028,327 | 17,826 | 39,308,040 | 66.6 |
| FG2160_19 | set A 3rd leaf | 42,278,491 | 12,768 | 27,881,642 | 65.9 |
| FG2160_20 | set C 1st leaf | 89,569,743 | 27,050 | 60,391,917 | 67.4 |
| FG2160_21 | set C 2nd leaf | 47,767,370 | 14,425 | 33,030,134 | 69.1 |
| FG2160_22 | set C 3rd leaf | 42,021,440 | 12,690 | 28,656,784 | 68.2 |

**Supplementary Figure 4.** RNA-seq transcriptome of *Mitragyna speciosa* experimental setup and quality metrics. **A)** Plant tissue sourcing for RNA extraction. **B)** Kratom MIA content in leaf samples determined by HPLC-UV **C)** Yield of Illumina sequence reads and number and percentage of reads that could be aligned against the *Mitragyna speciosa* reference genome sequence.

A)

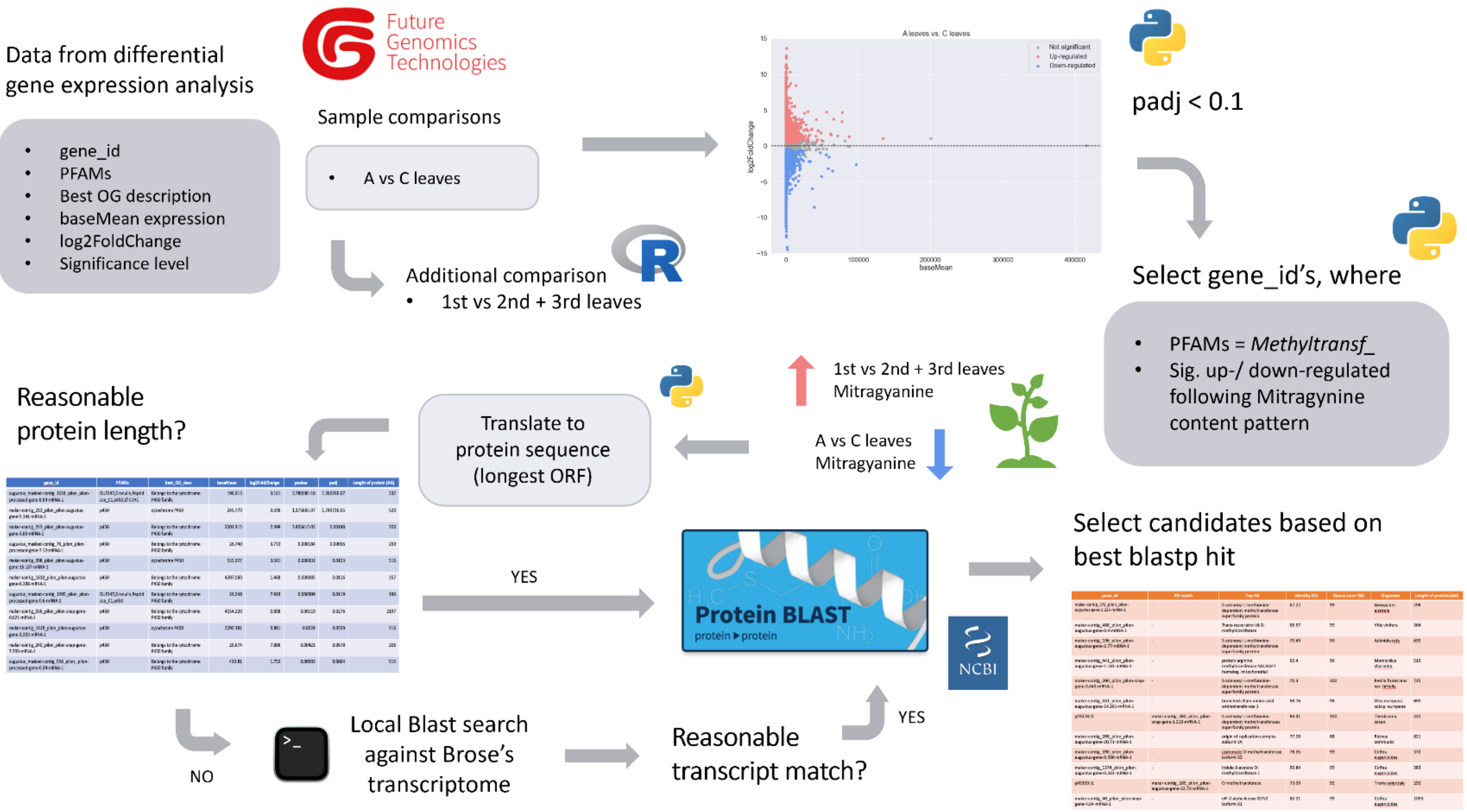

B)

| Candidate number | Short name | Plasmid name | AA_seq | Selection strategy | Family |
|---|---|---|---|---|---|
| 1 | MspOMT_FG177-2-119 | pRS414U-P1_GAL1-MspOMT_FG177-2-119-tADH1 | MDSSATKAKTSYDEEGEHCSYAMQLATSAALPNVLNAIIRLNVLEIIARAGPGAQLSPSEIAAHLHTQNSNAAVMLDRMLRLLASYSVLTCSVVGGGGGGGGDDKDDQHQRVYGLAPVAKYFLQNEDGGSLRPLLDLFQGKVLTDIWCELEPAVLEGGTAFNRKHGIHIFEYTGRDQKFNESFNKAMIDHTFIVVKNVLQFYRGFEHLKTLVDIGGGLGVTLGMITSKYPNIKGINFDLPHVVQNAPTYPGVEHIGGDMFKSIPKGDAVFMKWILHDWDDEHCLKLLKNCYKALPDNGKVIAVDAILPTIPDNSASSKSNIQMDLFTLASYFPGGKERTEQEFLALANEAGFKGIRLECLVCNMWVMEFYK* | Homology to MIA OMTs | Methyltransf_2 |

| | | | | | |
|---|---|---|---|---|---|
| 2 | MspOMT_FG177-2-48 | pRS414U-P1_GAL1-MspOMT_FG177-2-48-tADH1 | MASSILHDWDDEHCLKLLKTCYKALPDNGSMDSSATQAKISYDEEEEHCSYALQLATSAALPVVLNAIIKLNVLEIIAKAGPGAQLSPSEIAAHLPTENSGAAVMLDRMLRLLASYSVLTCSVVGGGDKDNQHQRAYGLAPVAKYELEPAVLKGGTAFNRTHGTHIFQYTGSDQKFNETFNKAMIDHTFIVGRKVLQFYKGFEHLKTLVDVGGGLGVNLSMITSKYPSIKGINFDLPHVVQNAPAYPGVEHVGGDMFESIPKGDAIFMKWILHDWDDDHCLTLLKNCYKALPDNGKVIAVDAILPMVPDNSASCKSNCQVDLFTLALYFPGGKERTEQEFLALAKEAGFKGIRLECFACNMWVIEFYK* | Homology to MIA OMTs | Methyltransf_2 |
| 3 | MspOMT_FG488-0-4 | pRS414U-P1_GAL1-MspOMT_FG488-0-4-tADH1 | MDLARKSDADKLLEAQAHIWNHTFNFINSMCLKCAIQLGIPDIIKKHGRPMTLEELTDALPINNAKSPFIYRLMRILIHSGFFIKAKVTDTDVEKGYMLTSASELLLANDPFSMKTFLLAMLDPILTAPWQHFSQWFQSNDENPFYTCHGRAFWELAGQEPRLNQFFNEAMANDARLVSSIVIKHCKHVFSGLNSLLDVGGGTGTFGKAIAAAFPDLKCTVLDLPHVVDGLDVGGKNLAYIGGDMFEDIPPADAVLLKWILHDWTDEECVRILEKCKQAIPSKENGGKVIIIDMVRNDQQKEADDDARKSIETQLFFDILMMVDTKGRERNEKEWAKLFYEAGFKDYKITQVVGLRSLIEVYYY* | Homology to MIA OMTs | Methyltransf_2 |
| 4 | MspOMT_FG491-0-134 | pRS414U-P1_GAL1-MspOMT_FG491-0-134-tADH1 | MDLARNGDQAGELLQAQAHIWNHIFNFINSMSLKCAVQLDIPDIIHKHGQPMTLAQLIHALPISSAKAPFVHRLMRILIHSGFFVKAKIPGNELEEGYELTSASKLLLKSNPFSVTPFLLAMLDPILTDPWHHFSQWFQCEDESPFCTCHGKAFWKLAGEQPRLNQFFNEGMGSDARLISSLVMKNCKHVFEGLNSLVDVGGGTGTLAKAIADAFPDLKCTVLDLPHVVAGLDGSSKNLAYVGGDMFEAIPPTDAVLLKWILHDWSDEECVQILKKCKEAIPSKGKGGKVIIIDMLVKDQHTGDDKDEAIETQLFFDMLMMVLVKGRERNQKDWAKLFFEAGFNDYRITPVLGLRSLVEVYYY* | Homology to MIA OMTs | Methyltransf_2 |
| 5 | MspOMT_FG491-0-145 | pRS414U-P1_GAL1-MspOMT_FG491-0-145-tADH1 | MDLARNGDRAGELLQAQAHIWNHIFNYINSMSLKCAVQLGIPDIIEKHGQPMTLAQLCHALSISNTKASFLYRLMRVLIHSGFFIKAKIPGNELEEGYELTSASKLLLKSNPFSVTPFLLAMLAPISTDLWHHFSKWFQSDDETPFHTCHGTSGWEMASQEPWINQSFNEAMASDARLLSSILIEHCGHVFSGLNSLVDVGGGTGTLAKAIADAFPDLKCTVLDLPYVVDGLDGGRKNLAFIGGDIFEAVPPADAVLLKWTLHDWSDEDCVRILEKCKQAIPSKENGGKVIIIDMVINDQQKGADDGEHKSIETQLFFDMLMMVDTKGRERNEKEWAEIFSAAGFKDYKITHALGLRSLIEIYY* | Homology to MIA OMTs | Methyltransf_2 |

| | | | | | |
|---|---|---|---|---|---|
| 6 | MspOMT_FG776-65-39 | pRS414U-P1_GAL1-MspOMT_FG776-65-39-tADH1 | MGSAEEEASLFAMQLASASVLPMVLKSAIELDLLELIAKAGPGAYVSPSKLAAQLPTHNPQAPVMLDRILRLLATYSVLNCKLKNLPDGGVERLYGLAPVCKFLTKNADGVSMAPLLLMNQDKVLMESWYHLKDAVIDGGIPFNKAYGMTAFEYHGTDPRFNKVFNQGMSNHSTITMKKILEVYKGFEGVKTVVDVGGGTGATLNMIIDKYPTIKGINFELPHVVEDAPFYPGVEHVGGDMFVSVPEGDAIFMKWICHDWSDDHCRKLLKNCYHALPEGGKVILAECILPDAPDTSLATQNVVHIDVIMLAHNPGGKERTEKEFEALAKGAGFKEFRKICSAVNTWIMELCK* | Homology to MIA OMTs | Methyltransf_2 |
| 7 | MspOMT_FG84-23-157 | pRS414U-P1_GAL1-MspOMT_FG84-23-157-tADH1 | MDLARNGDQAGELLQAQAQIWNHIFNFINSMSLKCAVQLDIPDIIHKHGQPMTLAQLIHALPISSAKAPFVYRLMRILIHSGFFLKAKIPGNELEEGYELTSASKLLLKSNPFSVTPFLLSMLDPILTDPWHQFSQWFQCDDETPFYTCHQKAFWKLAGEHPRVNQFFNEGMASDARLISSLVMKHCKHVFEGLNSLVDVGGGTGTLAMTIADAFPDLKCTVLDLPHVVAGLDGSSKNLAYVGGDMFEAIPPTDAVLLKWILHDWSDEECVQILKKCKEAIPSKGKGGKVIIIDMLVKDQQRGDDKDEAIETQLFFDMLMMVLVKGRERNQKDWAKLFFEAGFNDYKITPVLGLRSLIEVYYY* | Homology to MIA OMTs | Methyltransf_2 |
| 8 | MspOMT_FG93-2-61 | pRS414U-P1_GAL1-MspOMT_FG93-2-61-tADH1 | MEISENAQLKSASPSSIENGQEGEKQDHFSHVMQLVTSASLPMVLYTAVKLNLFEIIAKAGPGAKLSPSEIACRLPTTKNPDAASMLDRMLRLLSSHSLLTYDVVEVAGGAETDIGYERVYGLSPVGEYLIPDEEGNSVAPLVELLLDKVLINTWYELGNAVLEGGIPFNRVHGTHAFDYPSRNPRYNELFNNGMVGPTAITMKELLQQYKGFELLQTLVDVGGGLGITLHKIIAKYPSIRGINFDLPHVIEDAPPYPGMEHIGGDMFESVPGGDAIFMKMILHDWSDDHCLKLLKNCYRALPDHGKVIVVDLILPVKPDTNIFVKGIFQADVIMMTQNPGGKERSEADVRALAIGAGFKGIKLECCAGNVGILELYK* | Homology to MIA OMTs | Methyltransf_2 |
| 9 | MspOMT_g30176.t1 | pRS414U-P1_GAL1-MspOMT_g30176.t1-tADH1 | MDTNVERSKLLHGQAEIWKHTFRFADSMALKCAVELGIADIIHSHGHPLTLAEIATNITNSSSPHIPYLSRIMRFLVRNKIFTSTDIPSDHGGGSPSTTLYGLTAASWWLLNDNDGLSLAPFLLMENHPWLVSPWHQLSSCVREGGIAFQKAHGKEIWELASENPEFNTIFNDGMGCTGKITMKAILSGFKDAWNSIESLVDVGGGTGQTIAEIVKVYPHIRGINFDLPHVVAAAPKYDGVCHVGGDMFDSIPTAQAVFMKWIMHDWGDEDCVKILKICRKAIPEKTGKIFIVDVVLKPDGDGLFDSIGMIFDLLMIAHSSGGKERTEPEWKKLLEKGGFPRYNITEIPACFSIIEAYPV* | Homology to MIA OMTs | Methyltransf_2 |

| | | | | | |
|---|---|---|---|---|---|
| 10 | MspOMT_g51987.t1.1.5bbdf93c | pRS414U-P1_GAL1-MspOMT_g51987.t1.1.5bbdf93c-tADH1 | MEKEDCLAKTNSNENSEVVELGGKVEEEEQLFSYAMQLVTSVSLPRVLLAAFRLRVFEIIAEAGPGAQLSASEIAAQVSGKNKDAAAMLDRMLRLLASHSVLSCGARSQGRVYGLAPVAKYFVQSKADGGGTLGSLLALIQDKVFIDSWYQLEDAVREGGVPFDRVYGTHAFEYPGRDPRFNEVFNKAMLDHTTIAINRILERYKGFENLKTLVDVGGGLGVTLNVITTKYPNLKGINFDLPHVIQRAPGYPGVEHIGGDMFESVPEGEAIFMKWILHDWDDGHCLKLLKNCYKALSDNGKVIVVDALVPVIPDTSPCVKTTCQSDLIMMTQNPGGKERSEPEFLALATAAGFKGIRLECYVCNLWVMEFYK* | Homology to MIA OMTs | Methyltransf_2 |
| 11 | MspOMT_FG1374-0-261 | pRS414U-P1_GAL1-MspOMT_FG1374-0-261-tADH1 | MVSAEDNVVVSNMKLEKLLSMKGGKGEASYVNNSQAQAQHARSMLHLLNDTLDVVRINSPEIPFVVADLGCSCGSNTLYIIDVIIKHMIKRYEALGYDEPPEFSAFFSDLPSNDFNTLFQLLPPYGGSMEECLASDSHRSYFAAGVPGSFYRRLFPARSIDVFYSAFSLHWLSQVPDIVLDKGSTAYNKGRIFIHGANEITANAYRKQFQTDLAGFLRSRSLELKRGGSMFLVCLGRTSMDPTDQGGAGLLFGTDFQDAWDDLVQEGLITSEQRDNFNIPVYAPSLQDFKEVVEADGSFVINKLEVFKGGSPLVVNHPDDAAEVGRALANSCRSVSGVLVDAHIGDQLSEELFSRVERRATSHAKEILEQLQFFHIVASLSLP* | Differential expression, FG annotation | Methyltransf_7 |
| 12 | MspOMT_FG151-7-370 | pRS414U-P1_GAL1-MspOMT_FG151-7-370-tADH1 | MDEKEGPYFIFPGGRTMFPDGAVQHIEKLKQYVPIAGGVLRTALDMGCGMQQFLEVDRLLRPGRYLVISGPPVQWPKQDREWVDLQGVARALCYELIVVDGNTVVWKKPVDSCLPNQNEFGLGFCDESDDPNKCVSRTSTVKGEVASGTIPKWPDRLTEAPSRATLVKNGIDVFAADTQRWVRRVAYYKKSLNLKLGTSSTRNVMDMNALFGGFAAALSSDPVWVMNVVPAHKPSTLSVIYDRGLIGMYHDWCEPFSTYPRTYDLIHAASIESLIKDSSSGKMSCTHQTSSRHRLISLPL* | Differential expression, FG annotation | Methyltransf_29 |
| 13 | MspOMT_FG196-2-77 | pRS414U-P1_GAL1-MspOMT_FG196-2-77-tADH1 | MATLQPPVSQLPQKSPAPKLLNKPLFNILLLTIFCSLSYFIGSYTNSIYSISPSPSLIRRDTNCGFEVKNNLNTTVENVQEPLNFEPLHTLSLPMEDTPESHQPFPYCPKNYTNYCPCQDPNKERLFTIDRRFHRERHCPEKGEALRCLVQKPAGYRRPFPWPKSRDYAWFKNVPFKRLTEYKKSQNWVRQEGDKLFFPGGGTSFPKGVKGYIDLLKRVVPLNSGAIRTVLDVGCGVASFGASLMDYNIVTMSIAPRDIHEAQVQFALERGVPALLGVLSTYRLPFPSRCFDMIHCSRCLVQWTDYGGLYLMEIDRLLRPGGYWVLSGPPISWRISYKGWERNASDLEIEQNNLEDLARKLCWKKVRESDPIAVWQKPTNHIHCAQKLKTWKAPKFCSHDFPDHGWYKKMEPCITPLPKVEHVRSTSGGYLEKWPKRLKTAPPRIRSATVEGVTMRTFDEDNKIWKRRVLYYGSVLKFLFKGKYRNIMDMNSGLGGFAAVLSEYPAWVMNVVPYDAKNNTLGIVYERGLIGTYMDWCEPFSTYPRTYDLIHSDGLFSLYMEKCDILDILFEMYRILRPEGAIIIRDHVDVIVKIKEITDRMRWNNHMALLSPGSQQQHHLHNPSPPPPTRPPSPPPPSS* | Differential expression, FG annotation | Methyltransf_29 |

| | | | | | |
|---|---|---|---|---|---|
| 14 | MspOMT_FG240-6-196 | pRS414U-P1_GAL1-MspOMT_FG240-6-196-tADH1 | MDREKKTDKKEEPEEAIEQILFQVSECYVYLMCILKSYYLGSLNHILLGSYMEILSGFYFASGQPRINRLLVLCRADEWNVNKWAWEGGLKVVSKGEECIIRLEDKTTGELYARAFLRDGEPHPVEPVIDSSRYFVLRVEENIGGRGRHAFIGIGFRERPQAYDFQAALHDHMKYLNKKKTAEEMEQQYQKTSLTDYSLKKGETLVLQIKTKSGRSTGSKFFEQGLNNLSLEKKGTQKEPVISIKPPPPPPAPLSPVVSPTKSPVELPSELSLKEASEVKDSIAPKEQSKESASPENQNIQDMPDDDFGDFQAAG* | Differential expression, FG annotation | DUF1681,DUF4201,Methyltransf_11 |
| 15 | MspOMT_FG379-3-225 | pRS414U-P1_GAL1-MspOMT_FG379-3-225-tADH1 | MIVNIHQNPNSWNASILTSNRVFFSIQRSTLIFNSTSTLHTNRFSINASLREDNNQTKQSAAEAGKIRRLVLAQQGRTKLNPKPDREFYAYPRFVTHVDDNFISGLTNLYREKLKPESEILDLMSSWVSHLPPEVKYKKVVGHGLNAQELARNPRLDYFFVKDLNQEQGLQLEDCSFDAVLCTVSVQYLQQPEKVFAEVFRVLRPGGVFIVSFSNRLFYEKAISAWRDGTSYSRVQLVVQYFQCVEGFTHPEVIRKLPTDTRDKQTPFSWIMNLVGLISGSDPFYAVLAYKNFKPTYE* | Differential expression, FG annotation | Methyltransf_11 |
| 16 | MspOMT_FG379-4-619 | pRS414U-P1_GAL1-MspOMT_FG379-4-619-tADH1 | MVRGLNAQTARAFPWTPSSFQPLWSHQRFDFSVSLPSFPGVCSTSSLKVSSCREHDLRVLRQGRNVHILERRQTETHSFQESPNELTCVMKFGGSSVSSAERMREVADLICSFPEERPVIVLSAMGKTTNALLQAGEKAVSCGVSNVNELKELDFIKELHLRTLHELGLESFIISRHLDELEQLLKGIAMMKELTLRTRDYLVSFGECMSTRIFAAYLNKIGVKARQYDAFEIGFITTDDFTNADILEATYPAVAKRLHDDWISDPAIPIVTGFLGKGWRTCAVTTLGRGGSDLTATTIGKALGLREIQVWKDVDGVLTCDPNIYPRAEPVPFLTFEEAAELAYFGAQVLHPQSMRPAREADIPVRVKNSYNPMASGTLITRSRDMSEAVLTSIVLKRNVTMLDIVSTRMVGQFGFLAKELDRVVEELEKIAVVNLLQRRSIISLIGNVQRSSLILEKAFHVLRTNGVNVQMISQGASKLLLGSRVGGDLEMVGDAVGAFV* | Differential expression, FG annotation | AA_kinase,Methyltransf_7 |
| 17 | MspOMT_FG390-9-290 | pRS414U-P1_GAL1-MspOMT_FG390-9-290-tADH1 | MEIIQILHMNGGEGETSYAKNSVVQRKIISFGNASIEEAVNDILCKNFPESMGIADLGCSSGPNTLTLISEVIEIVNSKSHKTGNPPPEIRVCLNDLPGNDFNTIFMSLPAFYQKQEEENGKGLRNNCFVSCVAGSFYGRLFPKKSLHFVHSSSSLHWLSQVPPLLDVDAIPPLNLGKIYISKTSPPSVMYAYLLQFQKDFSLFLKSRSQEMVPGGRMVLSFLGRTSADPTTEDSCYHWELLATALTCLASEGRVREEKINSFNAPYYAPSVEEVSIEVEKEGSFLINSLKAFEVEWDDGFQSQENKLQPSSRGRRVAKTIRAVVESMLERHFGRDILDDLFVKYAELVDDYSSRTTPKYTNLVISATRKGL* | Differential expression, FG annotation | ENOD93,GH3,Methyltransf_7 |

| | | | | | |
|---|---|---|---|---|---|
| 18 | MspOMT_FG390-9-449 | pRS414U-P1_GAL1-MspOMT_FG390-9-449-tADH1 | MAIAKFRRLAKRSYKSYGFCVKLTAIVILGLCFVFVWSVFSPSKFSVTSQRETFDDFAEPVSANGKATDSGVHFNKNKFKNGKESGPEKKVQSLEGKDKRKVKGSPPLKPGNMHKDQKQGSKGKKQGVNGSKPVKEVGGQKIEETERSEGEDLQEEKQDEDVDNREEETDNAEVNVNEEGDGNQGDSDLADALDLDQEDGEKVEDDGGLSTNSKKKKRNLGPLFDPKAHYVWKLCSTRSKHNYIPCIDIESASGRLQSYRHHERSCPKTSVMCLVPLPRDGYGTPVRWPESKVKILYKNVAHPKLATFVKSHKWVVEDGEYLTFLQNQSVLKGGIQHYLESIEEMVPDIEWGKNIRVVLDVGCTDSSFVASLLQRNVLALTLGLKDDLVDLAQLALERGFPAVVSPLAMRRLPFPSGVFDAVHCGECSISWHANGGKLLLEMNRILRPGGYFILSSKHDSIEIEEAMSKVTASVCWNILADKADEVSDIGVKIYQKPESNDIYELRRKKVPPLCKENENPDAAWYASIKTCLHTIPSSIEERGTEWPEGWPKRLETFPDWINNRERLIAESEHWKAIVNSSYLTGMGIDWLSIRNVMDMKAINGGFAAALSEQKVWVMNVVPVHAPNTLPIIYERGLIGVYHDWCEAFGTYPRSYDLLHADHLFSRLKNRCKQPVVIVVEMDRILRPGGWAIIRDKVEILDPLEGILRSLHWEIRLTFRKDREGILCAQKTDWRP* | Differential expression, FG annotation | Methyltransf_29 |
| 19 | MspOMT_FG488-0-4 | pRS414U-P1_GAL1-MspOMT_FG488-0-4-tADH1 | MDLARKSDADKLLEAQAHIWNHTFNFINSMCLKCAIQLGIPDIIKKHGRPMTLEELTDALPINNAKSPFIYRLMRILIHSGFFIKAKVTDTDVEKGYMLTSASELLLANDPFSMKTFLLAMLDPILTAPWQHFSQWFQSNDENPFYTCHGRAFWELAGQEPRLNQFFNEAMANDARLVSSIVIKHCKHVFSGLNSLLDVGGGTGTFGKAIAAAFPDLKCTVLDLPHVVDGLDVGGKNLAYIGGDMFEDIPPADAVLLKWILHDWTDEECVRILEKCKQAIPSKENGGKVIIDMVRNDQQKEADDDARKSIETQLFFDILMMVDTKGRERNEKEWAKLFYEAGFKDYKITQVVGLRSLIEVYYY* | Differential expression, FG annotation | Dimerisation,Methyltransf_2 |
| 20 | MspOMT_g19278.t1 | pRS414U-P1_GAL1-MspOMT_FGg19278.t1-tADH1 | MASSLLSGAENLKHITGSHFPGSEHQLKYFPKKGPVSSAKSSKFSTLAPRCSLSSSRPVSQPRFIQHKQEAFWFYRFLSIVYDHIINPGHWTEDMRDEALEPADLYSRHLTVVDVGGGTGFTTLGIVQHVDAKNVTILDQSPHQLAKAKQKEPLKECKIIEGDAEDLPFRTDYADRYISAGSIEYWPDPQRGIKEAYRVLKIGGKACVIGPVYPTFWLSRFFADVWMLFPKEEEYIEWFQKAGFKDVKLKRIGPKWYRGVRRHGLIMGCSVTGVKPMSGDSPLQLGPKQEDVKKPVNRFVFVLRFILGAMAAAYFVLVPIYMWLKDQIIPKGMPI* | Differential expression, FG annotation | CTP_transf_like,Methyltransf_11,Methyltransf_25,Ubie_methyltran |

| | | | | | |
|---|---|---|---|---|---|
| 21 | MspOMT_g21190.t1 | pRS414U-P1_GAL1-MspOMT_FGg21190.t1-tADH1 | MGSEGLLRSPELRQYLLETSVYPREAEILKELRAVTATHPRGIMATDADAGQMLSLLLEITNAKKTIEIGVFTGYSLLLTALTIPEDGKTIAIDPDQEAYDIGLPFINKAGVEHKINFINSVAHPVLDKLLEDPNEHDSFDFAFVDADKVSYSKYHEKLLKLVKVGGIIVYDNTLWMGTVAEAEDSVPETLRESRHYIKELNKLLAADARYLLETSVYPREPEILKELRAVTATHPQGILATDADEGQMLALLLEIINAKKTIEIGVFTGYSLLLTALTIPKEGKIIAIDPDQEAFNIGLPFIKKAGVEHKINFINSVAHPVLDKLLEDPNEHDSFDFAYVDADKVNYAKYHEKLLKLVKVGGIIVYDNTLWMGTVAEAEDSVPEEFGESRHYIMELNKLLAADARVHICQLPSGDGMTICRRLH* | Differential expression, FG annotation | S-adenosyl-L-methionine-dependent methyltransferases superfamily protein isoform 2 |
| 22 | MspOMT_g14031.t1 | MspOMT_g14031.t1 in pTwist Kan High Copy | MDSSATQAKISYDEEEEHCSYALQLATSAALPVVLNSIIKLNVLDIIAKAGPGAQLSPSEIAAHLSPENSGAAVMLDRMLRLLASYSVLTCSVVGGGDKDNQHQRAYGLAPVAKYFLRDEDGGSLGSLLDLLQDKVLVDIWYELEPAVLKGGTAFNRTHGTHIFQYTGSDQKFNETFNKAMIDHTFIVVRKVLQFYKGFGHLKTLVDVGGGLGVNLGLITSKYPSIKGINFDLPHVVQNAPAYPGVEHVGGDMFESIPKGDAIFMKWILHDWDDEHCLKLLKNCYKALPDNGKVIAVDAILPMVPDNSASCKSNCQVDLFTLALYFPGGKERTEQEFLALAKEAGFKGIRLECFACNMWVIEFYK | Differential expression, Brose annotation | O-methyltransferase domain |
| 23 | MspOMT_g21161.t2 | MspOMT_g21161.t2 in pTwist Kan High Copy | MGSEGLLRSPELRQYLLETSVYPREAEILKELRAVTATHPRGIMATDADAGQMLSLLLEITNAKKTIEIGVFTGYSLLLTALTIPEDGKTIAIDPDQEAYDIGLPFIKKAGVEHKINFINSVAHPVLDKLLEDQPNEHDSFDFAFVDADKVSYSKYHEKLLKLVKVGGIIVYDNTLWMGTVAEAEDSVPESLRESRHYIKELNKSLAADARVHICQLPSGDGMTICRRLQ | Differential expression, Brose annotation | O-methyltransferase |
| 24 | MspOMT_g32765.t1.1.5bbdf938 | MspOMT_g32765.t1.1.5bbdf938 in pTwist Kan High Copy | MAGIRLPPEDSDASQARPPAAADLISDDDRSVAADSWSIKSDYGSTLDDEQRHADASEALSAANFRAASDYSSDKEEPDAEAVQSMLGFQSYWDAQYADELANFRQHGHVGEVWFGADVMEIVASWTKSLCSEFSQDHLSNHANDSNYESVVQHEKEFSNWSVLDVGTGNGLLLQELAKQGFSDLTGTDYSEGAIDLARSLAERDGFTSIKFLVDDILETKLDRKFQLVIDKGTLDAIGLHPDGPVKRIMYWSSVSRLVAPGGLLVSFSLSLAI | Differential expression, Brose annotation | Methyltransferase domain |
| 25 | MspOMT_g40632.t1 | MspOMT_g40632.t1 in pTwist Kan High Copy | MATLENSAELLRAQAHVWSQTFNFKNSASLKCAVQLGIADVIQKHGKPITLSKLAAALPINPSKANYIYRLMRILVNAGFFAQEKEGYSLTSAGRLLLSDEPVNARAFILVVLDPDTIKPWNFLSDWFQNDDPSPFDTAHGKNFWDFTAEEPKFSKLFNEGMVSDSHLITMVLVTECKYVFEGLTSLVDVGGGTGTVARSIAEAFPNLKCTVFDLPHVVANQEGTLNLDFVAGNMLEKVPPANAILLKWILHDWSDEACVKILKNCKKAIPGRDIGGKVIIIDMVMESQLEDDESTEAQICLDMQMLVLYRAKERSEKEWAKLFSDAGFPSYKVFPALGTRAIIEVYP | Homology to MIA OMTs | Methyltransf_2 |

| | | | | | |
|---|---|---|---|---|---|
| 26 | MspOMT_g53423.t1 | MspOMT_g53423.t1 in pTwist Kan High Copy | MMKTILKTEALQEYILNTSTYPREHEQLKGLREETA<br>RIYGARAFMSVPPDEGLFLSMLLKTMNAKKTLEFG<br>VFTGYSLLTTALALPSDGQIVAIDPDQGAFEVGLEFI<br>KKAGVENKINFIKSDAISALNEMLNNETSEATFDFV<br>FVDADKPNYIHYHERLLKLVKIGGIIAYDNTLYGGYV<br>VSEEIEMHERSTVNRKAIIQFNNQIASDPRVEISQIPI<br>GDGVTLCRRIA | Differential expression, Brose annotation | O-methyltransferase |
| 27 | MspOMT_g53883.t1 | MspOMT_g53883.t1 in pTwist Kan High Copy | MATLENSAELLRAQAHVWSQTFNFKNSASLKCAV<br>QLGIADAIQKHGKPITLSELAAALPINPSKANYIYRL<br>MRILVNAGFFAQEKEGYSLTAAGRLLLSDEPVNAR<br>AFILVVLDPDTMKPWNFLSDWFQNDDPSPFDTAH<br>GKNFWDFTAEEPKFSKLFNEGMVSDSHLITMVLVT<br>ECKYVFEGLTSLVDVGGGTGTVARSIAEAFPNLKC<br>TVFDLPHVVANQEGTLNLDFVAGNMLEKVPPANAI<br>LLKWILHDWSDEDCVKILKNCKKAIPGRDIGGKVIII<br>DMVMESQLEGDESTEAQICLDMQMLVLYRAKERS<br>EKEWAKLFSDAGFPSYKVFPALGTRAIIEVYP | Homology to MIA OMTs | Methyltransf_2 |
| 28 | MspOMT_g58593.t1.4.5bbdf934 | MspOMT_g58593.t1.4.5bbdf934 in pTwist Kan High Copy | MSTCAAFFVSPASLATISQRCSYTPPQLLTTLRCGL<br>LLPAQLDKQEAEVEHHYSRNSLRYWDKFYKLHKN<br>KFFKDRHYLEKDWGQYFCDSDAHDTNSPQGKVVL<br>EVGCGAGNTIFPLIAAYPKLFVHACDFSRQAVSLVK<br>SHANFNNDRIDVFVCDVAKDDLCINIKSSSVDVVTLI<br>FMLSAVAPSKMPSVLQNLKRILKPNGHILLRDYAFG<br>DSAQVKLQNKNQKIGENFCFRGDGTCSFYFSEDF<br>LSSLFERAGFNIVDINTCSRQIENHSKNIIMSRRWIR<br>AIFSQI | Differential expression, Brose annotation | Methyltransferase domain |
| 29 | MspOMT_g68830.t1.2.5bbdf93e.2.5bbe1a06 | MspOMT_g68830.t1.2.5bbdf93e.2 in pTwist Kan High Copy | MGGSLDAIRRLLRGRGGIEKLIMMDTSYDMVKLCK<br>NAEASISNETIETSYVVGDEEFLPIKESSVDLIISSLG<br>LHWTNDLPGAMIQCRLALKPDGLFLAAIFGGETLK<br>ELRIACTVAQMEREGGISPRLSPLAQVRDAGNLLT<br>RAGFTLPGVDFDEYIVRYKNPLELIEHLRAMGETNA<br>LLQSSKVRLIHLFGYMIFINTSPPLSFIKLFFSNKLDI<br>RLLFNILLLYPCCLNNMK | Differential expression, Brose annotation | Methyltransferase domain |
| 30 | MspOMT_g71617.t1 | MspOMT_g71617.t1 in pTwist Kan High Copy | MIEGAIVEGLDTKSLMSSSNAFNIVDLGCSVGPNTF<br>ITMQNVVDAVEKKYHSQGHGSNKLQFQVFFNDHV<br>SNDFNTLFASLPIEGQYFAAGVPGSFYDQLFPSSSI<br>HFAHSSYALHWLSKLPEGLLDKNSPAWNQGRIHY<br>TGASDEVMNAYASQFAKDMETFLSARATETVPGG<br>LVVIIVPGLPEGVHYSELPMGILFNFLGSCLLDMAN<br>ETIINGAEIDSFNLPIYAASVQEMTKLVEKNGSFSIQ<br>RMEMADPRSKLPNSINAQDLIMHLRAGMEGLFTKH<br>FGSKNVEEMFNRTFQKIPEISRWLESGYKKGTQLF<br>AVLKRK | Differential expression, Brose annotation | SAM dependent carboxyl methyltransferase |

**Supplementary Figure 5.** Candidate MsOMT gene candidates screened and selection strategy. **A)** Bioinformatic workflow for selecting candidate genes from the transcriptome dataset generated in this study ("FG annotation"). **B)** Table of candidate MsOMTs genes screened in MIA-KM-2.

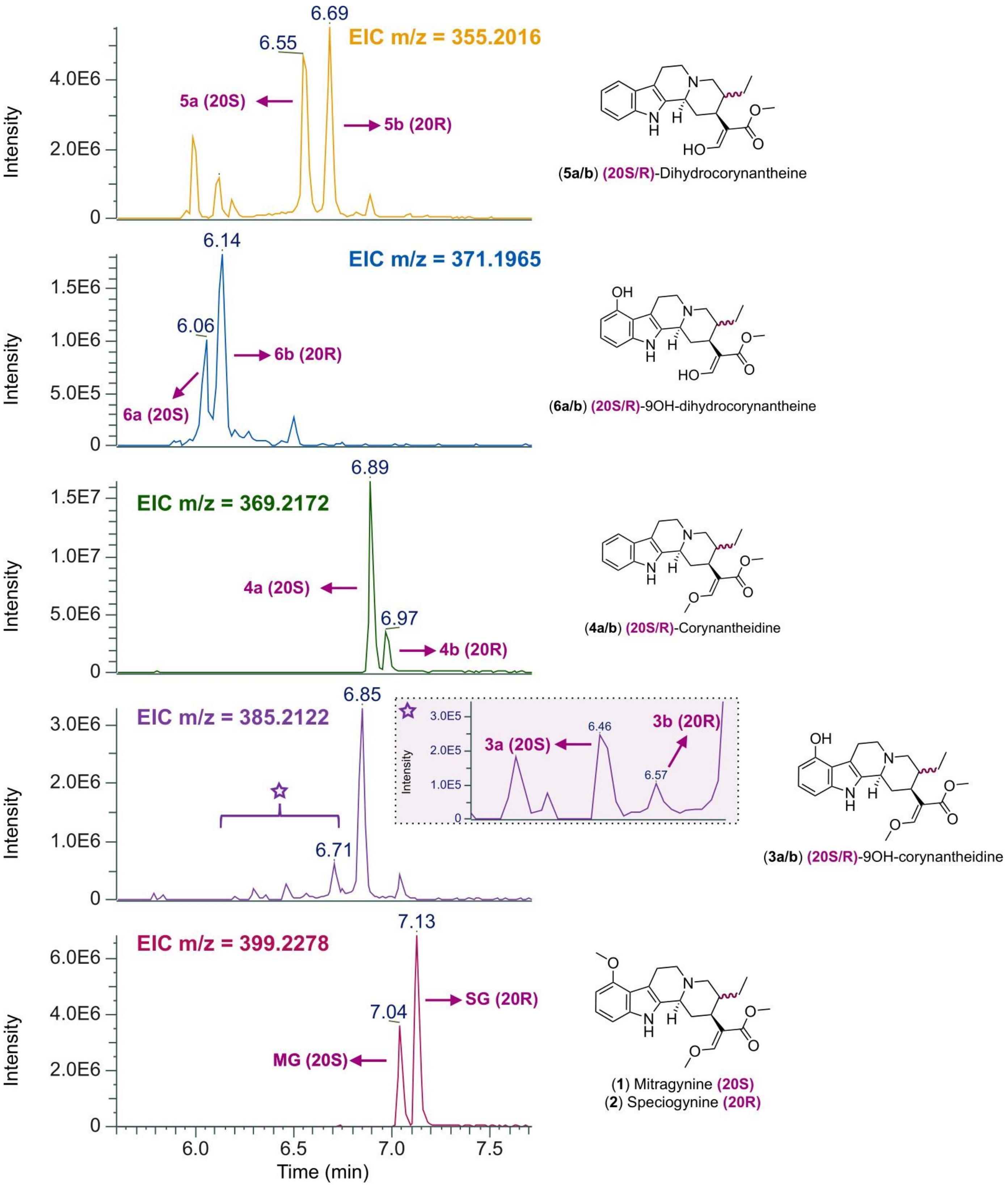

**Supplementary Figure 6.** Stereochemistry of different pathway intermediates generated by MIA-KM-5. (EIC for pathway intermediates downstream of MsDCS1 in MIA-KM-5). Stereochemistry of the different peaks were either determined using authentic analytical standards (MG, SG and 20R-corynantheidine) or by deduction from data obtained with MIA-KM-7 (CpDCS) producing exclusively the 20R stereochemistry.

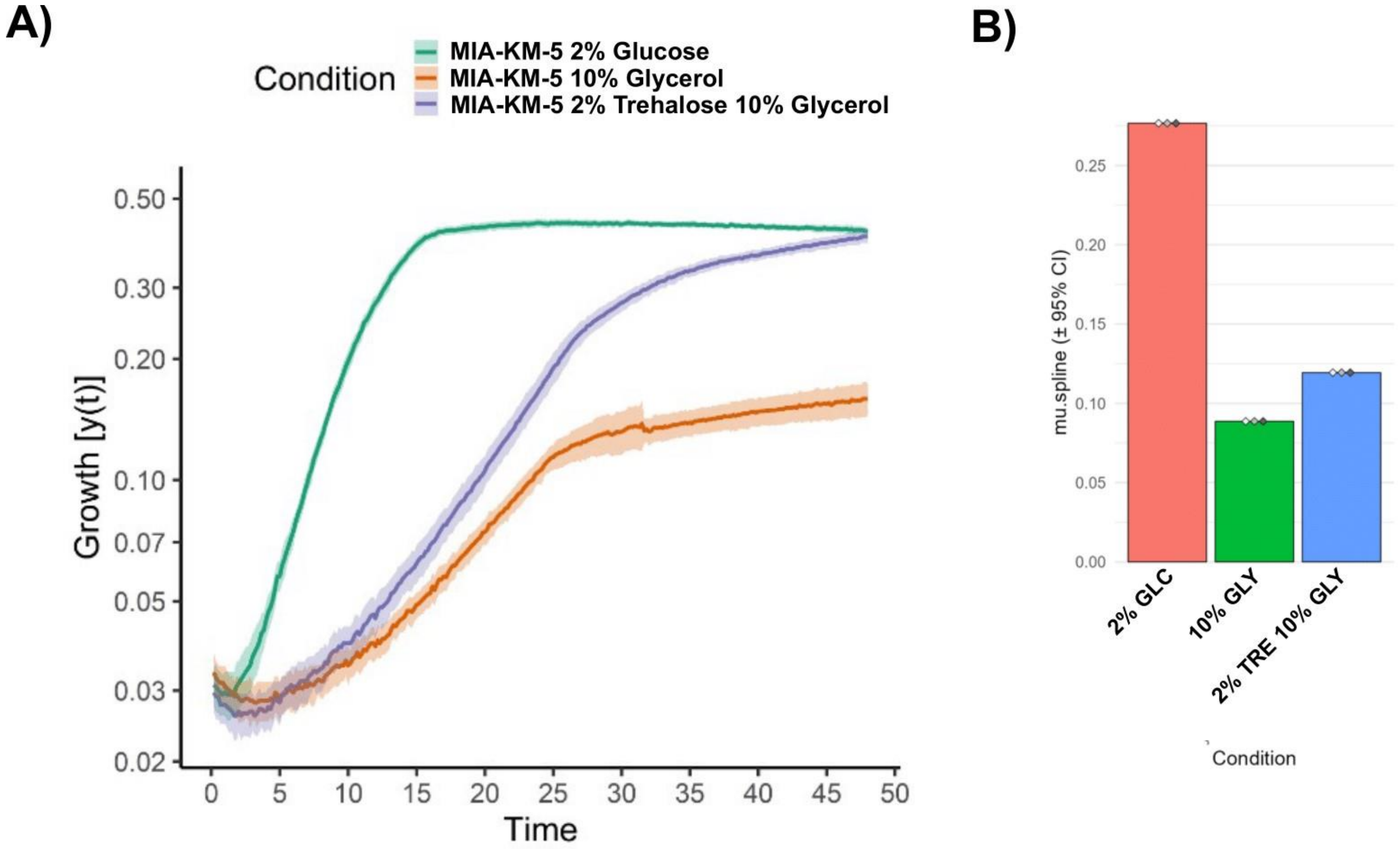

**Supplementary Figure 7.** Growth characterization of MIA-KM-5 on best performing carbon source mix 2% trehalose (TRE) and 10% glycerol (GLY) compared to glucose (GLC). **A)** Growth curve of MIA-KM-5 on different carbon sources in 96 well plate in a plate reader at 25°C (3xSC, Pbov 1%, 3mM tryptophan, pH 5.5). **B)** Maximum growth rate calculated for each condition ($\mu_{max}$ in $h^{-1}$). Data was plotted and processed using QurvE (Wirth et al., 2023).

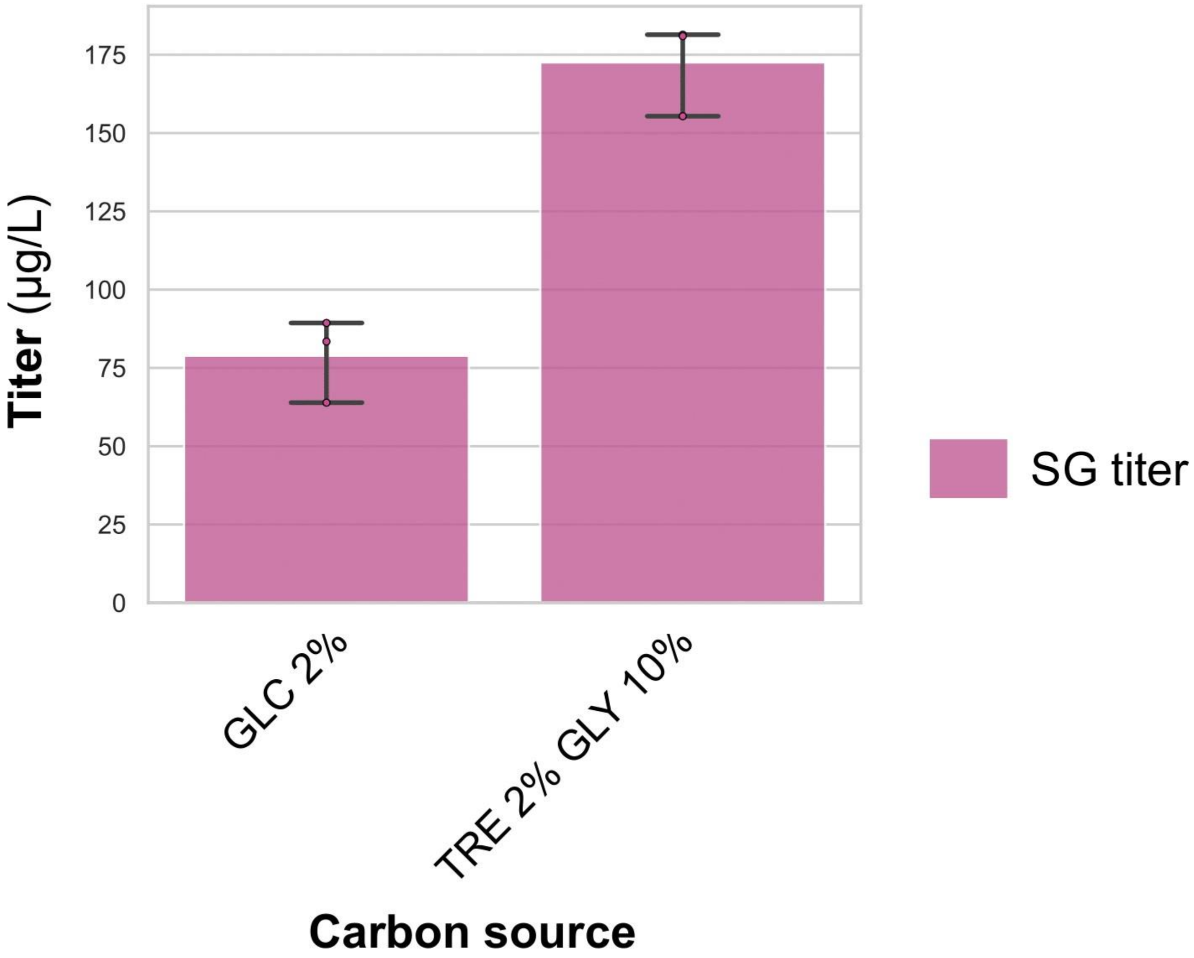

**Supplementary Figure 8.** Production of speciogynine by MIA-KM-7 (CpDCS) in optimized DW96 batch conditions. GLC = Glucose, TRE = Trehalose and GLY = Glycerol. Cultivation at 25°C (3xSC, Pbov 1%, 3mM tryptophan, pH 5.5).

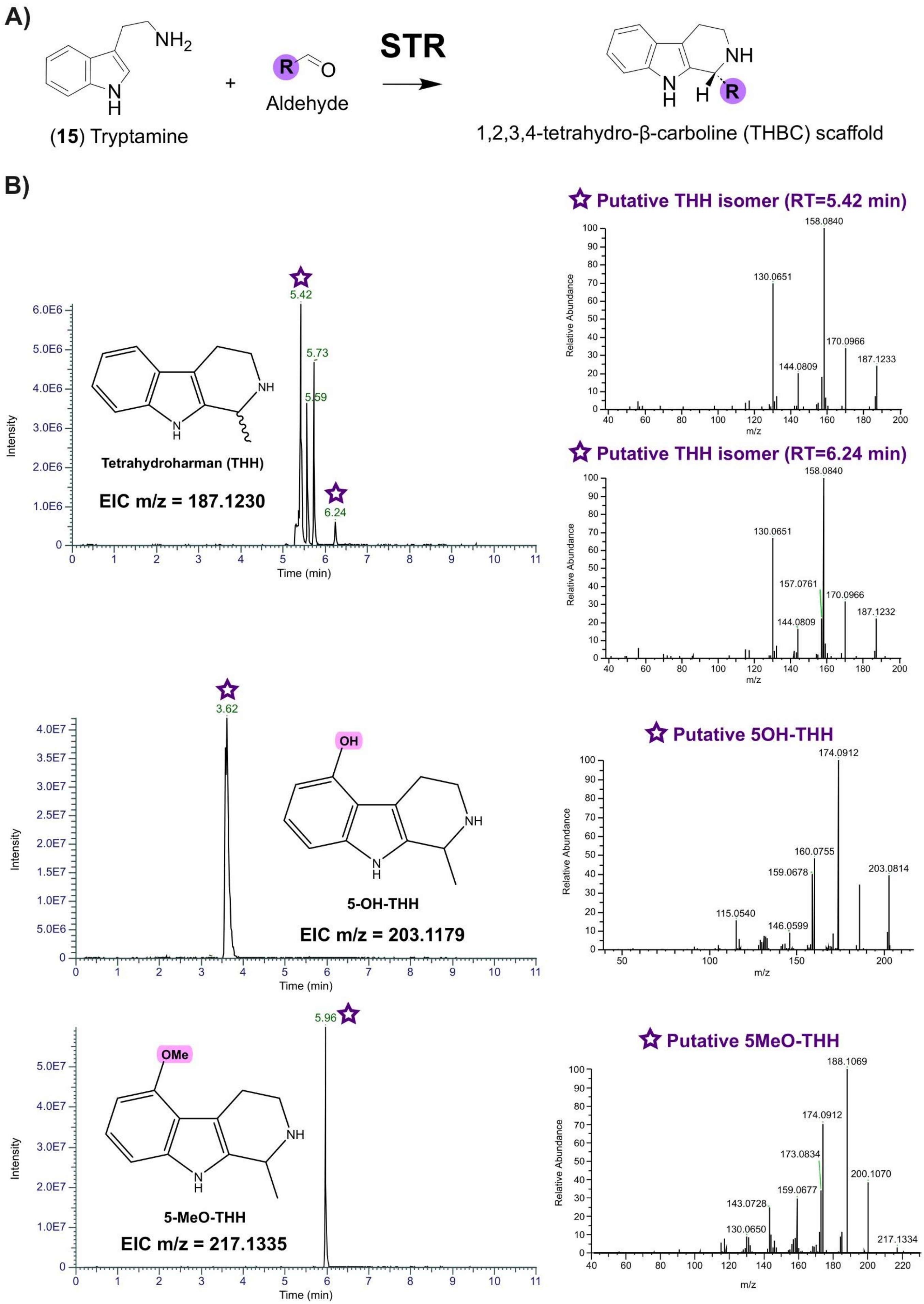

A)
NH2
(15) Tryptamine
+
R
O
Aldehyde
STR
NH
N
H
H
R
1,2,3,4-tetrahydro-β-carboline (THBC) scaffold
B)
Putative THH isomer (RT=5.42 min)
158.0840
130.0651
170.0966
187.1233
144.0809
Relative Abundance
m/z
5.42
5.73
5.59
6.24
Tetrahydroharman (THH)
EIC m/z = 187.1230
Intensity
Time (min)
Putative THH isomer (RT=6.24 min)
158.0840
130.0651
157.0761
170.0966
187.1232
144.0809
3.62
OH
5-OH-THH
EIC m/z = 203.1179
Putative 5OH-THH
174.0912
160.0755
159.0678
203.0814
115.0540
146.0599
5.96
OMe
5-MeO-THH
EIC m/z = 217.1335
Putative 5MeO-THH
188.1069
174.0912
173.0834
200.1070
143.0728
159.0677
130.0650
217.1334

**Supplementary Figure 9.** STR-derived shunt products identified in kratom yeast strains using untargeted metabolomics data processing pipeline. **A)** Strictosidine synthase (STR) can condense tryptamine with short chain aldehydes such as acetaldehyde produced by yeast. **B)** EIC and MS2 spectra for tetrahydroharman (THH), putative 5OH-THH and putative 5MeO-THH. Purple color indicates a putative identification using exact mass and MS2 spectra computation for structure prediction with SIRIUS (Dührkop et al., 2019). The star symbol indicates from which peak MS2 had been acquired.

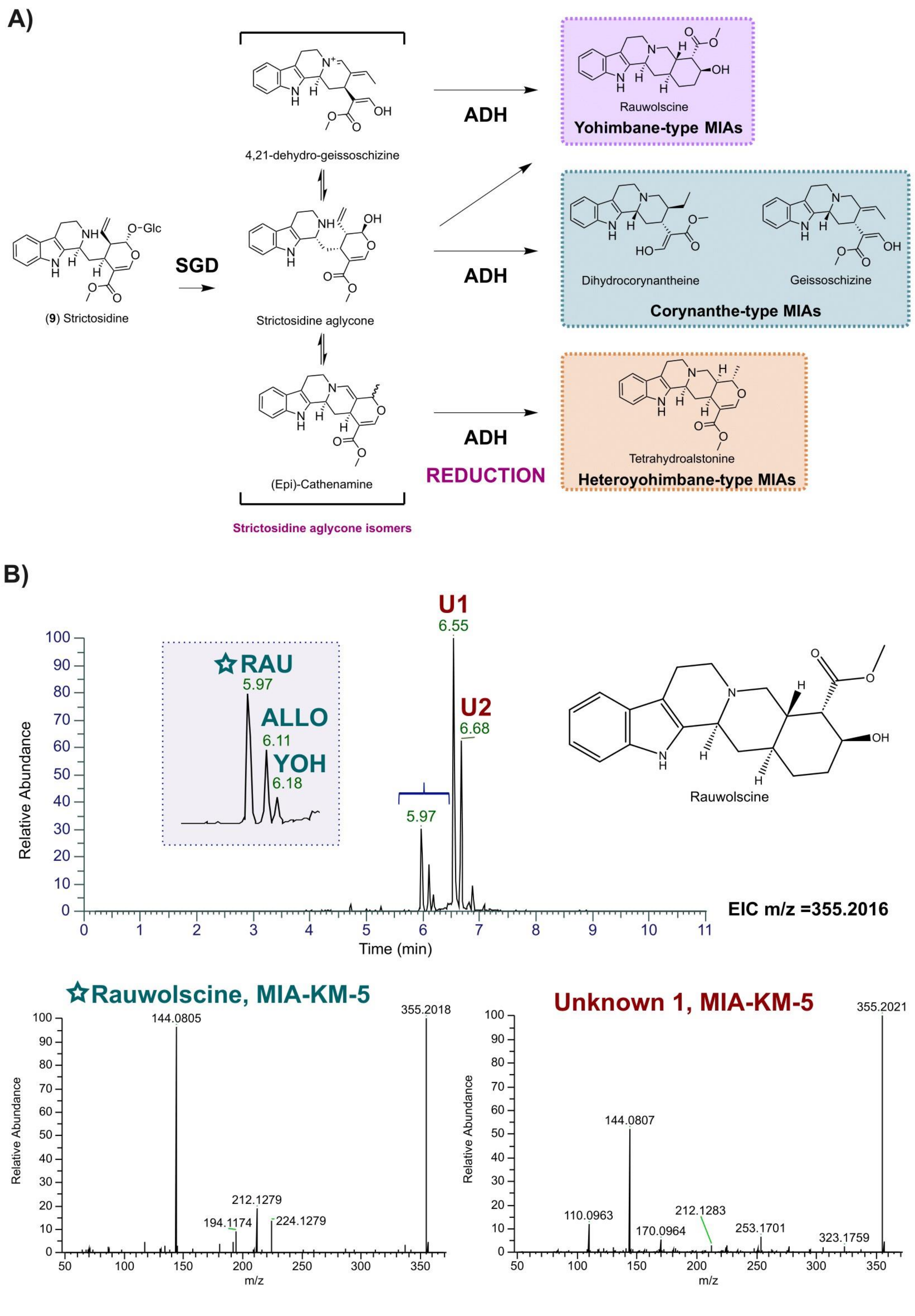
A)
(9) Strictosidine
SGD
4,21-dehydro-geissoschizine
Strictosidine aglycone
(Epi)-Cathenamine
Strictosidine aglycone isomers
ADH
ADH
ADH
REDUCTION
Rauwolscine
Yohimbane-type MIAs
Dihydrocorynantheine
Geissoschizine
Corynanthe-type MIAs
Tetrahydroalstonine
Heteroyohimbane-type MIAs
B)
U1
6.55
U2
6.68
☆RAU
5.97
ALLO
6.11
YOH
6.18
5.97
Relative Abundance
Time (min)
EIC m/z =355.2016
Rauwolscine
☆Rauwolscine, MIA-KM-5
144.0805
355.2018
212.1279
194.1174
224.1279
Unknown 1, MIA-KM-5
355.2021
144.0807
110.0963
212.1283
170.0964
253.1701
323.1759
m/z

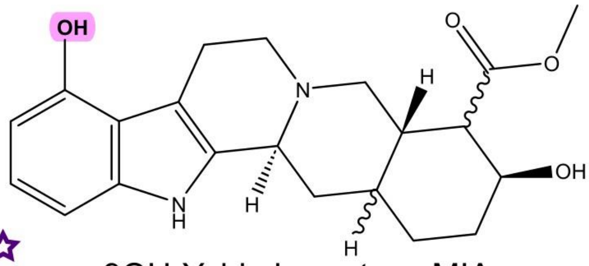

9OH-Yohimbane-type MIA

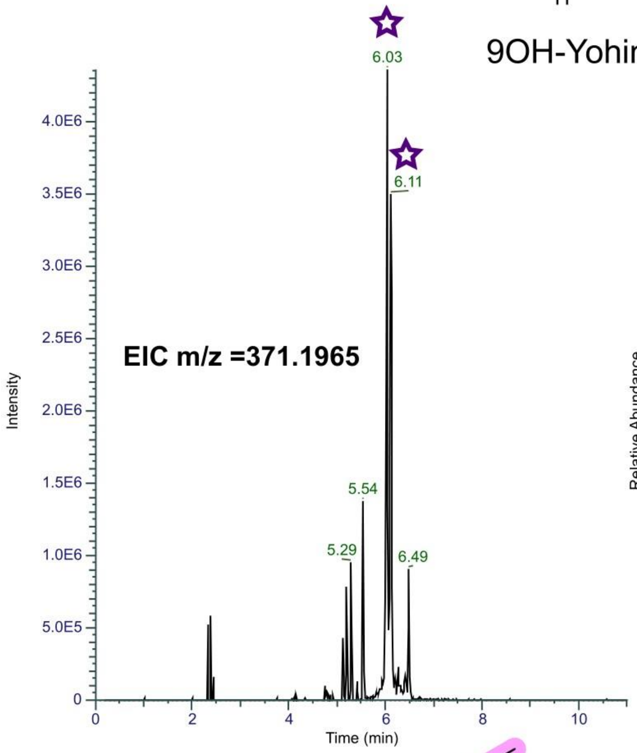

**☆ Putative 9OH-yohimbane-type MIA**

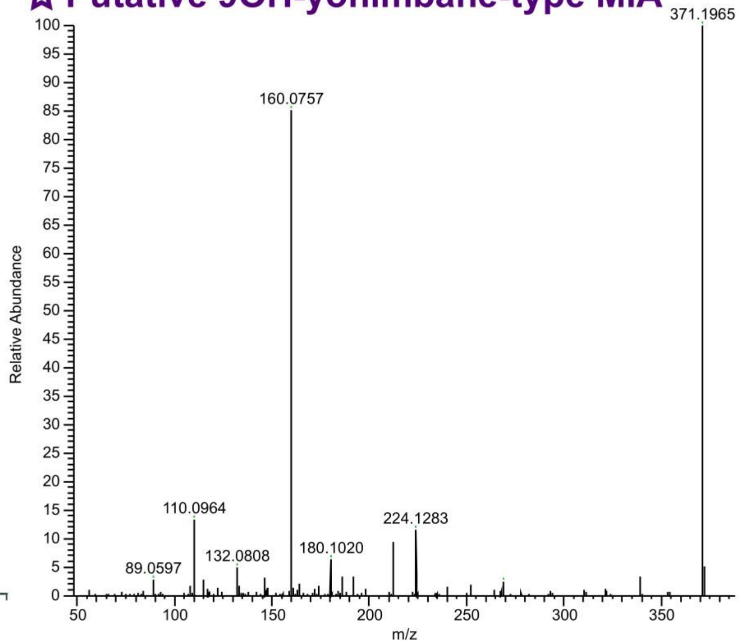

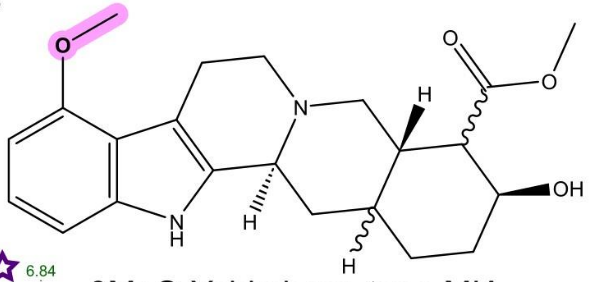

9MeO-Yohimbane-type MIA

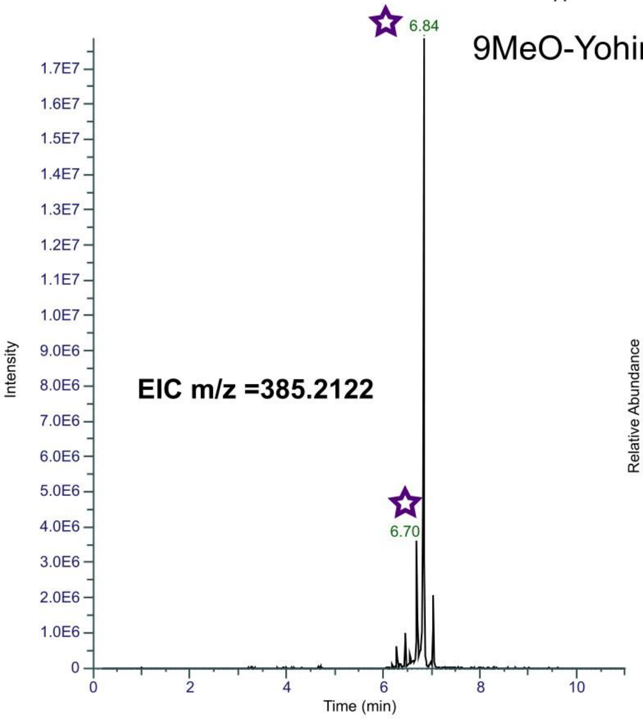

**☆ Putative 9MeO-yohimbane-type MIA**

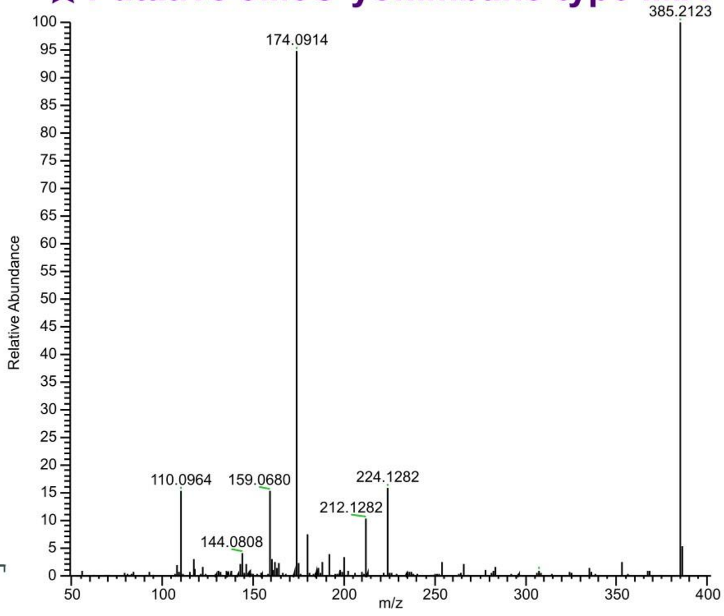

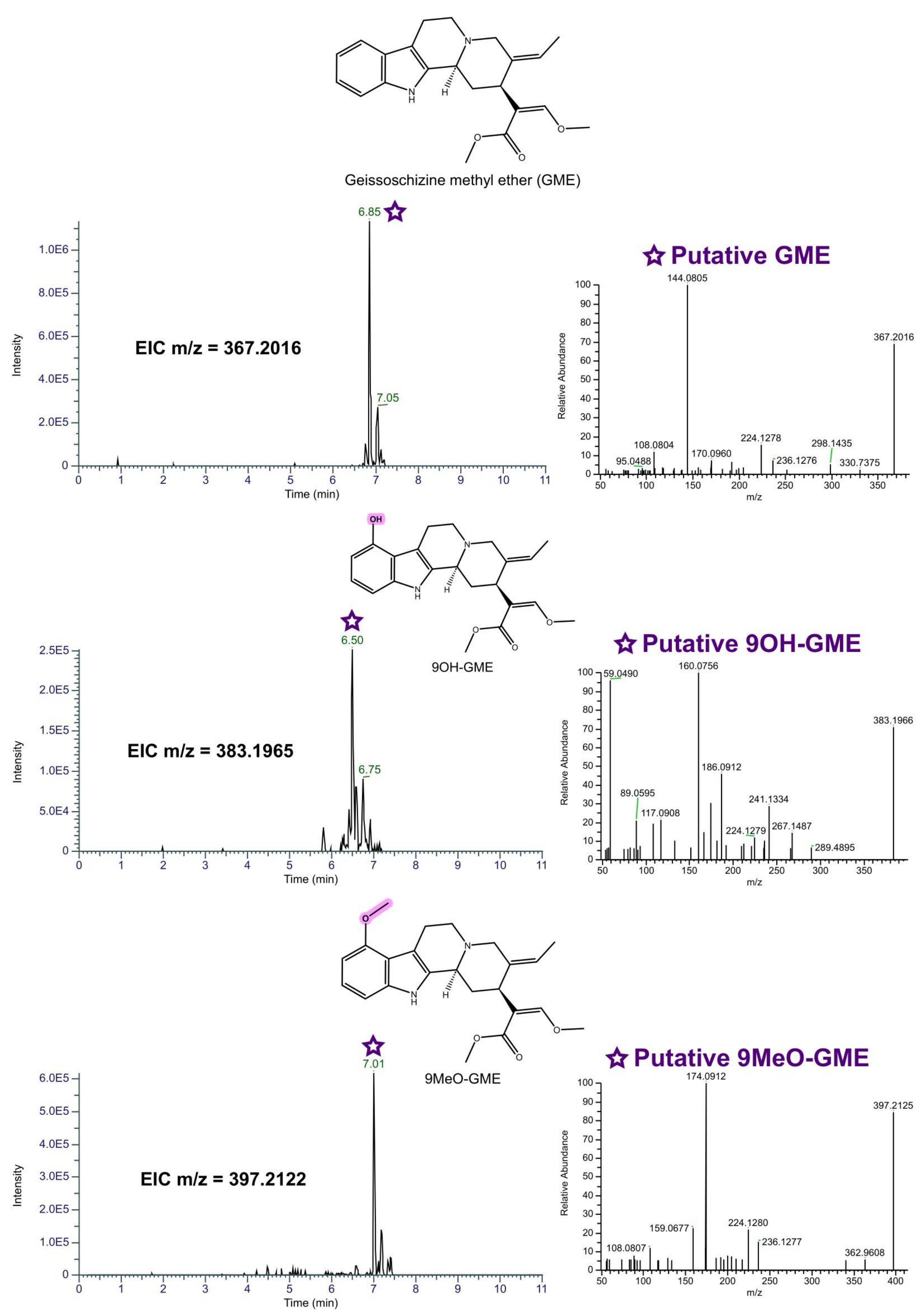

Geissoschizine methyl ether (GME)
EIC m/z = 367.2016
6.85
7.05
Intensity
Time (min)
Putative GME
144.0805
367.2016
108.0804
95.0488
170.0960
224.1278
236.1276
298.1435
330.7375
Relative Abundance
m/z
OH
9OH-GME
EIC m/z = 383.1965
6.50
6.75
Putative 9OH-GME
59.0490
160.0756
383.1966
186.0912
89.0595
117.0908
241.1334
224.1279
267.1487
289.4895
9MeO-GME
EIC m/z = 397.2122
7.01
Putative 9MeO-GME
174.0912
397.2125
159.0677
224.1280
108.0807
236.1277
362.9608

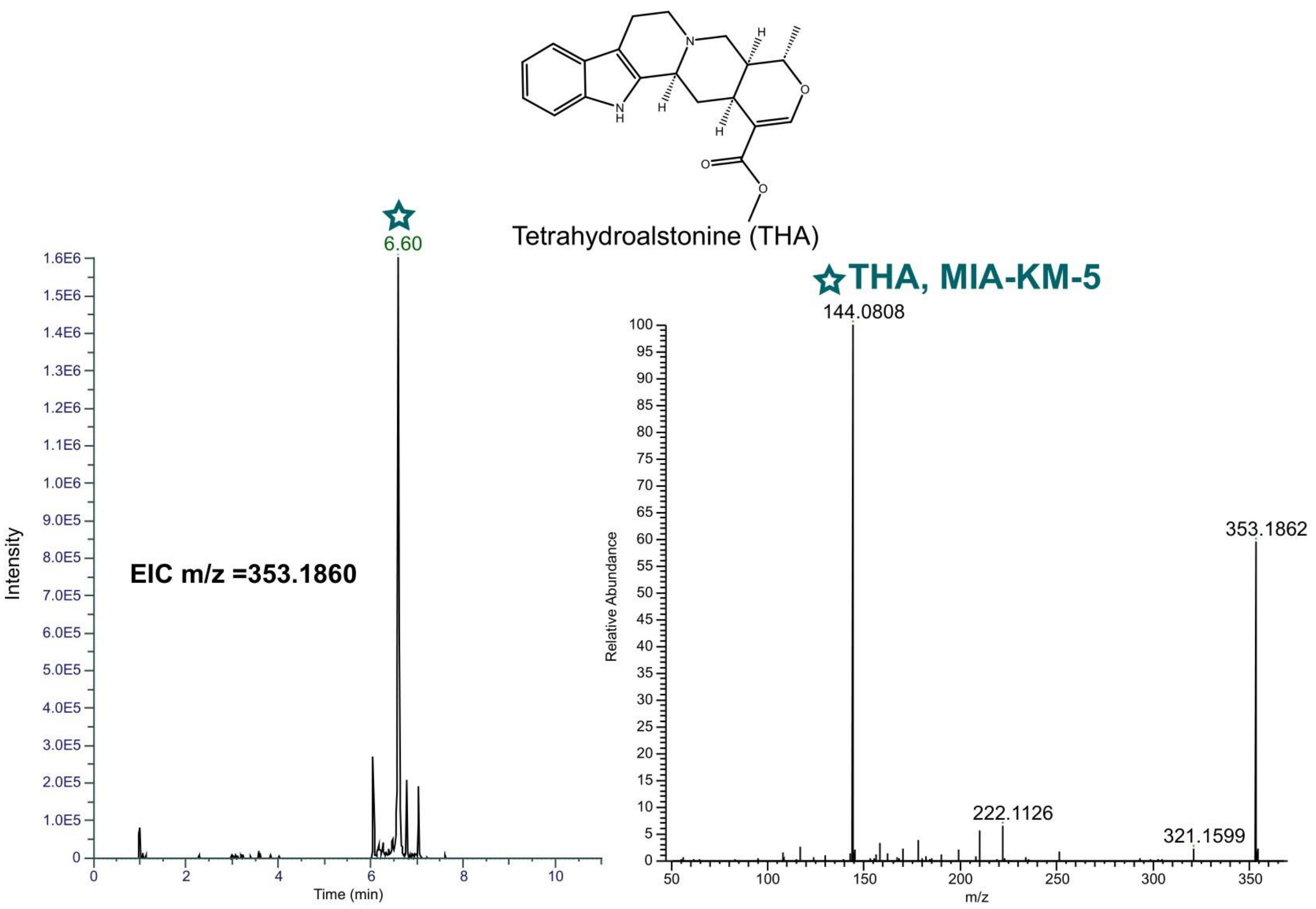

N
NH
H
H
H
O
O
O
Tetrahydroalstonine (THA)
6.60
THA, MIA-KM-5
EIC m/z =353.1860
Intensity
Time (min)
144.0808
353.1862
222.1126
321.1599
Relative Abundance
m/z

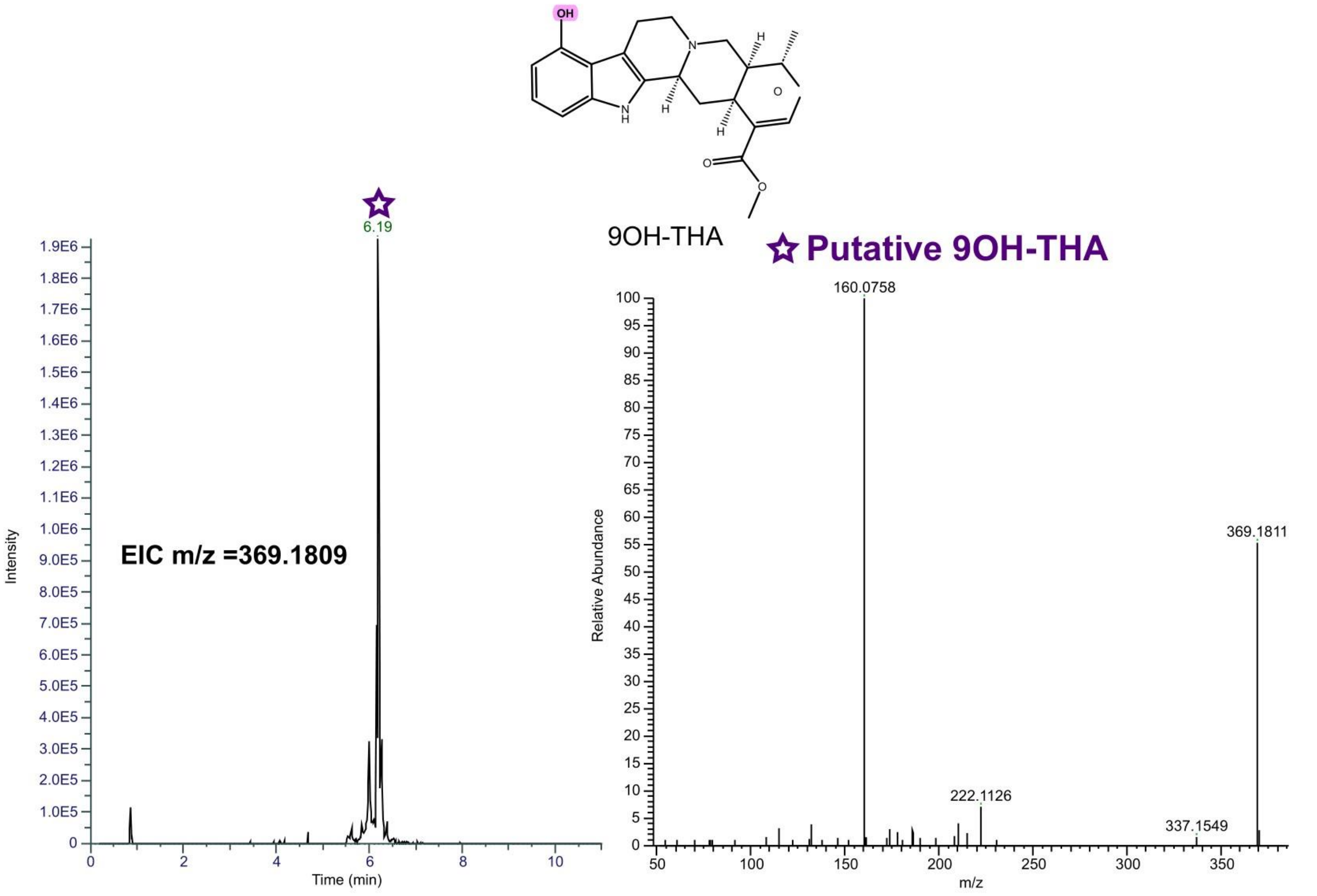

OH
N
NH
H
H
H
O
O
O
9OH-THA
6.19
Putative 9OH-THA
EIC m/z =369.1809
Intensity
Time (min)
160.0758
369.1811
222.1126
337.1549
Relative Abundance
m/z

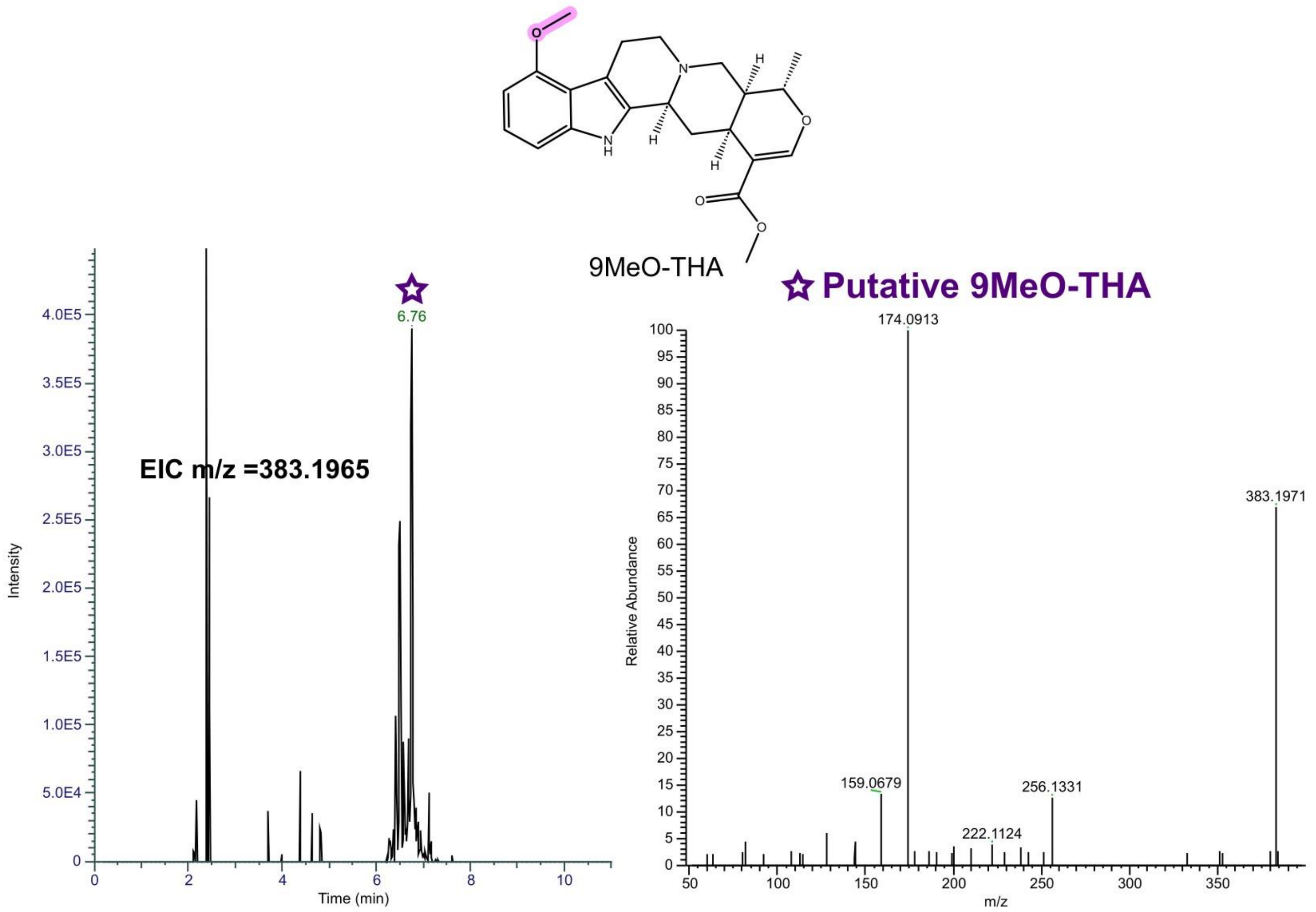

**Supplementary Figure 10.** DCS-derived shunt products identified in kratom yeast strains using untargeted metabolomics data processing pipeline. **A)** Strictosidine deglucosylation by SGD leads to multiple strictosidine aglycone isoforms in chemical equilibrium that get stabilized through reduction by medium-chain alcohol dehydrogenases (ADHs) leading to diverse MIA scaffolds. **B)** EIC and MS2 spectra for the different MIAs identified including yohimbane-type MIAs, heteroyohimbane-type MIAs and geissoschizine methyl ether as well as hydroxylated and metoxylated analogs resulting from incorporation of 4OH-tryptamine in the pathway and subsequent O-methylation by Hp9OMT. Blue color indicates the product has been identified using an authentic analytical standard matching RT, absolute mass and MS2 spectra. Purple color indicates a putative identification using exact mass, MS2 spectra computation for structure prediction with SIRIUS (Dührkop et al., 2019) and knowledge on MIA pathways. The star symbol indicates from which peak MS2 had been acquired.

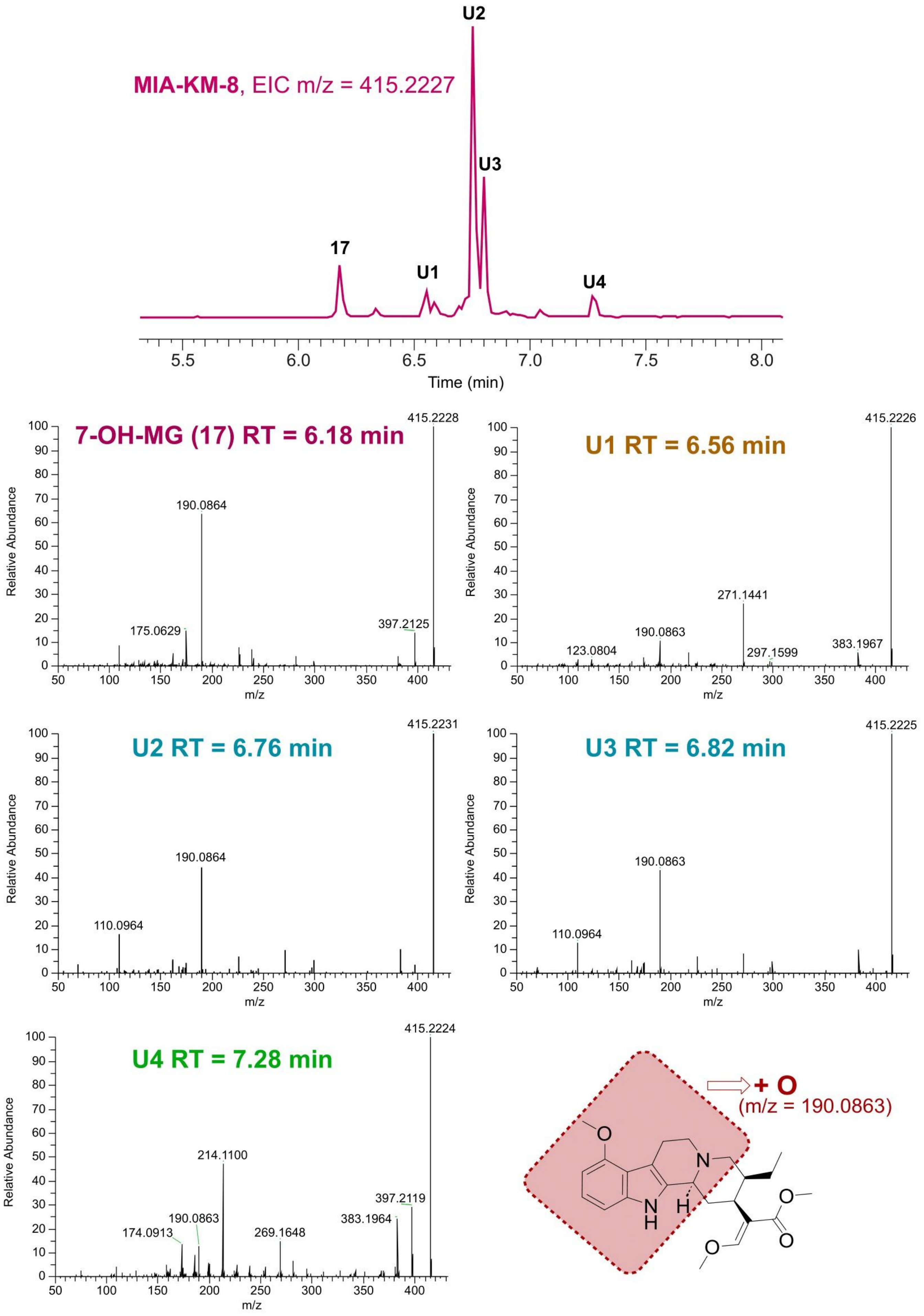

MIA-KM-8, EIC m/z = 415.2227
17
U1
U2
U3
U4
Time (min)
7-OH-MG (17) RT = 6.18 min
415.2228
190.0864
175.0629
397.2125
U1 RT = 6.56 min
415.2226
271.1441
190.0863
123.0804
297.1599
383.1967
U2 RT = 6.76 min
415.2231
190.0864
110.0964
U3 RT = 6.82 min
415.2225
190.0863
110.0964
U4 RT = 7.28 min
415.2224
214.1100
397.2119
383.1964
190.0863
174.0913
269.1648
Relative Abundance
m/z
+ O
(m/z = 190.0863)

**Supplementary Figure 11.** MS/MS fragmentation of HsCYP3A4-generated oxidation products (U = unknown). Fragmentation of mono-oxidated indoloquinolizine moiety of mitragynine (in red) leads to identifying fragment m/z=190.0863 found in all unknown products.

#### Supplementary Tables

**Supplementary table 1. Codon optimized gene sequences cloned in this study** (start codons are highlighted in bold; stop codons are underlined).

| Name | Organism | Reference | Nucleotide sequence |
|---|---|---|---|
| RseSGD | *Rauvolfia serpentina* | (Zhang et al., 2022) | **ATG**GACAACACTCAAGCTGAACCTTTAGTTGTTGCTATCGTTCCAAAGCCAAACGCCAGTACCGAACACACCAATTCTCATTTGATTCCAGTCACCAGATCCAAGATTGTCGTTCACAGACGTGACTTCCCGCAAGATTTCATTTTCGGTGCCGGTGGTTCCGCTTACCAATGTGAAGGCGCTTACAACGAAGGTAACAGAGGTCCATCTATCTGGGACACTTTCACCCAAAGATCTCCAGCCAAGATTTCTGACGGTTCTAATGGTAACCAAGCTATCAACTGTTACCACATGTACAAAGAAGACATCAAAATCATGAAGCAAACTGGTCTTGAGTCCTACAGATTCTCCATCTCTTGGTCCAGAGTTTTACCTGGTGGTAGATTGGCCGCTGGTGTTAACAAGGACGGTGTTAAGTTCTACCATGATTTTATCGATGAACTATTGGCAAATGGTATCAAGCCATCCGTGACTTTGTTCCACTGGGATTTGCCACAAGCTTTGGAAGATGAATATGGTGGTTTTTTATCCCACAGAATCGTCGATGATTTCTGTGAATATGCCGAATTCTGTTTCTGGGAATTCGGTGACAAGATCAAGTACTGGACTACCTTCAACGAACCACACACTTTTGCCGTCAACGGTTACGCTTTAGGAGAATTTGCTCCAGGTAGAGGTGGTAAGGGTGACGAAGGTGACCCAGCTATTGAACCATACGTTGTCACTCACAACATTTTGTTGGCTCACAAGGCTGCTGTCGAAGAATACAGAAACAAGTTCCAAAAGTGCCAGGAAGGTGAAATTGGTATTGTCTTAAACTCCATGTGGATGGAACCATTGTCCGATGTTCAAGCTGACATTGACGCTCAAAAGAGAGCTTTGGATTTCATGTTGGGTTGGTTCTTGGAACCATTGACCACCGGTGACTACCCAAAGAGCATGAGAGAATTGGTTAAAGGTAGACTACCAAAGTTTTCTGCTGATGACTCTGAAAAGTTGAAAGGTTGTTACGACTTCATCGGTATGAACTACTACACTGCTACTTACGTTACCAACGCTGTTAAATCCAACTCTGAAAAGTTGTCTTACGAAACTGACGACCAAGTTACTAAGACCTTCGAAAGAAACCAAAAGCCAATTGGTCATGCCTTGTACGGTGGTTGGCAACACGTCGTCCCATGGGGTTTATACAAATTGTTGGTCTACACCAAGGAAACCTACCATGTACCAGTTTTGTATGTCACTGAATCTGGTATGGTTGAAGAAAACAAGACCAAGATCTTGTTGTCAGAAGCTAGAAGAGATGCTGAAAGAACTGATTATCACCAAAAGCACTTGGCTTCTGTCCGTGATGCCATCGATGACGGTGTTAACGTTAAGGGTTACTTCGTCTGGTCTTTCTTCGACAACTTTGAATGGAACTTGGGCTACATCTGTAGATACGGTATCATTCACGTTGACTACAAGTCTTTCGAACGTTACCCAAAGGAATCTGCTATCTGGTACAAGAACTTCATTGCTGGTAAATCTACCACATCCCCAGCTAAGAGAAGACGTGAAGAAGCCCAAGTTGAATTGGTCAAGAGACAAAAGACTTGA |
| MsDCS1 | *Mitragyna speciosa* | (Schotte et al., 2023) | **ATG**GCTGGTAAGTGTGCACAAGAAGAACATACTGTTAAGGCCTTCGGTTGGGCTGCTCGTGAAGCTTCTGGAGCCCTATCTCCATACGGTTTCTCCAGAAGAGCTACTGGTGAAAGAGATGTTAGAGTTAAGATCTTGTACTGTGGTATCTGTAGAACTGACGCTGAAATGATCTCTGATAAATTCTGTTTCACCAAGTACCCACATGTTCCAGGTCACGAAATTGTCGGTGTCGTTTCTGAGGTTGGTAACAAAGTCCAAAAGTTCAAGGTTGGTGCTAAAGTTGGTGTCACCGGTATCATTGGTTGTTGTCGTACCTGTTACTCTTGCACCAACGGTCTGGAATCCTACTGTCCAAACGTTGCCTTGACTGAAGCCGGTGAAGGTGGTTGCTCCAATTTTATCGTCCTAGACGAAGATTTCGTATTCAGATGGCCAGAAAAGTTGCCATTAGACTTGGGTGCTCCATTATTGTGTGCCGGTGCTGCTTCTTACTCCCCACTGAAGAACTTCGGTTTGGACAAGCCAGGCTTGCACATTGGTATTGCTGGCTTAGGTGGTATGGGTCACGTTGCTGTCAAGTTTGCTAAGGCTTTCGGGGCCAAGGTCACTGTCATCTCTACATCTGACAACAAGAAGGAAGAAGCTATCAAGAAATATGGTGCTGATGCTTTCTTGAACTCATCCAACCCAGAACAAATGAGAGCCGCCGCTGGTACCTTGGCTGCTATTGTTGATACTATTCCATCCCCTCACTCTTTGGTTCCATTGTTGGATTTGTTGTTGCCACACGGTAAGGTCATTGTTTTGGGTGCCCCTTCTGAACCTTTTGTCTTGCCAGTTATGCCATTGTTGCAAGGTGGTAGAGTCGTTGCTGGTAGTTCCGGTGCCAGCTTGAAGCAAATCCAAGAAATGTTGGACTTCGCTGCTGAACACAACATTGTCGCTGACGCTGAAGTTATACCAATTGACTACATCAACACCGCTATCAAGAGAATCGAAAAGGGTGACATTAAGTACAGATTCGTGGTCGATATCGGTAACACTTTAAAGTCTGCTTGA |

| | | | |
|---|---|---|---|
| MsDCS2 | *Mitragyna speciosa* | (Schotte et al., 2023) | **ATG**GCTGAAAAGTCTCCAGAAGAAGAACATCCAGTTAAGGCTTTCGGTTTG<br>GCTGCCAAGGACTCTTCTGGTATTTTGTCCCCTTTCAACTTCTCCAGAAGAG<br>CTACTGGTGACCACGATGTTCAATTGCGTGTTTTGTACTGTGGTTTGTGTTA<br>CTACGACACCAAGATGATCAAGAACAAGAGAGGTGTCACCAGATACCCATT<br>TGTTTTCGGTCACGAAATTGTCGGTGAAGTTACAGAAATCGGTAGAGAAGT<br>CCAAAAGTTCAAGGTTGGTGACAAGGTCGGTGTTGGTTGTATGGTCGCTTC<br>TTGCCGTTCATGCGAATCTTGTGCAAACAACTGTGAAAACTACTGTCCAAAC<br>GTTTCTGTTACTGATGGTGCCTTCTTCTTCAAGACCGGTGAAGTCTTATACG<br>GTGGCTGTTCTGATATCATGGTTGCCGATGAAAATTTTGTCATCAGATGGCC<br>AGAGAATTTCCCATTGGATGCTGGTGCCCCTCTATTATGTGCCGGTATAAC<br>CACTTACTCCCCATTGAGAAACTTTGGTCTGGATAAACCAGGTATTCACGTG<br>GGTATCTACGGTCTAGGTGGCTTGGGTCATGTTGCTGTCCAATTCGCTAAG<br>GCTTTCGGTGCTAAAGTCACTGTAATCTCTAGCTCCGACAGAAAGAGAATG<br>GAAGCTATTGAAAAATTGGGTGCTGACTCCTTCTTGGTCAACTCCAACTTGG<br>AAGAAATGCAAGCTGCTATGGGTACCATGCACGGTATCATCGATACCGTTC<br>CAGCTAACCACTCTTTGGTCCCATTGTTGGACTTGTTGAAGCCACAAGGTAA<br>GTTGATTGTTGTCGGTGGTCCAGAAAAGCCATTCGAATTGCCAGTCTTCCC<br>ATTATTGCAAGGTGGTAGATTGGTTGCTGGTTCTGCTACCGGTGGGATCAA<br>GCAAACTCAAGAAATGATTGACTTTGCCGCTGAACACAACATCTTGCCACAC<br>GTTGAAGTTGTTTCCGTTGACTACGTTAACACTGCTATCGAAAGAACTGAAC<br>GTGGTGATGTCAAGTACAGATTCGTCATTGACATTGGTAACACTTTATATTG<br>A |
| CpDCS | *Cinchona pubescens* | (Schotte et al., 2023) | **ATG**ATGGCTGGTAAGTCTCAAGAAGATGGTCAAACAGTTAAAGCTTTGGGT<br>TGGGCTGCTCGTGAAGTTTCTGGTGCAATCTCTCCATTCGATTTCAGTAGAA<br>GAGCTCCAGGTGAAAGAGATGTTCAAGTTAAGATCTTGTACTGTGGTATCT<br>GTTCTTTCGACACTGAAATGATTAACAACAAGTTCGGTTTCACCAGATACCC<br>ATTTGTTTTGGGTCACGAGATTGTCGGTGTTGTCTCCGAAGTCGGTAGAAA<br>GGTCCAAAAGTTCAAGATTGGTGACAAAGTCGGTGTTGGTACCATGATTGG<br>TTCATGTAGAACCTGTTACTCTTGTACTCATAACTTGGAAAACTACTGTCCAA<br>AGGTTACTTTGACTGAAGCCACCTCCGGTGGTTGCTCCAATTTGGTGATTG<br>CTGATGAAGACTTCGTTTTCCACTGGCCAGTTAACCTGCCATTGGACTTAG<br>GAGCTCCATTGTTGTGTGCTGGCATCACTGTCTACTCTCCATTAAAGAACTT<br>CGAATTGGACAAGCCTGGTTTGAGAATCGGTGTCGTTGGCTTAGGTGGTAT<br>TGGTCATATCGCCGTCAAATTTGCTAAGGCTTTCGGTGCCAAGGTCACCGT<br>CATCTCCTCTTCTGAATCCAAGAAGGTCGAAGCTATTGAAAAGTACGGTGCT<br>GACTCTTTCTTGGTCAGTTCTGACCCAGGTCAAATGTTGGCTGCTGCCGGT<br>ACTTTGGATGGTGTAATTGATACCGTTCCAGCTCCACACTCTATCCTACCTT<br>TCTTAGACTTGTTGTTACCAAGAGGTAAGTTGATCATTTTGGGTGCCCCAAT<br>GGAACCATTCGTCTTGCCAATTTATCCATTGTTGCAAGGTGGTCGTGTTGTT<br>GCTGGTTCTGCTACTGGTGGTTTGAAGCAAATCCAAGAAATGTTGCACTTC<br>GCTGCTGAACACAACATCGTTGCCGACGGAGAAGTTATTCCAATCGACGAT<br>ATCAACACTGCTATCAAGAGAATTGAAAAGGGTGATGTCAAGTACAGATTTG<br>TTGTTGACATTGGTAACACCTTGAAATCCGCCGGTTCCAGAGACTTGGGTT<br>GA |
| PcPsiH | *Psylocibe cubensis* | (Milne et al., 2020) | **ATG**ATCGCTGTTTTGTTCTCTTTCGTTATCGCTGGTTGTATCTACTACATCGT<br>TTCTAGAAGAGTTAGAAGATCTAGATTGCCACCAGGTCCACCAGGTATCCC<br>AATCCCATTCATCGGTAACATGTTCGACATGCCAGAAGAATCTCCATGGTTG<br>ACTTTCTTGCAATGGGGTAGAGACTACAACACTGACATCTTGTACGTTGACG<br>CTGGTGGTACTGAAATGGTTATCTTGAACACTTTGGAAACTATCACTGACTT<br>GTTGGAAAAGAGAGGTTCTATCTACTCTGGTAGATTGGAATCTACTATGGTT<br>AACGAATTGATGGGTTGGGAATTCGACTTGGGTTTCATCACTTACGGTGAC<br>AGATGGAGAGAAGAAAGAAGAATGTTCGCTAAGGAATTCTCTGAAAAGGGT<br>ATCAAGCAATTCAGACACGCTCAAGTTAAGGCTGCTCACCAATTGGTTCAAC<br>AATTGACTAAGACTCCAGACAGATGGGCTCAACACATCAGACACCAAATCG<br>CTGCTATGTCTTTGGACATCGGTTACGGTATCGACTTGGCTGAAGACGACC<br>CATGGTTGGAAGCTACTCACTTGGCTAACGAAGGTTTGGCTATCGCTTCTG<br>TTCCAGGTAAGTTCTGGGTTGACTCTTTCCCATCTTTGAAGTACTTGCCAGC<br>TTGGTTCCCAGGTGCTGTTTTCAAGAGAAAGGCTAAGGTTTGGAGAGAAGC<br>TGCTGACCACATGGTTGACATGCCATACGAAACTATGAGAAAGTTGGCTCC<br>ACAAGGTTTGACTAGACCATCTTACGCTTCTGCTAGATTGCAAGCTATGGAC |

| | | | |
|---|---|---|---|
| | | | TTGAACGGTGACTTGGAACACCAAGAACACGTTATCAAGAACACTGCTGCT<br>GAAGTTAACGTTGGTGGTGGTGACACTACTGTTTCTGCTATGTCTGCTTTCA<br>TCTTGGCTATGGTTAAGTACCCAGAAGTTCAAAGAAAGGTTCAAGCTGAATT<br>GGACGCTTTGACTAACAACGGTCAAATCCCAGACTACGACGAAGAAGACGA<br>CTCTTTGCCATACTTGACTGCTTGTATCAAGGAATTGTTCAGATGGAACCAA<br>ATCGCTCCATTGGCTATCCCACACAAGTTGATGAAGGACGACGTTTACAGA<br>GGTTACTTGATCCCAAAGAACACTTTGGTTTTCGCTAACACTTGGGCTGTTT<br>TGAACGACCCAGAAGTTTACCCAGACCCATCTGTTTTCAGACCAGAAAGAT<br>ACTTGGGTCCAGACGGTAAGCCAGACAACACTGTTAGAGACCCAAGAAAG<br>GCTGCTTTCGGTTACGGTAGAAGAAACTGTCCAGGTATCCACTTGGCTCAA<br>TCTACTGTTTGGATCGCTGGTGCTACTTTGTTGTCTGCTTTCAACATCGAAA<br>GACCAGTTGACCAAAACGGTAAGCCAATCGACATCCCAGCTGACTTCACTA<br>CTGGTTTCTTCAGACACCCAGTTCCATTCCAATGTAGATTCGTTCCAAGAAC<br>TGAACAAGTTTCTCAATCTGTTTCTGGTCCATAA |
| PcCPR | *Psylocibe cubensis* | (Milne et al., 2020) | **ATG**GCTTCTTCTTCTTCTGACGTTTTCGTTTTGGGTTTGGGTGTTGTTTTGG<br>CTGCTTTGTACATCTTCAGAGACCAATTGTTCGCTGCTTCTAAGCCAAAGGT<br>TGCTCCAGTTTCTACTACTAAGCCAGCTAACGGTTCTGCTAACCCAAGAGA<br>CTTCATCGCTAAGATGAAGCAAGGTAAGAAGAGAATCGTTATCTTCTACGGT<br>TCTCAAACTGGTACTGCTGAAGAATACGCTATCAGATTGGCTAAGGAAGCT<br>AAGCAAAAGTTCGGTTTGGCTTCTTTGGTTTGTGACCCAGAAGAATACGACT<br>TCGAAAAGTTGGACCAATTGCCAGAAGACTCTATCGCTTTCTTCGTTGTTGC<br>TACTTACGGTGAAGGTGAACCAACTGACAACGCTGTTCAATTGTTGCAAAAC<br>TTGCAAGACGAATCTTTCGAATTCTCTTCTGGTGAAAGAAAGTTGTCTGGTT<br>TGAAGTACGTTGTTTTCGGTTTGGGTAACAAGACTTACGAACACTACAACTT<br>GATCGGTAGAACTGTTGACGCTCAATTGGCTAAGATGGGTGCTATCAGAAT<br>CGGTGAAAGAGGTGAAGGTGACGACGACAAGTCTATGGAAGAAGACTACTT<br>GGAATGGAAGGACGGTATGTGGGAAGCTTTCGCTACTGCTATGGGTGTTGA<br>AGAAGGTCAAGGTGGTGACTCTGCTGACTTCGTTGTTTCTGAATTGGAATCT<br>CACCCACCAGAAAAGGTTTACCAAGGTGAATTCTCTGCTAGAGCTTTGACTA<br>AGACTAAGGGTATCCACGACGCTAAGAACCCATTCGCTGCTCCAATCGCTG<br>TTGCTAGAGAATTGTTCCAATCTGTTGTTGACAGAAACTGTGTTCACGTTGA<br>ATTCAACATCGAAGGTTCTGGTATCACTTACCAACACGGTGACCACGTTGG<br>TTTGTGGCCATTGAACCCAGACGTTGAAGTTGAAAGATTGTTGTGTGTTTTG<br>GGTTTGGCTGAAAAGAGAGACGCTGTTATCTCTATCGAATCTTTGGACCCA<br>GCTTTGGCTAAGGTTCCATTCCCAGTTCCAACTACTTACGGTGCTGTTTTGA<br>GACACTACATCGACATCTCTGCTGTTGCTGGTAGACAAATCTTGGGTACTTT<br>GTCTAAGTTCGCTCCAACTCCAGAAGCTGAAGCTTTCTTGAGAAACTTGAAC<br>ACTAACAAGGAAGAATACCACAACGTTGTTGCTAACGGTTGTTTGAAGTTGG<br>GTGAAATCTTGCAAATCGCTACTGGTAACGACATCACTGTTCCACCAACTAC<br>TGCTAACACTACTAAGTGGCCAATCCCATTCGACATCATCGTTTCTGCTATC<br>CCAAGATTGCAACCAAGATACTACTCTATCTCTTCTTCTCCAAAGATCCACC<br>CAAACACTATCCACGCTACTGTTGTTGTTTTGAAGTACGAAAACGTTCCAAC<br>TGAACCAATCCCAAGAAAGTGGGTTTACGGTGTTGGTTCTAACTTCTTGTTG<br>AACTTGAAGTACGCTGTTAACAAGGAACCAGTTCCATACATCACTCAAAACG<br>GTGAACAAAGAGTTGGTGTTCCAGAATACTTGATCGCTGGTCCAAGAGGTT<br>CTTACAAGACTGAATCTTTCTACAAGGCTCCAATCCACGTTAGAAGATCTAC<br>TTTCAGATTGCCAACTAACCCAAAGTCTCCAGTTATCATGATCGGTCCAGGT<br>ACTGGTGTTGCTCCATTCAGAGGTTTCGTTCAAGAAAGAGTTGCTTTGGCTA<br>GAAGATCTATCGAAAAGAACGGTCCAGACTCTTTGGCTGACTGGGGTAGAA<br>TCTCTTTGTTCTACGGTTGTAGAAGATCTGACGAAGACTTCTTGTACAAGGA<br>CGAATGGCCACAATACGAAGCTGAATTGAAGGGTAAGTTCAAGTTGCACTG<br>TGCTTTCTCTAGACAAAACTACAAGCCAGACGGTTCTAAGATCTACGTTCAA<br>GACTTGATCTGGGAAGACAGAGAACACATCGCTGACGCTATCTTGAACGGT<br>AAGGGTTACGTTTACATCTGTGGTGAAGCTAAGTCTATGTCTAAGCAAGTTG<br>AAGAAGTTTTGGCTAAGATCTTGGGTGAAGCTAAGGGTGGTTCTGGTCCAG<br>TTGAAGGTGTTGCTGAAGTTAAGTTGTTGAAGGAAAGATCTAGATTGATGTT<br>GGACGTTTGGTCTTAA |
| EnolMT | *Mitragyna speciosa* | (Schotte et al., 2023) | **ATG**CAACCTCAAAGAGGTAGAAAGAGAGAAAGAGAAAGAGACAGAGAGGA<br>AATGGAATCTGTCCAATCTAACTCTTCTTCCTCCGACCAATTTGCTATGAAG<br>GGTGGTGATGATGACTTCTCCTACACCAAGAACTCCACCTGGCAACGTGAT |

| | | | |
|---|---|---|---|
| | | | GCTATCCAAGCCACAAAGTTCTTTATCCAAGAAAGTATCGCTGAAAAATTAG<br>ATGTTAACAAGTTCTGTGGTAAGGCTTTCTGTGTTGCCGATTTGGGCTGTTC<br>CGTTGGTCCAAACACTCTAATTGCCATGCAAAACATCGTTGAAGCTGTCGAA<br>TTGAAATTCAAGAACAGAAAGGGTTTCCACTCTCCAACCATTCCAGAATTCC<br>AAGTTTTCTTCAACGATCATACTGTTAATGATTTCAACACTTTGTTCCGTTCC<br>CTGCCAACTGGTCACGACAAGAGATACTACGGTGTCGGTGTTCCAGGTTCC<br>TTCTACGGGAGATTATTCCCATGCGATTCCATCCACATCATGCACACCTCTT<br>TTTCTACTCCATTCTTGTCTCAAGTCCCAAAGGAAGTCATTGATAAGAACTC<br>GGCTGCTTGGAACAAAGGTAGAATCCATCACAATTACGCTAAAGCTGATGT<br>CTTGAAGGCTTACGAAGCTCAACACGCCGAAGACATCGACTGTTTCTTGAC<br>TGCTAGAGCTAAGGAATTGGTCCACGGTGGTTTGTTGATGGATGTTACCTC<br>TTTCAGACCAGATGGTGTCCCTCATACTCACGTTTTGACCAACATTGGTATG<br>GAAGTTTTGGGTTACTGTTTGATGGACTTGGCTGGTTTGATTGACGAAGAAA<br>ACGTCGACTCTTACAACGTTCCAGTTTATTTGCAATCCCCAGAAGAATTGAA<br>GCAAGCTGTCCAAAGAAACAAGTACTTCTCTATTGAAAAGATGGAATCCGTG<br>CCAATGATGATTGACTCTGACGTATCTGCTAAGGCCCAACAATACTCATTGG<br>GTATGAGAGCCGTTATGGGTGACGTTATCAGAGAACAATTCGGTGCTGAAA<br>TTGTCGACAAATTGTTTGACTTATTTAAGAAGAAGTTGGAAGAACACCCAAA<br>CTTCGCTAAGGGTGTTGTTTTAGACATGTTCGTCTTGCTAAAGCGTAACGCT<br>GAAGACTGA |
| Hp9OMT | *Hamelia patens* | (Kim et al., 2023) | **ATG**TACGTCGTCATCTCCCACAGATACATCTGTCACCCATCCAAGGTTTCTT<br>TGTCTTCTATGTTCTCTAACCAAACTCATCCTTTGGCTATGGATGATGCCCA<br>ACAAAATTATTTAAGAGTTTTGGAAATGGGTTGTACCCAAATTTTGCACGCT<br>GTTTTCACCACTGTTATTGAATTGAACGTCTTTGAGATCATTGCCAAGGCCG<br>GTCCAGAAGCTCAATTGTCTGCTGCTGAAATTTCCTCTCACTTGCCAACTCA<br>AAACCAACAAGCTCCAGCTATCTTGGAAAGAATGTTGCAATTGCTCGCTTCT<br>TACTCTGTCTTGAAGTGTGTTCACGTTTCCAACCAAGATGGTAGAGGTACCA<br>GATTGTACGGTTTGACTCCAATGTGTCGTTACTTGGTTGCTGACACCATGG<br>GTATCTCCATGGGTCCAGCCATGTTGTGTTACACCGACAAGGCTATGGCTG<br>ATTCCTGGTCTTATTTGAAGGACGCCGTGTTAGAAGGTAAGATTCCATTCAA<br>CAAAGCTAACAAGATGGACTTGTTCGAATACTTCGGTAAGTCTAGTACTTTG<br>AACGAAACTTTCAACCAAGCTATGCACTCTGAAACCTTTTTCGTCTTGAGAG<br>CTGTCTTGCAAAACTACAAGGGTTTCGAAGCTTTGAAAGAATTGGTCGATGT<br>TGGTGGTGGTTTGGGTATCACTTTAAACGCTATTATTTCTAAGTACCCAGGT<br>ATCAGAGGTATCAACTTCGATTTGCCACAAGTTATCAAGGATGCCCCAATCA<br>GAACCGGTGTTGAAAATTTACCAGGTGACATGTTTGAATACGTTCCAAAGG<br>GTGAAGCGATTCTATTAAAAAACATTTTGCACGACTGGACTGACGAACATTG<br>TCTTAAATTGTTGAAGAACTGTTACAACGCCCTGCCTGAACACGGTAAGGTC<br>ATTGTCATCGAAATGATTTTGCCAACCTCTCCAGAAAATGACTTGTTATCCA<br>GAGCAGTCTTCTTCGTTGACATCATGATGTTAGCTTTGACATCTGGCGGAA<br>GAGAACGTACTTTGAAGGAATTCGACGCTCTGGCTAAGGGTGCTGGTTTCA<br>TTGCTTGTAAGTTAGTTTGCCAAACCTTCGGTTACGGTATCTTGGAATTCTA<br>CAAGTCATCCACTTTGAACAGCTCCGAAATCCCATGA |
| HsCYP3A4 | *Homo sapiens* | (Cheng et al., 2006) | **ATG**GCTTTGATCCCAGACTTAGCCATGGAAACGTGGTTATTGTTGGCTGTTT<br>CCTTGGTTTTGCTATATTTGTACGGTACCCACTCTCACGGTTTGTTCAAGAA<br>GTTGGGTATTCCAGGTCCAACCCCATTGCCATTCTTGGGTAACATTCTTTCT<br>TACCATAAGGGTTTCTGTATGTTCGACATGGAATGTCACAAGAAATATGGTA<br>AGGTCTGGGGTTTCTACGATGGCCAACAACCAGTGCTCGCCATCACTGACC<br>CTGATATGATCAAGACCGTTTTGGTTAAGGAATGTTACTCTGTTTTCACTAA<br>CAGAAGACCGTTCGGTCCAGTTGGTTTCATGAAGTCTGCTATTTCCATTGCT<br>GAAGATGAAGAATGGAAGAGATTGAGATCTCTGTTGTCTCCAACTTTCACTT<br>CCGGTAAGTTAAAGGAAATGGTTCCAATTATCGCCCAATACGGTGACGTCT<br>TGGTCAGAAACTTGCGTAGAGAAGCTGAAACTGGTAAGCCAGTTACCTTGA<br>AAGATGTTTTCGGTGCTTACTCCATGGACGTCATTACTTCTACCTCTTTCGG<br>TGTTAACATCGACTCTTTGAACAACCCACAAGATCCTTTCGTCGAAAATACC<br>AAGAAGCTATTGAGATTCGATTTCTTGGATCCATTCTTCTTGTCGATCACCG<br>TCTTTCCATTTTTGATTCCAATCTTGGAAGTCTTAAATATCTGTGTCTTTCCA<br>AGAGAAGTCACTAACTTCTTGAGAAAGTCCGTCAAAAGAATGAAGGAATCA<br>CGTTTGGAAGACACCCAAAAGCACAGAGTTGACTTCTTGCAATTGATGATTG<br>ATTCTCAAAACTCTAAGGAAACAGAATCCCATAAAGCTTTGTCTGACTTGGA |

| | | | |
|---|---|---|---|
| | | | ATTGGTTGCTCAATCTATCATCTTTATCTTCGCTGGTTACGAAACCACCTCCTCCGTTTTGTCCTTCATCATGTACGAATTGGCTACCCACCCAGATGTCCAACAAAAGTTGCAAGAAGAAATTGATGCTGTCCTACCAAACAAGGCTCCACCAACTTACGACACTGTTTTACAAATGGAATACTTGGACATGGTTGTCAACGAAACTTTGAGATTATTCCCAATCGCTATGAGATTGGAACGTGTCTGCAAGAAGGATGTTGAAATCAACGGTATGTTCATTCCAAAGGGTGTCGTTGTTATGATTCCATCCTACGCTTTGCACCGTGACCCAAAGTACTGGACTGAACCAGAAAAGTTCTTGCCAGAAAGATTCTCCAAGAAGAACAAGGACAACATCGACCCATACATCTACACTCCATTTGGTTCTGGTCCAAGAAACTGTATCGGTATGAGATTTGCTTTGATGAACATGAAATTGGCCTTGATTAGAGTTTTACAAAATTTCTCTTTCAAGCCATGTAAGGAGACTCAAATTCCTTTGAAATTAAGCTTAGGTGGTTTACTACAACCAGAAAAGCCAGTCGTTCTGAAGGTTGAATCTAGAGATGGTACTGTCAGTGGTGCCTGA |
| HsCPR | *Homo sapiens* | (Cheng et al., 2006) | **ATG**GGTGACTCTCACGTCGATACCTCCTCCACTGTCTCTGAAGCTGTTGCTGAAGAAGTCTCCTTATTCTCCATGACTGACATGATCTTATTTTCCCTGATTGTGGGCTTGTTGACTTACTGGTTCTTGTTCAGAAAGAAGAAGGAAGAAGTCCCAGAATTCACCAAGATCCAAACCTTGACATCTTCTGTCAGAGAATCCTCTTTCGTCGAAAAGATGAAGAAGACCGGTAGAAACATCATCGTCTTCTATGGTTCTCAAACTGGTACTGCTGAAGAATTTGCTAACAGATTATCAAAGGATGCTCACAGATACGGTATGAGAGGTATGTCTGCTGACCCAGAAGAATACGATCTAGCTGATTTGTCCTCCTTACCAGAAATTGACAACGCCCTCGTCGTCTTCTGTATGGCCACCTATGGGGAAGGTGATCCAACCGATAATGCTCAAGATTTTTACGACTGGTTGCAAGAAACTGATGTTGACTTGTCTGGTGTCAAGTTCGCCGTTTTCGGTTTGGGTAACAAGACCTACGAACATTTCAACGCTATGGGTAAGTACGTTGACAAGAGATTGGAGCAATTAGGTGCCCAAAGAATCTTCGAACTCGGTTTGGGTGATGACGATGGTAACTTGGAAGAAGATTTCATTACCTGGAGAGAACAATTCTGGCCAGCTGTCTGTGAACACTTCGGTGTCGAAGCTACCGGTGAAGAGAGCTCTATCCGTCAATACGAACTGGTAGTTCACACTGATATTGATGCCGCCAAGGTATACATGGGTGAAATGGGTAGACTTAAATCCTACGAAAACCAAAAGCCACCTTTCGACGCTAAGAACCCATTTTTGGCTGCTGTAACCACTAATAGAAAATTGAACCAAGGTACTGAAAGACATTTGATGCACTTGGAATTGGACATCTCTGACTCCAAGATCAGATACGAAAGCGGTGACCACGTTGCCGTCTACCCAGCCAACGACTCTGCTTTAGTTAACCAGTTGGGTAAGATTTTGGGCGCGGATTTGGATGTCGTTATGTCTTTGAACAACTTGGACGAAGAATCCAACAAGAAACATCCATTCCCATGTCCAACTTCTTACAGAACTGCTTTGACTTACTACTTGGACATTACTAACCCTCCTCGTACAAATGTTTTGTACGAACTAGCTCAATACGCTTCCGAACCATCCGAACAAGAATTGCTGAGAAAGATGGCTTCTAGTTCTGGTGAAGGTAAGGAATTATATTTGTCTTGGGTTGTTGAAGCCAGAAGACACATCTTGGCCATTCTTCAAGACTGTCCATCTTTAAGACCACCAATTGACCACTTATGCGAATTGTTGCCAAGATTGCAAGCTAGATATTACTCTATTGCTTCATCCTCCAAGGTCCACCCAAACTCTGTTCACATCTGTGCCGTTGTTGTCGAATACGAAACTAAAGCTGGTCGTATCAACAAGGGTGTTGCTACCAACTGGCTACGTGCAAAGGAACCAGCAGGTGAAAACGGTGGTCGTGCTTTGGTTCCAATGTTCGTTAGAAAGTCTCAATTCAGATTGCCATTCAAGGCTACTACTCCAGTTATCATGGTTGGTCCAGGTACCGGTGTTGCTCCATTCATTGGTTTCATCCAAGAAAGAGCTTGGTTGAGACAACAAGGTAAAGAAGTTGGGGAAACCTTGTTGTACTACGGTTGTAGAAGATCTGATGAAGACTACTTGTACCGTGAAGAATTGGCTCAATTCCACAGAGATGGTGCTTTGACCCAATTGAATGTCGCTTTTTCTCGTGAACAATCCCACAAGGTTTACGTCCAACACTTGTTGAAGCAAGACAGAGAACATTTGTGGAAGTTGATTGAAGGTGGTGCTCATATCTACGTTTGTGGTGACGCCAGAAACATGGCTAGAGATGTTCAAAACACTTTCTACGATATTGTTGCTGAATTGGGTGCCATGGAACACGCTCAAGCTGTCGACTACATCAAGAAATTGATGACCAAGGGTAGATACTCATTGGACGTCTGGTCCTGA |

**Supplementary table 2. Plasmids cloned and used in this study**

| Name | Description | Reference |
|---|---|---|
| pCfB9336 | pgRNA_II-1_NatMX | (Babaei et al., 2021) |
| pCfB9337 | pgRNA_IV-1_NatMX | (Babaei et al., 2021) |

| | | |
|---|---|---|
| pCfB9340 | pgRNA_VIII-1_NatMX | (Babaei et al., 2021) |
| pCfB9342 | pgRNA_XIII-1_NatMX | (Babaei et al., 2021) |
| pCfB9355 | II-1_Markerfree_BackBone | (Babaei et al., 2021) |
| pCfB9356 | IV-1_Markerfree_BackBone | (Babaei et al., 2021) |
| pCfB9359 | VIII-1_Markerfree_BackBone | (Babaei et al., 2021) |
| pCfB9361 | XIII-1_Markerfree_BackBone | (Babaei et al., 2021) |
| MoClo Yeast Toolkit (YTK) | https://www.addgene.org/kits/moclo-ytk/ | (Lee et al., 2015) |
| pCfB9073 | XI-1 integration plasmid for PcCPR expressed from pTEF1 and PcPsiH expressed from pPGK1 | (Milne et al., 2020) |
| pMHO10 | VIII-1 integration plasmid for PcCPR expressed from pTEF1 and PcPsiH expressed from pPGK1 | This study. |
| pMC-X4 | preassembled yeast MoClo level-2 backbone with GFP dropout for genomic integration in site X-4 | (Otto et al., 2021) |
| pMHGG2_Vec_II-1 | preassembled yeast MoClo level-2 backbone with GFP dropout for genomic integration in site II-1 | This study. |
| pMHGG2_Vec_IV-1 | preassembled yeast MoClo level-2 backbone with GFP dropout for genomic integration in site IV-1 | This study. |
| pMHGG2_Vec_XIII-1 | preassembled yeast MoClo level-2 backbone with GFP dropout for genomic integration in site XIII-1 | This study. |
| pMHGG0_3_EnolMT | Level-0 yeast MoClo plasmid containing EnolMT CDS part. | This study. |
| pMHGG0_3_CpDCS | Level-0 yeast MoClo plasmid containing CpDCS CDS part. | This study. |
| pMHGG0_3_MsDCS1 | Level-0 yeast MoClo plasmid containing MsDCS1 CDS part. | This study. |
| pMHGG0_3_MsDCS2 | Level-0 yeast MoClo plasmid containing EnolMT CDS part. | This study. |
| pMHGG0_3_RsSGD | Level-0 yeast MoClo plasmid containing MsDCS2 CDS part. | This study. |
| pMHGG0_Hp9OMT | Level-0 yeast MoClo plasmid containing Hp9OMT CDS part. | This study. |
| pMHGG0_3_HsCYP3A4 | Level-0 yeast MoClo plasmid containing HsCYP3A4 CDS part. | This study. |
| pMHGG0_3_HsCPR | Level-0 yeast MoClo plasmid containing HsCPR CDS part. | This study. |
| pMHGG1_Vec_Pos1 | Level-1 yeast MoClo vector plasmid containing GFP dropout. Includes 2µ yeast ORI and URA yeast selection marker. Gene position number 1 in level-2 plasmid. | This study. |
| pMHGG1_Vec_Pos2 | Level-1 yeast MoClo vector plasmid containing GFP dropout. Includes 2µ yeast ORI and URA yeast selection marker. Gene position number 2 in level-2 plasmid. | This study. |
| pMHGG1_Vec_Pos3* | Level-1 yeast MoClo vector plasmid containing GFP dropout. Includes 2µ yeast ORI and URA yeast selection marker. Gene position number 3 and final gene in level-2 plasmid. | This study. |
| pMHGG1_Vec_Pos2* | Level-1 yeast MoClo vector plasmid containing GFP dropout. Includes 2µ yeast ORI and URA yeast selection marker. Gene position number 2 and final gene in level-2 plasmid. | This study. |
| pMHGG1_Vec_Pos1* | Level-1 yeast MoClo vector plasmid containing GFP dropout. Includes 2µ yeast ORI and URA yeast selection marker. Gene position number 1 and final gene in level-2 plasmid. | This study. |
| pMHGG1_EnolMT_Pos1 | Level-1 yeast MoClo plasmid containing EnolMT transcription unit. Includes 2µ yeast ORI and URA yeast selection marker. | This study. |

| | | |
|---|---|---|
| pMHGG1_RsSGD_Pos2 | Level-1 yeast MoClo plasmid containing RsSGD transcription unit. Includes 2μ yeast ORI and URA yeast selection marker. | This study. |
| pMHGG1_MsDCS1_Pos3 | Level-1 yeast MoClo plasmid containing MsDCS1 transcription unit. Includes 2μ yeast ORI and URA yeast selection marker. | This study. |
| pMHGG1_MsDCS2_Pos3 | Level-1 yeast MoClo plasmid containing MsDCS2 transcription unit. Includes 2μ yeast ORI and URA yeast selection marker. | This study. |
| pMHGG1_CpDCS_Pos3 | Level-1 yeast MoClo plasmid containing CpDCS transcription unit. Includes 2μ yeast ORI and URA yeast selection marker. | This study. |
| pMHGG1_Hp9OMT_Pos1* | Level-1 yeast MoClo plasmid containing Hp9OMT transcription unit. Includes 2μ yeast ORI and URA yeast selection marker. | This study. |
| pMHGG1_HsCYP3A4_Pos1 | Level-1 yeast MoClo plasmid containing HsCYP3A4 transcription unit. Includes 2μ yeast ORI and URA yeast selection marker. | This study. |
| pMHGG1_HsCPR_Pos2* | Level-1 yeast MoClo plasmid containing HsCPR transcription unit. Includes 2μ yeast ORI and URA yeast selection marker. | This study. |
| pMHGG2_MsDCS1 | Level-2 multigene vector for integration at site II-1. Contains transcription units for RsSGD, MsDCS1 and EnolMT. | This study. |
| pMHGG2_MsDCS2 | Level-2 multigene vector for integration at site II-1. Contains transcription units for RsSGD, MsDCS2 and EnolMT. | This study. |
| pMHGG2_CpDCS | Level-2 multigene vector for integration at site II-1. Contains transcription units for RsSGD, CpDCS and EnolMT. | This study. |
| pMHGG2_Hp9OMT | Level-2 multigene vector for integration at site XIII-1. Contains transcription unit for Hp9OMT. | This study. |
| pMHGG2_HsCYP3A4 | Level-2 multigene vector for integration at site IV-1. Contains transcription units for HsCYP3A4 and HsCPR. | This study. |

#### Supplementary table 3. Strains used in this study

| Name | Parent | Genotype | Reference |
|---|---|---|---|
| MIA-B0 | CEN.PK2-1C | MATa; his3D1; leu2-3_112; ura3-52; trp1-289; pTEF1-SpCas9-tCYC1, | (Zhang et al., 2022) |
| MIA-CM-3 | MIA-B0 | MATa, his3D1, leu2-3_112, ura3-52, trp1-289, atf1Δ oye2Δ adh6Δ oye3Δ ari1Δ, PTEF1-SpyCas9-TCYC1, PTEF1-CroCPR-TPRM9, PPGK1-CroCYB5-TIDP1, PMLS1-AgrGPPS2-TVPS13, PFBA1-GgaFPSN144W-TIDP1, PPGK1-IDI1-TPRM9, PTDH3-tHMG1-TADH1, PICL1-ERG20F96W, N127WtCroGES-TCYC1, PPGK1-CroTDC-TPRM9, PTDH3-CroG8H-TADH1, PPGK1-Vmi8HGO-A -TADH1,PFBA1-NcaISY-TCYC1, PTEF1-NcaMLPLA-TADH1, | (Zhang et al., 2022) |

| | | | |
|---|---|---|---|
| | | PTEF2-CroIO-TCYC1, PFBA1-CroADH2-TCPS1,PPGK1-Cro7DLGT-TVPS13, PTEF2-Cro7DLH-TCYC1, PTDH3-CroLAMT-TADH1, PTPI1-CroSLS-TIDP1,PFBA1-CroSTR-TPRM9 | |
| MIA-CM-5 | MIA-B0 | MATa, his3D1, leu2-3_112, ura3-52, trp1-289, atf1Δ oye2Δ adh6Δ oye3Δ ari1Δ, PTEF1-SpyCas9-TCYC1, PTEF1-CroCPR-TPRM9, PPGK1-CroCYB5-TIDP1, **PTEF2-AgrGPPS2-TVPS13**, PFBA1-GgaFPSN144W-TIDP1, PPGK1-IDI1-TPRM9, PTDH3-tHMG1-TADH1, **PCCW12-ERG20F96W**, N127WtCroGES-TCYC1, PPGK1-CroTDC-TPRM9, PTDH3-CroG8H-TADH1, PPGK1-Vmi8HGO-A -TADH1,PFBA1-NcaISY-TCYC1, PTEF1-NcaMLPLA-TADH1, PTEF2-CroIO-TCYC1, PFBA1-CroADH2-TCPS1,PPGK1-Cro7DLGT-TVPS13, PTEF2-Cro7DLH-TCYC1, PTDH3-CroLAMT-TADH1, PTPI1-CroSLS-TIDP1,PFBA1-CroSTR-TPRM9. | (Zhang et al., 2022) |
| MIA-CZ-1 | MIA-CM-5 | MATa, his3D1, leu2-3_112, ura3-52, trp1-289, atf1Δ oye2Δ adh6Δ oye3Δ ari1Δ, **rox1Δ**, PTEF1-SpyCas9-TCYC1, PTEF1-CroCPR-TPRM9, PPGK1-CroCYB5-TIDP1, PTEF2-AgrGPPS2-TVPS13, PFBA1-GgaFPSN144W-TIDP1, PPGK1-IDI1-TPRM9, PTDH3-tHMG1-TADH1, PCCW12-ERG20F96W, N127WtCroGES-TCYC1, PPGK1-CroTDC-TPRM9, PTDH3-CroG8H-TADH1, PPGK1-Vmi8HGO-A -TADH1,PFBA1-NcaISY-TCYC1, PTEF1-NcaMLPLA-TADH1, PTEF2-CroIO-TCYC1, PFBA1-CroADH2-TCPS1,PPGK1-Cro7DLGT-TVPS13, PTEF2-Cro7DLH-TCYC1, PTDH3-CroLAMT-TADH1, PTPI1-CroSLS-TIDP1,PFBA1-CroSTR-TPRM9, **PCCW12-ZWF1-TADH1, PPGK1-INO2-TCYC1.** | This study. |
| MIA-KM-1 | MIA-CZ-1 | MATa, his3D1, leu2-3_112, ura3-52, trp1-289, atf1Δ oye2Δ adh6Δ oye3Δ ari1Δ, rox1Δ, PTEF1-SpyCas9-TCYC1, PTEF1-CroCPR-TPRM9, PPGK1-CroCYB5-TIDP1, PTEF2-AgrGPPS2-TVPS13, PFBA1-GgaFPSN144W-TIDP1, PPGK1-IDI1-TPRM9, PTDH3-tHMG1-TADH1, PCCW12-ERG20F96W, N127WtCroGES-TCYC1, PPGK1-CroTDC-TPRM9, PTDH3-CroG8H-TADH1, PPGK1-Vmi8HGO-A -TADH1,PFBA1-NcaISY-TCYC1, PTEF1-NcaMLPLA-TADH1, PTEF2-CroIO-TCYC1, PFBA1-CroADH2-TCPS1,PPGK1-Cro7DLGT-TVPS13, PTEF2-Cro7DLH-TCYC1, PTDH3-CroLAMT-TADH1, PTPI1-CroSLS-TIDP1,PFBA1-CroSTR-TPRM9, PCCW12-ZWF1-TADH1, PPGK1-INO2-TCYC1, **PTEF1-PcCPR-TADH1, PPGK1-PcPsiH-TCYC1**. | This study. |
| MIA-KM-2 | MIA-KM-1 | MATa, his3D1, leu2-3_112, ura3-52, trp1-289, atf1Δ oye2Δ adh6Δ oye3Δ ari1Δ, rox1Δ, PTEF1-SpyCas9-TCYC1, PTEF1-CroCPR-TPRM9, PPGK1-CroCYB5-TIDP1, PTEF2-AgrGPPS2-TVPS13, PFBA1-GgaFPSN144W-TIDP1, PPGK1-IDI1-TPRM9, PTDH3-tHMG1-TADH1, PCCW12-ERG20F96W, N127WtCroGES-TCYC1, PPGK1-CroTDC-TPRM9, PTDH3-CroG8H-TADH1, PPGK1-Vmi8HGO-A -TADH1,PFBA1-NcaISY-TCYC1, PTEF1-NcaMLPLA-TADH1, PTEF2-CroIO-TCYC1, PFBA1-CroADH2-TCPS1,PPGK1-Cro7DLGT-TVPS13, PTEF2-Cro7DLH-TCYC1, PTDH3-CroLAMT-TADH1, PTPI1-CroSLS-TIDP1,PFBA1-CroSTR-TPRM9, PCCW12-ZWF1-TADH1, PPGK1-INO2-TCYC1, PTEF1-PcCPR-TADH1, PPGK1-PcPsiH-TCYC1, **PCCW12-RseSGD-TPGK1, PHHF2-EnolMT-TTDH1, PTDH3-MsDCS1-TENO2.** | This study. |
| MIA-KM-3 | MIA-KM-1 | MATa, his3D1, leu2-3_112, ura3-52, trp1-289, atf1Δ oye2Δ adh6Δ oye3Δ ari1Δ, rox1Δ, PTEF1-SpyCas9-TCYC1, PTEF1-CroCPR-TPRM9, PPGK1-CroCYB5-TIDP1, PTEF2-AgrGPPS2-TVPS13, PFBA1-GgaFPSN144W-TIDP1, PPGK1-IDI1-TPRM9, PTDH3-tHMG1-TADH1, PCCW12-ERG20F96W, N127WtCroGES-TCYC1, PPGK1-CroTDC-TPRM9, PTDH3-CroG8H-TADH1, PPGK1-Vmi8HGO-A -TADH1,PFBA1-NcaISY-TCYC1, PTEF1-NcaMLPLA-TADH1, PTEF2-CroIO-TCYC1, PFBA1-CroADH2-TCPS1,PPGK1-Cro7DLGT-TVPS13, PTEF2-Cro7DLH-TCYC1, PTDH3-CroLAMT-TADH1, PTPI1-CroSLS-TIDP1,PFBA1-CroSTR-TPRM9, PCCW12-ZWF1-TADH1, PPGK1-INO2-TCYC1, PTEF1-PcCPR-TADH1, | This study. |

| | | | |
|---|---|---|---|
| | | PPGK1-PcPsiH-TCYC1, **PCCW12-RseSGD-TPGK1, PHHF2-EnolMT-TTDH1, PTDH3-MsDCS2-TENO2.** | |
| MIA-KM-4 | MIA-KM-1 | MATa, his3D1, leu2-3_112, ura3-52, trp1-289, atf1Δ oye2Δ adh6Δ oye3Δ ari1Δ, rox1Δ, PTEF1-SpyCas9-TCYC1, PTEF1-CroCPR-TPRM9, PPGK1-CroCYB5-TIDP1, PTEF2-AgrGPPS2-TVPS13, PFBA1-GgaFPSN144W-TIDP1, PPGK1-IDI1-TPRM9, PTDH3-tHMG1-TADH1, PCCW12-ERG20F96W, N127WtCroGES-TCYC1, PPGK1-CroTDC-TPRM9, PTDH3-CroG8H-TADH1, PPGK1-Vmi8HGO-A -TADH1,PFBA1-NcaISY-TCYC1, PTEF1-NcaMLPLA-TADH1, PTEF2-CroIO-TCYC1, PFBA1-CroADH2-TCPS1,PPGK1-Cro7DLGT-TVPS13, PTEF2-Cro7DLH-TCYC1, PTDH3-CroLAMT-TADH1, PTPI1-CroSLS-TIDP1,PFBA1-CroSTR-TPRM9, PCCW12-ZWF1-TADH1, PPGK1-INO2-TCYC1, PTEF1-PcCPR-TADH1, PPGK1-PcPsiH-TCYC1, **PCCW12-RseSGD-TPGK1, PHHF2-EnolMT-TTDH1, PTDH3-CpDCS-TENO2.** | This study. |
| MIA-KM-5 | MIA-KM-2 | MATa, his3D1, leu2-3_112, ura3-52, trp1-289, atf1Δ oye2Δ adh6Δ oye3Δ ari1Δ, rox1Δ, PTEF1-SpyCas9-TCYC1, PTEF1-CroCPR-TPRM9, PPGK1-CroCYB5-TIDP1, PTEF2-AgrGPPS2-TVPS13, PFBA1-GgaFPSN144W-TIDP1, PPGK1-IDI1-TPRM9, PTDH3-tHMG1-TADH1, PCCW12-ERG20F96W, N127WtCroGES-TCYC1, PPGK1-CroTDC-TPRM9, PTDH3-CroG8H-TADH1, PPGK1-Vmi8HGO-A -TADH1,PFBA1-NcaISY-TCYC1, PTEF1-NcaMLPLA-TADH1, PTEF2-CroIO-TCYC1, PFBA1-CroADH2-TCPS1,PPGK1-Cro7DLGT-TVPS13, PTEF2-Cro7DLH-TCYC1, PTDH3-CroLAMT-TADH1, PTPI1-CroSLS-TIDP1,PFBA1-CroSTR-TPRM9, PCCW12-ZWF1-TADH1, PPGK1-INO2-TCYC1, PTEF1-PcCPR-TADH1, PPGK1-PcPsiH-TCYC1, PCCW12-RseSGD-TPGK1, PHHF2-EnolMT-TTDH1, PTDH3-MsDCS1-TENO2, **PTEF2-Hp9OMT-TTDH1.** | This study. |
| MIA-KM-6 | MIA-KM-3 | MATa, his3D1, leu2-3_112, ura3-52, trp1-289, atf1Δ oye2Δ adh6Δ oye3Δ ari1Δ, rox1Δ, PTEF1-SpyCas9-TCYC1, PTEF1-CroCPR-TPRM9, PPGK1-CroCYB5-TIDP1, PTEF2-AgrGPPS2-TVPS13, PFBA1-GgaFPSN144W-TIDP1, PPGK1-IDI1-TPRM9, PTDH3-tHMG1-TADH1, PCCW12-ERG20F96W, N127WtCroGES-TCYC1, PPGK1-CroTDC-TPRM9, PTDH3-CroG8H-TADH1, PPGK1-Vmi8HGO-A -TADH1,PFBA1-NcaISY-TCYC1, PTEF1-NcaMLPLA-TADH1, PTEF2-CroIO-TCYC1, PFBA1-CroADH2-TCPS1,PPGK1-Cro7DLGT-TVPS13, PTEF2-Cro7DLH-TCYC1, PTDH3-CroLAMT-TADH1, PTPI1-CroSLS-TIDP1,PFBA1-CroSTR-TPRM9, PCCW12-ZWF1-TADH1, PPGK1-INO2-TCYC1, PTEF1-PcCPR-TADH1, PPGK1-PcPsiH-TCYC1, PCCW12-RseSGD-TPGK1, PHHF2-EnolMT-TTDH1, PTDH3-MsDCS2-TENO2, **PTEF2-Hp9OMT-TTDH1.** | This study. |
| MIA-KM-7 | MIA-KM-4 | MATa, his3D1, leu2-3_112, ura3-52, trp1-289, atf1Δ oye2Δ adh6Δ oye3Δ ari1Δ, rox1Δ, PTEF1-SpyCas9-TCYC1, PTEF1-CroCPR-TPRM9, PPGK1-CroCYB5-TIDP1, PTEF2-AgrGPPS2-TVPS13, PFBA1-GgaFPSN144W-TIDP1, PPGK1-IDI1-TPRM9, PTDH3-tHMG1-TADH1, PCCW12-ERG20F96W, N127WtCroGES-TCYC1, PPGK1-CroTDC-TPRM9, PTDH3-CroG8H-TADH1, PPGK1-Vmi8HGO-A -TADH1,PFBA1-NcaISY-TCYC1, PTEF1-NcaMLPLA-TADH1, PTEF2-CroIO-TCYC1, PFBA1-CroADH2-TCPS1,PPGK1-Cro7DLGT-TVPS13, PTEF2-Cro7DLH-TCYC1, PTDH3-CroLAMT-TADH1, PTPI1-CroSLS-TIDP1,PFBA1-CroSTR-TPRM9, PCCW12-ZWF1-TADH1, PPGK1-INO2-TCYC1, PTEF1-PcCPR-TADH1, PPGK1-PcPsiH-TCYC1, PCCW12-RseSGD-TPGK1, PHHF2-EnolMT-TTDH1, PTDH3-CpDCS-TENO2, **PTEF2-Hp9OMT-TTDH1.** | This study. |
| MIA-KM-8 | MIA-KM-5 | MATa, his3D1, leu2-3_112, ura3-52, trp1-289, atf1Δ oye2Δ adh6Δ oye3Δ ari1Δ, rox1Δ, PTEF1-SpyCas9-TCYC1, PTEF1-CroCPR-TPRM9, PPGK1-CroCYB5-TIDP1, PTEF2-AgrGPPS2-TVPS13, | This study. |

| | | |
|---|---|---|
| | | PFBA1-GgaFPSN144W-TIDP1, PPGK1-IDI1-TPRM9, PTDH3-tHMG1-TADH1, PCCW12-ERG20F96W, N127WtCroGES-TCYC1, PPGK1-CroTDC-TPRM9, PTDH3-CroG8H-TADH1, PPGK1-Vmi8HGO-A -TADH1,PFBA1-NcaISY-TCYC1, PTEF1-NcaMLPLA-TADH1, PTEF2-CroIO-TCYC1, PFBA1-CroADH2-TCPS1,PPGK1-Cro7DLGT-TVPS13, PTEF2-Cro7DLH-TCYC1, PTDH3-CroLAMT-TADH1, PTPI1-CroSLS-TIDP1,PFBA1-CroSTR-TPRM9, PCCW12-ZWF1-TADH1, PPGK1-INO2-TCYC1, PTEF1-PcCPR-TADH1, PPGK1-PcPsiH-TCYC1, PCCW12-RseSGD-TPGK1, PHHF2-EnolMT-TTDH1, PTDH3-MsDCS1-TENO2, PTEF2-Hp9OMT-TTDH1, **PTEF2-HsCYP4A4-TSSA1, PRPL18B-HsCPR-TTDH1**. |

#### Supplementary table 4. Chemical standards used in this study.

| Chemical | CAS | Supplier |
|---|---|---|
| Mitragynine | 4098-40-2 | Biosynth Ltd (UK) |
| Speciogynine | 4697-67-0 | Cayman Chemical (USA) |
| 20S-corynantheidine | 23407-35-4 | Toronto Research Chemicals (Canada) |
| 4-OH-tryptamine | 55206-11-6 | Toronto Research Chemicals (Canada) |
| Loganic acid | 22255-40-9 | Carl Roth GmbH + Co (Germany) |
| Loganin | 18524-94-2 | Santa Cruz Biotechnology (USA) |
| Secologanin | 19351-63-4 | Sigma-Aldrich |
| Tryptamine | 61-54-1 | Sigma-Aldrich |
| Strictosidine | 20824-29-7 | PHYTOCONSULT (Netherlands) discontinued |
| Tetrahydroalstonine | 6474-90-4 | Chengdu Push Bio-technology Co., Ltd. (China) discontinued |
| Rauwolscine | 131-03-3 | Sigma-Aldrich |
| Yohimbine hydrochloride | 65-19-0 | Sigma-Aldrich |

#### Supplementary table 7. Compounds retention time (RT), m/z and main and fragments in LC-MS/MS analysis.

| Compound | Formula | RT (min) | Theoretical Monoisotopic mass | Theoretical $[M+H]^{+1}$ | Observed $[M+H]^{+1}$ | ppm | MS-MS Identifier Peaks |
|---|---|---|---|---|---|---|---|
| Tryptophan | C11H12N2O2 | 4.8 | 204.0898 | 205.0976 | 205.0971 | -2.4 | 118.06<br>146.05<br>188.07 |
| Tryptamine | C10H12N2 | 5.2 | 160.0995 | 161.1078 | 161.1073 | -3.10 | 144.08 |
| 4OH-Tryptamine | C10H12N2O | 2.5 | 176.0944 | 177.1022 | 177.1021 | -0.56 | 160.08 |
| Secologanin | C17H24O10 | 5.8 | 388.1369 | 389.1447 | 389.1446 | -0.3 | 107.04<br>165.05 |

| | | | | | | | |
|---|---|---|---|---|---|---|---|
| Strictosidine | C27H34N2O9 | 6.4 | 530.2264 | 531.2342 | 531.2343 | 0.19 | 144.08<br>514.20<br>531.23 |
| 9OH-Strictosidine | C27H34N2O10 | 5.8 | 546.2208 | 547.2286 | 547.2287 | 0.18 | 160.08<br>530.20<br>547.23 |
| Corynantheidine | C22H28N2O3 | 6.9 (20S) and 7.0 (20R) | 368.2094 | 369.2172 | 369.2184 | 3.25 | 144.08<br>369.22 |
| 9OH-Corynantheidine | C22H28N2O4 | 6.5 (20S) and 6.6 (20R) | 384.2044 | 385.2122 | 385.2120 | -0.52 | 160.08<br>385.21 |
| Dihydrocorynantheine | C21H26N2O3 | 6.6 (20S) and 6.7 (20R) | 354.1938 | 355.2016 | 355.2016 | 0.00 | 144.08<br>355.20 |
| 9OH-Dihydrocorynantheine | C21H26N2O4 | 6.0 (20S) and 6.1 (20R) | 370.1887 | 371.1965 | 371.1965 | 0.00 | 160.08<br>371.20 |
| Mitragynine | C23H30N2O4 | 7.03 | 398.2200 | 399.2278 | 399.2279 | 0.25 | 399.23<br>174.09 |
| 7OH-mitragynine | C23H30N2O5 | 6.18 | 414.2149 | 415.2227 | 415.2228 | 0.24 | 415.22<br>190.09 |
| Speciogynine | C23H30N2O4 | 7.13 | 398.2200 | 399.2278 | 399.2280 | 0.50 | 399.23<br>174.09 |
| Tetrahydroalstonine | C21H24N2O3 | 5.5 | 352.1787 | 353.1865 | 353.1864 | -0.28 | 144.08 |
| Rauwolscine | C21H26N2O3 | 5.4 | 345.1938 | 355.2016 | 355.2023 | 1.97 | 144.08 |
| Geissoschizine methyl ether | C22H26N2O3 | 6.8 | 366.1938 | 367.2016 | 367.2016 | -0.05 | 367.20<br>144.08 |

#### Supplementary Material References

Babaei, M., Sartori, L., Karpukhin, A., Abashkin, D., Matrosova, E., & Borodina, I. (2021). Expansion of EasyClone-MarkerFree toolkit for Saccharomyces cerevisiae genome with new integration sites. *FEMS Yeast Research*, *21*(4), foab027. https://doi.org/10.1093/femsyr/foab027

Cheng, J., Wan, D., Gu, J., Gong, Y., Yang, S., Hao, D., & Yang, L. (2006). Establishment of a yeast system that stably expresses human cytochrome P450 reductase: Application for the study of drug metabolism of cytochrome P450s in vitro. *Protein Expression and Purification*, *47*(2), 467–476. https://doi.org/10.1016/j.pep.2005.11.022

Dührkop, K., Fleischauer, M., Ludwig, M., Aksenov, A. A., Melnik, A. V., Meusel, M., Dorrestein, P. C., Rousu, J., & Böcker, S. (2019). SIRIUS 4: A rapid tool for turning tandem mass spectra into metabolite structure information. *Nature Methods*, *16*(4), 299–302. https://doi.org/10.1038/s41592-019-0344-8

Kim, K., Shahsavarani, M., Garza-García, J. J. O., Carlisle, J. E., Guo, J., De Luca, V., & Qu, Y. (2023). Biosynthesis of kratom opioids. *New Phytologist*, *240*(2), 757–769. https://doi.org/10.1111/nph.19162

Lee, M. E., DeLoache, W. C., Cervantes, B., & Dueber, J. E. (2015). A Highly Characterized Yeast Toolkit for Modular, Multipart Assembly. *ACS Synthetic Biology*, *4*(9), 975–986. https://doi.org/10.1021/sb500366v

Milne, N., Thomsen, P., Mølgaard Knudsen, N., Rubaszka, P., Kristensen, M., & Borodina, I. (2020). Metabolic engineering of Saccharomyces cerevisiae for the de novo production of psilocybin and related tryptamine derivatives. *Metabolic Engineering*, *60*, 25–36. https://doi.org/10.1016/j.ymben.2019.12.007

Otto, M., Skrekas, C., Gossing, M., Gustafsson, J., Siewers, V., & David, F. (2021). Expansion of the Yeast Modular Cloning Toolkit for CRISPR-Based Applications, Genomic Integrations and Combinatorial Libraries. *ACS Synthetic Biology*, *10*(12), 3461–3474. https://doi.org/10.1021/acssynbio.1c00408

Schotte, C., Jiang, Y., Grzech, D., Dang, T.-T. T., Laforest, L. C., León, F., Mottinelli, M., Nadakuduti, S. S., McCurdy, C. R., & O'Connor, S. E. (2023). Directed Biosynthesis of Mitragynine Stereoisomers. *Journal of the American Chemical Society*, *145*(9), 4957–4963. https://doi.org/10.1021/jacs.2c13644

Wirth, N. T., Funk, J., Donati, S., & Nikel, P. I. (2023). QurvE: User-friendly software for the analysis of biological growth and fluorescence data. *Nature Protocols*, *18*(8), Article 8. https://doi.org/10.1038/s41596-023-00850-7

Zhang, J., Hansen, L. G., Gudich, O., Viehrig, K., Lassen, L. M. M., Schrübbers, L., Adhikari, K. B., Rubaszka, P., Carrasquer-Alvarez, E., Chen, L., D'Ambrosio, V., Lehka, B., Haidar, A. K., Nallapareddy, S.,

Giannakou, K., Laloux, M., Arsovska, D., Jørgensen, M. A. K., Chan, L. J. G., … Keasling, J. D. (2022). A microbial supply chain for production of the anti-cancer drug vinblastine. *Nature*, *609*(7926), Article 7926. https://doi.org/10.1038/s41586-022-05157-3
